# How Experience Becomes a Program: Episodic Memory as a Difference Engine

**DOI:** 10.64898/2026.11.16.767088

**Authors:** Yuhang He, Si Wu

## Abstract

A child who has seen a few sums can add numbers of any length: finite experience has become a program, a procedure with memory that holds for inputs of every size. Networks learn procedures by passing errors backward through every step of a computation, over many examples, and what they learn on short inputs seldom holds on long ones. Brains keep something networks discard, the experiences themselves, and we propose that they keep them in order to compare them. When two remembered experiences differ in one place, the first answer at which they differ depended on that place. This method of difference finds the cause of an answer in one step, however long the computation in between, and finds causes that act jointly, each of which alone is invisible to statistics. We show that such comparisons write programs. Comparing an experience with one that differs in a single symbol shows what each answer depends on; in a memory that recalls the past by counting events, each dependence is a linear equation, and the stack, the queue and the counter of computer science are solutions of such equations. Comparing what follows two beginnings shows which states a program must keep, and comparing a question with the question one symbol shorter builds each answer from the one before. Where experience does not decide, nothing is guessed: the learner asks, or says that it does not know. A machine built on these principles, with no back-propagation and no search over programs, writes exact programs for thirteen of the fifteen tasks of a standard benchmark, each in under a minute, from a few hundred short strings and, for two of the tasks, questions of its own; eleven of the programs hold at a million symbols. It learns tasks withheld from its design, builds multiplication from its own addition, and in 5,680 runs announces no false program. The hippocampus has the form of such a difference engine: it stores whole episodes after one exposure, keeps similar ones apart, recalls a neighbour of the present and compares the two. In simulated circuits of neurons the programs run exactly, a replayed sweep finds where two episodes differ, and a local rule learns the gains with no backward pass. On this account, episodic memory is the brain’s means of assigning credit by comparison.

## 1. Introduction

A child who has been shown a few sums adds numbers of any length, and children shown a few nested lists order new ones in the same way at once (Ferrigno et al., 2020). A finite stretch of experience has become a program: a procedure with memory that holds for inputs of every size. How this happens is an old question with two parts. The first is induction. Finitely many examples agree with infinitely many rules: the sums a child has seen agree with addition, and also with a rule that parts from addition on numbers longer than any in those sums; the child arrives at addition (Hume, 1748/2000; Goodman, 1955; Kripke, 1982). The second is credit assignment. An answer is the last step of a long computation, as the last digit of a sum may depend on a carry that began at the first, and the learner has to find which earlier moments it depended on (Minsky, 1961).

Artificial networks answer both parts with one method. Gradient descent moves all the parameters of a network at once so as to reduce its error, and assigns credit by carrying the error backward through every step of the computation (Rumelhart et al., 1986; Werbos, 1990). As an account of how brains learn procedures it meets three difficulties. It needs many examples where a child needs a few (Lake et al., 2017). An error carried back through every step of a computation has no known counterpart in neural circuits (Lillicrap & Santoro, 2019; Lillicrap et al., 2020). And what it finds on short inputs seldom holds on long ones: on fifteen tasks that span the Chomsky hierarchy, LSTMs and Transformers trained on strings of up to 40 symbols fail at greater lengths on most tasks, and networks given a stack or a tape reach further up the hierarchy (Joulin & Mikolov, 2015; Delétang et al., 2023; Ruoss et al., 2023).

Brains keep something that networks in training discard: the experiences themselves (Tulving, 1972). The hippocampus stores an episode after a single exposure and can bring it back later (Scoville & Milner, 1957; Teyler & DiScenna, 1986; Squire, 1992). Keeping whole episodes is costly, and a theory of learning should say what they are for. We propose that they are kept to be compared. Suppose a learner hears the string 10110 answered by its reversal 01101, and remembers hearing 11110 answered by 01111. The two experiences form a *minimal pair*, as *pat* and *bat* do in phonology: the strings differ in one place, the second symbol, and the answers first differ at the fourth. Whatever procedure produced the answers, the fourth answer depended on the second symbol (Figure 1a). Hume stated this rule of difference, and Mill made it one of his canons of experimental inquiry (Hume, 1739–1740; Mill, 1843).

**Figure 1.**
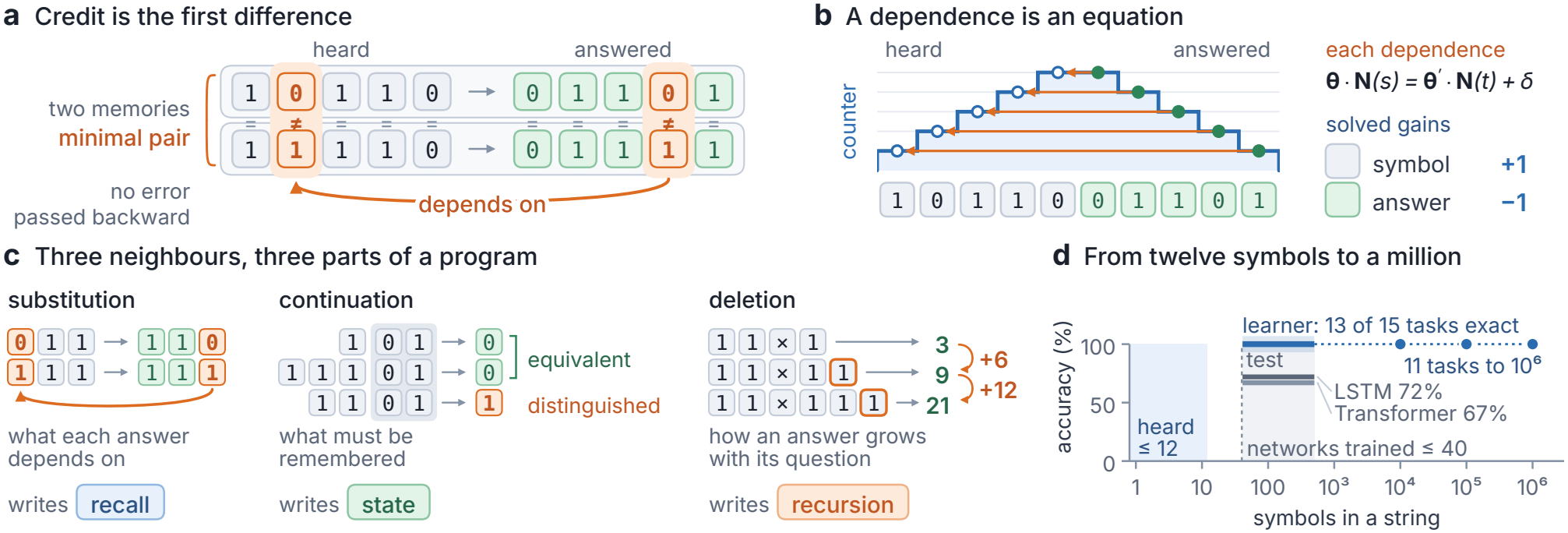
How experience becomes a program. **(a)** A minimal pair: two experiences of reversal differ in the second symbol heard, and the fourth answer, the first to differ, depends on it. **(b)** Dependences are equations in the gains; their solution, one counter turned +1 per symbol and *−*1 per answer, lets each answer recall the record stored at the same reading. **(c)** Three neighbours write three parts of a program: substitution (reversal) writes recalls, continuation (parity) writes states, and deletion (multiplication, lowest digit first, answers in decimal) writes recursion. **(d)** From strings of at most 12 symbols, programs exact up to 500 symbols for thirteen tasks and up to 10^6^ for eleven; grey, LSTMs and Transformers trained on up to 40 symbols (means over the tasks of Table 1).

The rule has two properties that make it a means of assigning credit. It reaches from an answer to its cause in one step, however long the computation between them, because only the two ends are compared: in the reversal of a string of a million symbols, one comparison traces the last answer to the first symbol. And it finds causes that statistics miss. When the causes of an answer act jointly, as the two digits of a column of a binary sum do, each cause alone is uncorrelated with the answer: over all sums, a digit of the answer is 1 as often when the first digit of its column is 0 as when it is 1. A statistic of single experiences finds such causes only by taking them all together, out of all the combinations of earlier moments. One comparison finds each of them.

**Table 1.**
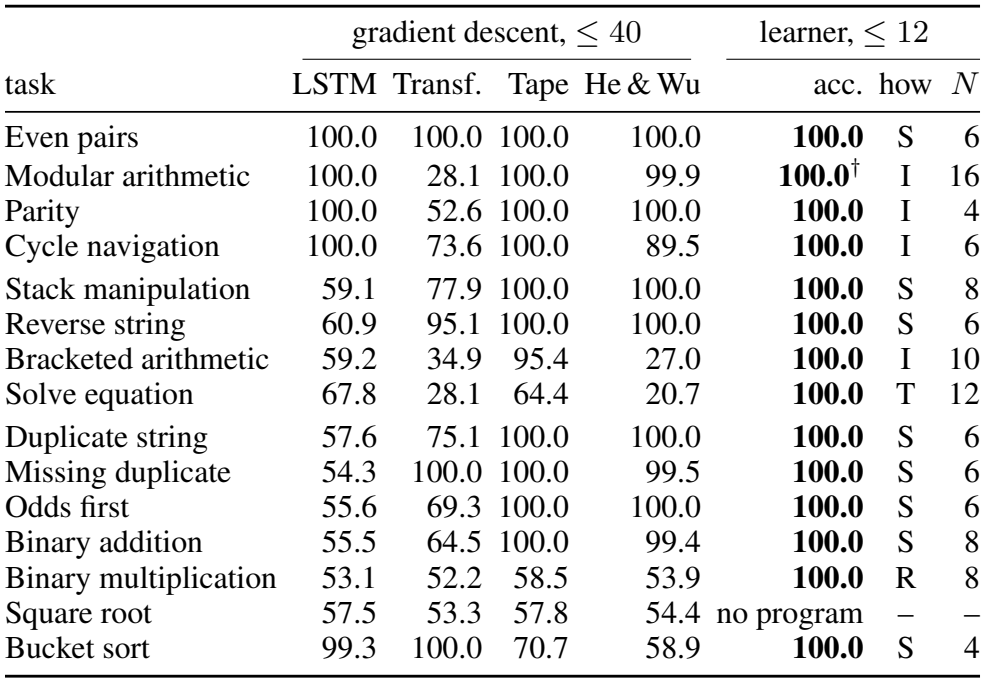
The benchmark of Delétang et al. (2023): accuracy (%) on strings of 41 to 500 symbols; rows: regular, deterministic context-free, context-sensitive tasks. LSTM and Tape-RNN: best of 30 runs; Transformer: best of eleven positional encodings (Ruoss et al., 2023); He & Wu: the first paper’s machine, mean of three seeds. Learner: lowest of ten seeds (bold: every seed exact); how its program runs (S in step, state in the last act; I in step, state in inner speech; T trial; R recursion); smallest *N* at which every seed is exact. ^*†*^From 5,843 strings of up to 16 symbols, with at most one question (Experiment 4).

A gradient is also a comparison: of a computation with a copy of itself changed by an infinitesimal amount, carried back through every step. A difference compares two experiences that actually happened, and needs only that both be kept. Keeping whole episodes is then the price of assigning credit by comparison, and on this view the hippocampus pays it.

We develop this idea into a theory of how experience becomes a program (Section 3), for the memory derived in the first paper of this series (He & Wu, 2026a). That memory stores each moment with the readings of a few counters, integers that each kind of event turns by a fixed amount, and recalls the moment at which one counter stood where another stands now (Section 2). An experience has three kinds of neighbour in memory, and comparing it with each writes one part of a program. A neighbour by substitution, the other half of a minimal pair, shows what each answer depends on. A dependence asks that a counter stand at the answer where another stood at the moment the answer depends on: a linear equation in how far each kind of event turns each counter, and the stack, the queue and the counter of computer science are solutions of such equations (Figure 1b). A neighbour by continuation, another beginning followed by the same string, shows which beginnings a program has to keep in different states, and consolidation in sleep merges those that experience has shown equivalent. A neighbour by deletion, the same question one symbol shorter, shows how an answer grows with its question, and gives recursion (Figure 1c). Nothing is guessed: where experience leaves two possibilities, the learner keeps both, and asks a question or says that it does not know.

For this learner, the format of its memory answers the first part of the old question. A program in this memory never reads how far a counter has gone, only what a recall returns and on which side of zero a counter stands. Among programs of bounded size, one that is right on all inputs up to a length fixed by the bound is therefore right at every length, and a rule that parts from addition beyond that length cannot be written. What can be projected beyond experience is fixed by what memory can express (Sections 3.5 and 6).

A machine built on these principles uses no backpropagation and searches no space of programs (Section 4). It hears a few hundred strings, a median of 457, none longer than 12 symbols. From them it writes exact programs for thirteen of the fifteen tasks above, for each of ten random draws of experience, in a median of 0.5 seconds; for two of the tasks it also asks questions (for multiplication, a median of 990, each the beginning of a string it heard). Of its programs, eleven answer strings of a million symbols (Figure 1d). Gradient descent on the same memory solves at most five of ten of these tasks. The learner also solves all fifteen tasks withheld from its design, builds long multiplication from its own addition, and in 5,680 runs announces no false program, while the same learner made to guess does.

The hippocampal system has the parts this learner needs (Section 5). To compare experiences, a memory has to store each one whole after one exposure, keep similar ones apart, recall a neighbour of the present and compare the two: functions attributed to the dentate gyrus, CA3 and CA1 (Marr, 1971; McClelland et al., 1995; Duncan et al., 2012). In the theory, theta sweeps and replay, in which place cells fire in rapid sequence (Foster & Wilson, 2007), bring two experiences into one cycle of activity; replay often shows an experience other than the present one, including paths never run (Karlsson & Frank, 2009; Gupta et al., 2010). A local rule then solves the equations without a backward pass. It moves each gain by small steps, and the attractor that holds a counter makes a gain near a whole number act as that number, so the counter learned is exact. In simulated circuits of neurons, the learned programs run exactly on strings ten times longer than any heard, a sweep finds where two episodes differ, and a ring learns the counter of the stack from dependences that nobody has sorted (Section 5.5). One requirement, credit by comparison, accounts for findings that several theories of the hippocampus each explain in part: that relational learning and memory for facts and events need the hippocampus and habits do not, that cells fire at a fixed displacement from an object or at a fixed lag after a step of a task, that codes split where futures differ, and that sleep consolidates. The theory also predicts what experiments in animals and people should find (Table 2).

**Table 2.** Predictions, each with the result it follows from and the account it is measured against; Appendix E.4 gives protocols for these and four more. P1 tests the principle most cheaply; P2 tests where the comparison is made, against replay prioritized for value (Mattar & Daw, 2018); P3 separates a comparison at one informative moment from learning that accumulates over trials.

|  |  |  |  |
|---|---|---|---|
| P1 | A rule with joint causes (an exclusive or) is learned in a step once minimal pairs have shown each cause, and not from as many experiences with matched statistics and no pairs. | Th. 1 | associative and Bayesian learning |
| P2 | After a surprise of either sign, sweeps hold the present episode together with its nearest neighbour of the other outcome more often than chance, and CA1 fires where the two differ. | Th. 1 | replay prioritized for value |
| P3 | Silencing CA1 at the first outcome that completes a minimal pair, and not at an equally surprising one that completes none, delays learning; silencing the dentate gyrus during learning spares only rules of one cue. | Th. 1 | learning by prediction error |
| P4 | Vector cells form for an object that the outcome depends on, more than for an equally salient one, at the first informative comparison and before behaviour improves. | Th. 2 | fields formed by exposure |
| P5 | When reward moves from every fourth lap to every fifth, the replayed lap cycle stretches with each miss, while the running lap code and anticipatory licking switch together in one step. | Th. 6 | hidden-state inference |
| P6 | Sleep, more than an equal time awake, merges the codes of situations met in one context with one outcome, and not of situations that only share an outcome. | Th. 3 | schemas built from overlap |
| P7 | After a new start that leads only into the stretch two routines share, the code there alternates between their codes more than on the routines' own laps, and the first choice where they part falls at chance, however unequal their practice. | Th. 3 | latent-cause inference |

This is the third of three papers. The first showed that a memory of the kind the brain keeps for space can run any program, and found its programs by gradient descent (He & Wu, 2026a); the second described how what a circuit knows acts on it as a constraint (He & Wu, 2026b); this one asks where programs come from. Babbage designed two engines: the analytical engine, a universal computer, which has its counterpart in the memory of the first paper, and the difference engine, which computes tables by adding differences (Babbage, 1864). We argue that episodic memory is a difference engine: it turns experience into programs by comparing experiences.

## 2. A Memory That Runs Programs

We use, unchanged, the memory derived in the first paper of this series (He & Wu, 2026a). Take reversal. Hearing abc, the memory stores a, b and c where a counter stands, at 1, 2 and 3; as the answers turn the counter back down, they recall c, b and a.

An experience is a sequence of moments. At each moment a symbol is heard or an answer is due, and the learner acts: it says a symbol, aloud or to itself, or nothing. A *counter c* is an integer that each kind of event (a symbol heard, an answer said, a record returned by memory) turns by a fixed amount, its *gain θ*_*c*_ for that kind; some kinds may reset it to zero. We take a counter to be a module of grid cells, which path integration advances as the animal moves (Hafting et al., 2005; Stensola et al., 2012). Every moment *s* is stored as a *record w*_*s*_ of what was heard and done, with the readings of all counters, as the hippocampal index stores the pattern of a moment (Teyler & DiScenna, 1986). A *recall ρ* has a key counter *k*_*ρ*_, a query counter *q*_*ρ*_ and a filter *γ*_*ρ*_ on the kinds of record it admits. At moment *t* it returns the latest admitted record stored when the key stood where the query stands now. A recall is attention whose keys and queries are counters, compared for equality alone; on this account a place cell makes the comparison, firing when the counters return to its record’s readings (O’Keefe & Dostrovsky, 1971). A *threshold* reports on which side of zero a counter stands. A *state m*, from a finite set, is carried from one act to the next by the acts themselves: by what was last said, aloud or to itself, together with what the recalls returned with it (Section 3.4). The *table* Φ turns all of these into the next act:

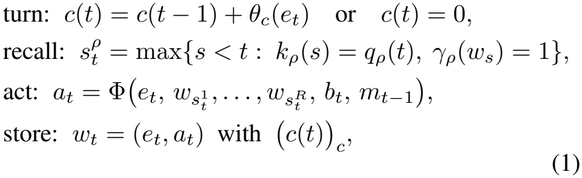

where *e*_*t*_ is the event at moment *t*, 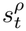 the moment whose record recall *ρ* returns, *R* the number of recalls, *b*_*t*_ the sides of the thresholds and *a*_*t*_ the act. What the table receives at a moment is its *situation*.

The table never receives the reading of a counter, only coincidences: what a recall returned, and on which side of zero a counter stands. The first paper proved two consequences: a table fixed on short inputs holds at every length, since a program meets the same finite set of situations at every length; and the memory computes *in step*, acting once for each symbol heard and each answer due, everything that a multitape Turing machine computes in step.

A program is thus a set of counters with their gains, recalls, thresholds, states and a table: the counters and recalls decide which past moment stands beside the present, and the table says what to do with it. To learn a program is to find both. The first paper found them together, by gradient descent through a relaxation of Eq. (1). Here they are found apart, by comparing experiences.

## 3. Learning by Comparison

Comparisons between remembered experiences write the parts of a program. Appendix B states each result below in full and proves it, Appendix A works the examples by hand, and Figure 2 shows them at work.

**Figure 2.**
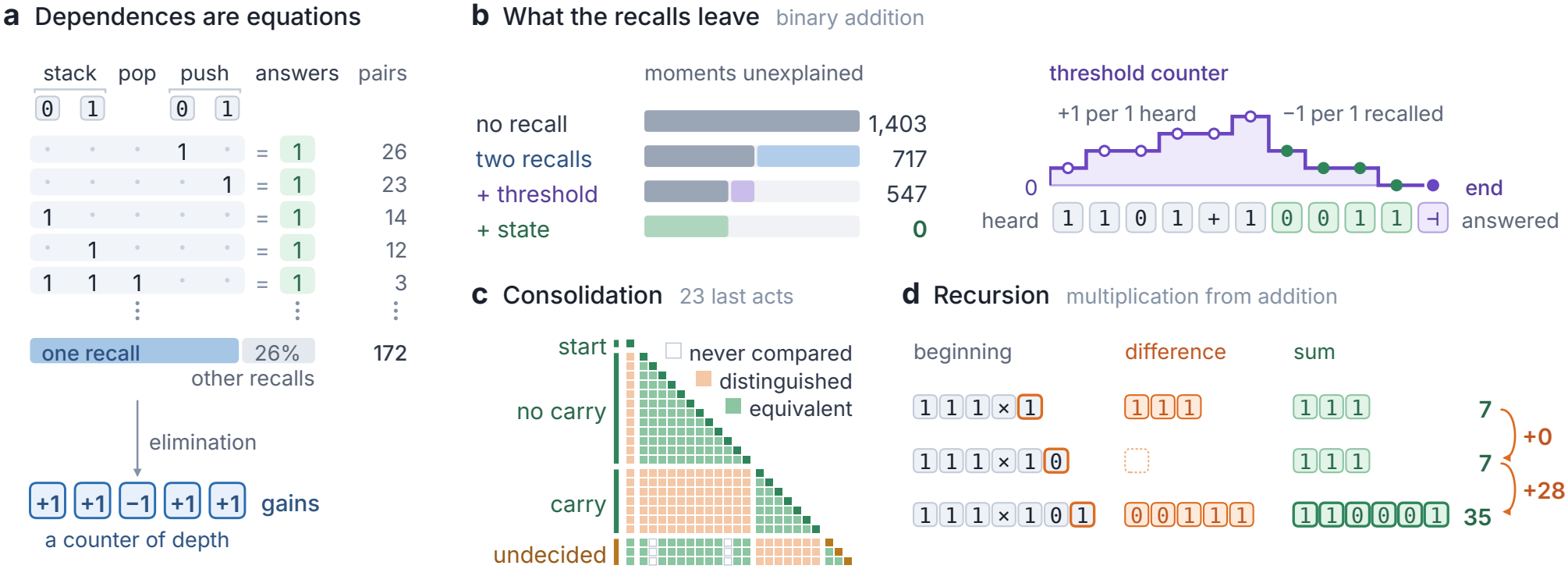
From comparisons to a program. **(a)** Each pair of dependences of the stack gives a linear equation in the gains of five kinds of symbol, counted between the two records recalled and between the two answers (right: how many pairs give each equation); 26% of the 172 pairs belong to other recalls. Elimination on random draws of the equations, keeping the solution on which most of them agree, returns a counter of depth. **(b)** Binary addition: moments left unexplained as each part is added (tint: explained), and the threshold counter on one sum. **(c)** Consolidation sorts the 23 last acts of addition into three states; three stay undecided. **(d)** Recursion on 111*×* 101 (7 *×*5, lowest digit first): differences between the answers to successive beginnings, summed with the learner’s own addition, give the product.

### 3.1. Credit is the first difference

A *question* is a string that can be heard, and a task assigns to each question the answers that are due. Two experiences form a *minimal pair* if their questions differ in one place, as 10110 and 11110 do in the Introduction; the answers, their reversals, first differ at the fourth.

#### Definition 1

(Dependence). In an experience, the answer at moment *t depends* on an earlier moment *s*, written *s* ↣ *t*, if the experience forms a minimal pair with another one that differs from it at *s*, and *t* is the first moment at which their answers differ.

Up to moment *t* the two experiences differ at *s* alone, so whatever procedure produced the answers, something of moment *s* reached the answer at *t*; in a program of Eq. (1), it travels through a recall, a counter or the state (Lemma 1). This is Mill’s method of difference (Mill, 1843), and it spans any distance in one step. Later differences can be consequences of the first, carried forward by the learner’s own answers as a carry is in addition, so the first difference is the one that counts.

#### Theorem 1

(Credit). *If an answer is a sum modulo two of independent, uniform causes, and every other record is independent of each cause given the other causes, no statistic of single experiences that leaves one cause out tells a cause from a non-cause. In a minimal pair, for every procedure, the answer at the first difference is not a function of the rest of what was heard and said before it, however many moments lie between the place that differs and that answer*.

Statistics of single experiences thus credit a cause that acts jointly only together with all the others, which is why parities are hard to learn from statistics (Kearns, 1998; Blum et al., 1994) and why rules of co-occurrence, such as Hebb’s (Hebb, 1949), fail to assign credit to joint causes. A comparison holds everything fixed but one place, as a controlled experiment does, and finds each joint cause at once; in the theory, learning combinations of cues in a step depends on the hippocampus for this reason (Section 5.1). Minimal pairs abound in short experience, where the 32 strings of five bits hold 80. They are also made, by a learner that changes one symbol of a question and asks for its answers, and by speech to young children, some of which repeats an utterance with one part varied (Küntay & Slobin, 1996).

### 3.2. Three neighbours

Each of the three kinds of neighbour that an experience has in memory writes a different part of a program. The neighbour by *substitution*, the other experience of a minimal pair, shows what each answer depends on: it writes the recalls. Two beginnings of experiences are neighbours by *continuation* when the same string is heard after both, *distinguished* if the answers to it differ and *equivalent* if they agree. In parity, 1 and 111 are equivalent under every continuation and 11 is distinguished from both, by the empty continuation or by 01 (Figure 1c), so parity needs two states. Continuation shows what has to be remembered: it writes the states. The neighbour by *deletion* of a question is the question without its last symbol. It shows how an answer grows with its question: it writes recursion.

#### Postulate

(Learning by comparison). A brain turns experience into a program by comparing each experience with its neighbours in memory. (i) *Credit is the first difference*. What an answer depends on is found by comparing the two experiences of a minimal pair. (ii) *A dependence is an equation*. The gains of a program’s counters are the solution of its dependences. (iii) *The rest is difference again*. What the recalls leave is a threshold, solved for where the answer changes, or a state, which carries what neighbours by continuation distinguish; consolidation merges what experience has shown equivalent and never distinguished. (iv) *Nothing is guessed*. A program found from the shorter experiences is announced only if it answers the longest ones; where experience leaves two possibilities, the learner keeps both, and asks or says that it does not know.

Part (i) takes the place of the backward pass, and part (ii) the place of search. We call the memory of Eq. (1) with this way of learning the *learner* (Algorithm 1, Appendix C).

### 3.3. A dependence is an equation

In reversal of five symbols, the fourth answer depends on the second symbol and the fifth on the first. The records lie one symbol apart and the answers one answer apart, in the opposite order, so a recall whose key and query are one counter serves both only if an answer turns the counter back as far as a symbol turns it forward (Figure 1b): an equation in the gains.

In general, a recall can serve a dependence *s* ↣ *t* only if its key counter stood at *s* where its query counter stands at *t*. A counter with gains ***θ*** that has not been reset stands at ***θ*** *·***N**(*t*) plus its start, where **N**(*t*) counts the events of each of *K* kinds up to moment *t*. For a key counter with gains ***θ*** and a query counter with gains ***θ***^*′*^ started *δ* higher, which together form the *relation* of the recall, the dependence asks

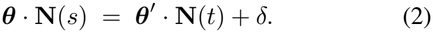

#### Theorem 2

(Dependences determine the counters). *Eq*. (2) *is linear in the gains, and Gaussian elimination solves it; at most* 2*K* + 1 *dependences of one recall, from short strings, fix its solution at every length. Nested dependences, serial ones, and an answer that changes when one count reaches another give the stack, the queue and the counter of computer science, a register that events raise and lower and that is tested for zero*.

Two dependences that one recall serves give an equation free of *δ*, so the learner works with pairs of dependences. The stack task of the benchmark has five kinds of symbol: two that fill the initial stack, a pop and two pushes. Never told that a stack needs a counter of depth, the learner writes it down as the solution (+1, +1, *−*1, +1, +1) on which most of the 172 pairs of dependences agree; the others belong to other recalls (Figure 2a). The query counter is built from the key counter and a few counters that the learner has from the start, such as the count of all symbols heard; of the 1,695 queries so built, the learner keeps the one that serves the most dependences (Appendix C).

### 3.4. The rest is difference again

In binary addition, minimal pairs find two recalls, which return the two digits of a column, and leave the carry and the end of the sum unexplained: at some moments the recalls return the same digits and the answers differ. Finding the causes of such a residue is Mill’s method of residues (Mill, 1843). An earlier moment reaches a later answer in one of three ways (Lemma 1): its record is recalled, it turns a counter that a recall compares or a threshold reads there, or it passes from moment to moment in the state. Minimal pairs find the recalls and the counters they compare. The end of a sum is of the second kind, a threshold: a counter turned forward by each 1 heard and back by each 1 recalled reaches zero when nothing is left to add (Figure 2b).

The state is fixed by neighbours by continuation: beginnings that some continuation distinguishes need different states, and by the theorem of Myhill and Nerode the classes of beginnings that none distinguishes are the states of the smallest program (Myhill, 1957; Nerode, 1958). When answers come throughout an experience, the last act, the last answer with what the recalls returned with it, can carry the state. When a task has one answer at the end, the learner speaks to itself after each symbol, naming the class of the beginning heard so far. We call this *inner speech* (Vygotsky, 1986); in bracketed arithmetic a recall keyed by depth brings back, at each closing bracket, the state named at the matching opening one.

Partial experience needs two changes to the theorem of Myhill and Nerode. The candidates for a state are the values that the last act takes, or, in inner speech, the beginnings heard. Consolidation merges two of them only if experience has shown them equivalent and never distinguished, and a candidate equivalent to two that are distinguished is merged with neither and stays *undecided*. Learners of automata that seek the fewest states choose among many solutions (Heule & Verwer, 2010; Gold, 1978), and so guess; consolidation does not search. It needs all experiences together, and we place it in sleep (Appendix B.7).

#### Theorem 3

(What comparison leaves). *If two moments have different answers although the event and what a correct program’s recalls return are the same at both, they differ in the side of a threshold, found by elimination where the answer changes, or in the state. Consolidation never puts two distinguished candidates in one class, nor merges an undecided one; from complete experience its classes are those of Myhill and Nerode*.

In binary addition, consolidation sorts the 23 last acts into three states, the beginning, carry and no carry, and leaves three undecided (Figure 2c). An undecided candidate answers only where the classes it may belong to agree; elsewhere the program says that it does not know, or the learner asks. Asking needs an act in the world, so the learner asks last, when nothing it can do by itself (Section 3.6) has settled the candidates.

### 3.5. Nothing is guessed

Each part of a program the learner writes is a statement about coincidences, which short strings verify completely; that the strings were long enough is checked by a test on the longest, the sampled test of a conjecture in automata learning (Angluin, 1987).

#### Theorem 4

(Nothing is guessed). *Fix bounds on the gains and starts of the counters and on the numbers of recalls, thresholds and states. There is a length such that a program within these bounds that is right on all inputs up to it is right at every length, or, if its counters are periodic, as grid modules are, at every length within their range. A program found from shorter strings that errs on a share ε of the longest strings passes a test on m of them, drawn at random, with probability at most* (1 *− ε*)^*m*^; *one of k candidates, at most k*(1 *− ε*)^*m*^.

A wrong answer needs two moments with different answers due taken for one (Appendix B.8), and where experience leaves two possibilities equal the learner chooses neither. A false program is therefore one that the test failed to catch, and for errors at the length tested the theorem bounds how often that happens.

### 3.6. Experience made by the machine

What the learner says is stored like what it hears, and turns counters: an answer is an event. The learner can therefore learn from experience that it makes for itself.

#### Theorem 5

(Experience made by the machine). *Let the learner have learned a program P and an addition ⊕. Dependences of a task on what P answers are found by comparison, however long P took (composition). If one symbol of a question stands for its answer and the learner predicts the last symbol of every true statement, it finds the answer by trying each candidate in that place (trial). The answer to a question is the sum under⊕ of the differences between the answers to its successive beginnings; if the differences have a program in step, the sum takes a number of steps of the order of the square of the length, although the task itself may have no program in step (recursion)*.

Trial is the method of difference applied to imagined experiences. For (*x*+2)=4 modulo five, the learner imagines the statements with each digit in place of *x*, which differ in one place, and its own prediction of the last symbol says which of them is true. Recursion is the method of differences that Babbage’s engine was designed to carry out (Babbage, 1864). The answer to 7 *×*5, heard lowest digit first as 111 *×*101, exceeds that to its neighbour by deletion, 7*×* 1, by the multiplicand shifted two places. Such differences have a program in step of 11 situations; added with the learner’s own addition, they give long multiplication, a procedure of quadratic time for which the learner finds no program in step (Figure 2d).

## 4. Experiments

Six experiments test six claims. Comparison alone writes exact programs for most tasks of a standard benchmark from a few hundred short strings, where networks, and gradient descent on the same memory, do not (Experiment 1). The learner, which does not guess, announces no false program, even from thin experience (Experiment 2). Each part of the theory carries tasks of its own (Experiment 3). Without minimal pairs no recall is learned, and questions make the pairs that experience lacks (Experiment 4). The learner solves tasks withheld from its design, and the tasks it solves stay the same when one of its constants at a time is set below or above its value (Experiment 5). It extends its own experience, and builds long multiplication from addition (Experiment 6). Experiment 7, in Section 5.5, runs the engine in circuits of neurons.

### Common setup

A task says which answers are due for each string. In binary addition, for instance, the learner hears 101+100, five and one written lowest digit first, and the answers due are 011, six, and a mark for the end. A string heard with its answers is an *experience*. Experience of size *N* holds 64 random strings of each length from 1 to *N*, repeated strings removed; the *standard experience* has *N* = 12, a median of 457 distinct strings. Where allowed, the learner may also ask a question of its own, a membership query (Angluin, 1988): it chooses a string shorter than the longest heard and is told its answers. The learner (Algorithm 1) has four constants, set once for all tasks (Appendix C.5). A program it announces is tested on 32 fresh strings of every length from 41 to 500 and, if it runs in step, on strings of a million symbols. A run ends with an *exact* program, right on every test string; a program that is never wrong but says on some string that it does not know; a *false* program, wrong on some string; or no program. Each condition has ten seeds, independent draws of experience, and a task is *solved* if all ten are exact.

### 4.1. Experiment 1: the benchmark without backpropagation

#### Why

The benchmark of Delétang et al. (2023) asks whether a procedure learned on strings of up to 40 symbols holds on strings of 41 to 500, and LSTMs and Transformers fail on most of its fifteen tasks (Table 1). If comparison writes programs, a few hundred short strings should do. *Design*.

The learner hears the fifteen tasks at sizes from *N* = 4 to 20. The LSTM, Transformers and Tape-RNN of Table 1 saw at least 10^6^ batches of 128 fresh strings. The first paper’s machine, which runs programs in the same memory, is trained by gradient descent through time on ten of the tasks, from the strings the learner heard and from fresh strings of up to 40 symbols (Appendix D.2).

#### Result

From the standard experience the learner writes exact programs for thirteen tasks, for every seed, in a median of 0.5 seconds on one core, and eleven hold at a million symbols (Table 1, Figure 3a; Appendix C.6). Eight run in step with the state in their last act, and three with the state in inner speech; equations are solved by trial and multiplication by recursion, the two tasks for which the learner asks questions (Experiment 6). Arithmetic modulo five needs more strings (Experiment 4), and for the square root the learner announces no program. Gradient descent on the same memory, in 165 accelerator hours, reaches a high mean accuracy on several tasks, as in the first paper (Table 1, He & Wu), and is exact for every seed on at most five of its ten tasks, never on sorting, addition or the stack (Table 14). *Reading*. On the same strings of at most ten symbols, the learner solves all ten of these tasks and gradient descent three. With one memory and one experience, the difference lies in how the program is found: a few dependences fix a relation (Theorem 2), and a program of coincidences that is right up to a length set by the bounds on its parts is right at every length (Theorem 4).

**Figure 3.**
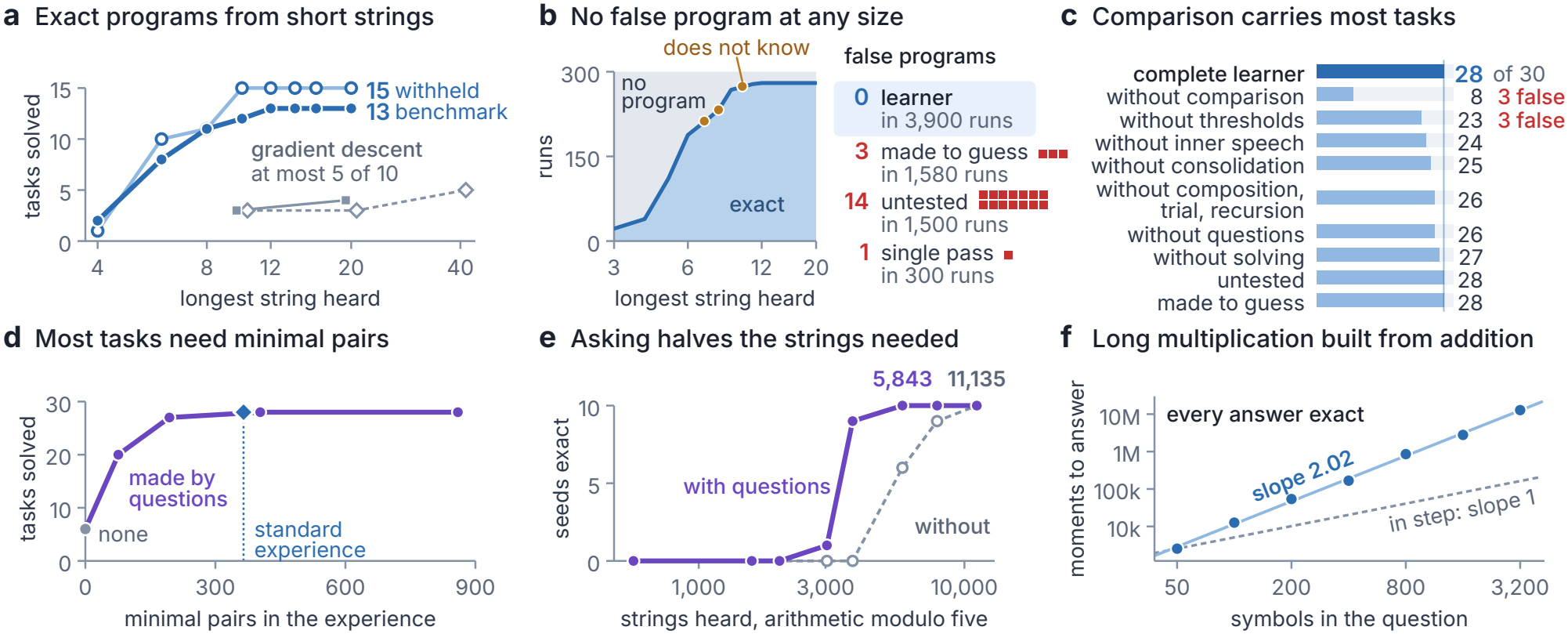
Programs from short experience. Solved: exact for every seed. **(a)** Experiments 1 and 5: tasks solved against the longest string heard; grey, gradient descent through time on the first paper’s machine, 165 accelerator hours in all (squares: the learner’s strings; diamonds: fresh strings). **(b)** Experiment 2: what the learner announces at each size, and a square for each false program of three variants. **(c)** Experiment 3: one part removed; the last two rows are variants of (b). **(d, e)** Experiment 4: minimal pairs, made by questions in a tenth, a quarter, half or all of the experiences (purple), and arithmetic modulo five with questions and without. **(f)** Experiment 6: moments to answer grow as the power 2.02 of the length (dashed: a program in step); every answer is exact up to 3,200 symbols.

### 4.2. Experiment 2: a learner that knows what it knows

#### Why

A learner that extrapolates can be wrong where nobody checks, most of all where experience is thin; by Theorem 4, a learner that does not guess should seldom be. *Design*. A census runs thirty tasks, the benchmark and the fifteen with-held tasks of Experiment 5, at thirteen sizes from *N* = 3 to Three variants each lack one safeguard of the postulate’s part (iv), and each hears the same experiences as the complete learner. At five sizes from *N* = 3 to 11, one is made to guess, taking the best supported of the candidates that experience leaves (also on arithmetic modulo five with more strings), and one announces without the test on the longest strings. The complete learner’s consolidation compares its candidates again until nothing changes; the third variant compares them once, so that early merges stand. It hears the experiences without minimal pairs of Experiment 4, where the learner falls back on inner speech, whose table grows state by state, so that an early merge can meet a state found later.

#### Result

Of the 3,900 census runs, 2,742 gave an exact program and 1,152 no program; six gave a program that never erred but said on some string that it did not know, and none a false program (Figure 3b), as in all 5,680 runs of the complete learner in this paper. The variants announced false programs: 3 in 1,580 runs made to guess, 14 in 1,500 untested, 1 in 300 with a single pass. *Reading*. Each false program of a variant answers every string it heard and errs beyond them, and each error is one that the missing safeguard would have prevented; one is an addition that merged a candidate the complete learner leaves undecided (Appendix D.3). Where experience is thin, the complete learner reports that it has no program, and why.

### 4.3. Experiment 3: what each part is for

#### Why

If the postulate describes how programs are learned, each of its parts should carry tasks that the others cannot reach. *Design*. Over the thirty tasks from the standard experience, one part of the learner is switched off at a time.

#### Result

The complete learner solves 28 tasks, as it does untested or made to guess; these variants err where experience is thin (Experiment 2; Figure 3c). Without the comparison of minimal pairs no dependence is found, and eight tasks remain, those whose programs are thresholds or inner speech. Without solving for counters the stack is lost, since no counter the learner starts with counts depth; without thresholds, addition, multiplication, majority and the sorts; without inner speech, parity, cycle navigation, bracketed arithmetic and equations; without composition, trial and recursion, or without questions, multiplication and equations; without consolidation, addition, subtraction and addition in base three. The few false programs come from the withheld task majority, whose program is one threshold (Appendix D.4). *Reading*. Each part carries tasks of its own, those that the theory assigns to it.

### 4.4. Experiment 4: minimal pairs and questions

#### Why

In the theory every dependence comes from a minimal pair: without pairs the tasks that need a recall should be lost, and questions should bring them back. *Design*. The standard experience is rebuilt without minimal pairs, drawing other strings in place of those that would form one: a median of 388 strings remains of 454 (medians over the thirty tasks). Then the learner changes one symbol in a share of its experiences and asks for the answers, making minimal pairs. Arithmetic modulo five is given more strings, with questions and without.

#### Result

Without minimal pairs six of the thirty tasks are solved, all by programs without a recall, and every run without an exact program reports that it has none (Figure 3d). The loss comes from the pairs: the census experience of *N* = 11, as large (390 strings) and with its pairs, solves 27, 26 of them without a question. With a question for every tenth experience 20 tasks are solved, and with one for every second 28. Arithmetic modulo five is exact for every seed from 5,843 strings with at most one question, and from 11,135 without (Figure 3e). *Reading*. Recalls are learned from minimal pairs, and a learner that can act makes the pairs it lacks. A question is a neighbour made on purpose: by substitution here, by continuation in arithmetic modulo five, by deletion in recursion.

### 4.4. Experiment 5: withheld tasks and the constants

#### Why

A learner designed while looking at fifteen tasks may be fitted to them, in its steps or in its constants. *Design*. Fifteen further tasks, among them exclusive or and conjunction of two strings, subtraction and majority, were fixed before the learner was written and never used in its design; they are learned as in Experiment 1, without questions. Each of the four constants is set below and above its value, one at a time.

#### Result

From the standard experience the learner solves all fifteen withheld tasks, in step, in at most 1.6 seconds each, with no false program in 1,200 runs (Figure 3a). Under each of the eight settings of the constants it solves the same 28 of the thirty tasks, with no false program (Appendix D.6.1). *Reading*. The same comparisons find tasks of new kinds, among them the exclusive or of two strings, the case of Theorem 1 in which each digit alone is independent of the answer. The constants set how strong evidence must be, and in this range they change the outcome of no task.

### 4.5. Experiment 6: experience made by the machine

#### Why

For some tasks the learner finds no program in step, among them a sum said highest digit first and multiplication; by Theorem 5, a learner with programs already learned reaches them from experience that it makes itself. *Design and result. Composition*. Four tasks are said highest digit first, so that the first answer depends on the last symbol heard. With the program of the same task said lowest digit first already learned, the learner solves all four (one of them, the increment, in step without composition): it computes the answer and reads it backward. *Trial*. For an equation such as (*x*+2)=4, modulo five, it finds the symbol that stands for the answer, learns to predict the last symbol of a true statement, and tries each value in that place. *Recursion*. For multiplication it walks from each string heard down through its beginnings, asking for the answers it does not remember (a median of 990), and finds a program in step for the differences of successive answers. The program it announces is long multiplication, exact at every length tested, up to 3,200 symbols, in a number of moments that grows as the power 2.02 of the length (Figure 3f). Without questions it finds none: a fifth of the strings heard have their neighbour by deletion in memory.

#### Reading

What the learner says is experience like what it hears. The program of the differences is one row of the schoolbook procedure, and the announced program adds the rows: a procedure of quadratic time built from programs in step, by Babbage’s method of differences.

## 5. The Difference Engine in the Brain

### 5.1. Why a brain keeps episodes

A comparison takes two experiences, and an animal lives through one at a time, so the second has to come from memory. That memory has to store each experience whole after one exposure, since the experience that will later form a minimal pair with it cannot be known in advance; to store similar experiences apart, since the experiences worth comparing are nearly the same; to recall a stored experience by its likeness to the present; and to compare the two.

These are the functions attributed to the hippocampal system (Figure 4a; Appendix E.1). It stores an episode after one exposure (Marr, 1971; Nakazawa et al., 2003), as an index to the cortical pattern of the moment (Teyler & DiScenna, 1986), through plasticity that binds events seconds apart (Bittner et al., 2017). The dentate gyrus gives similar experiences distinct codes (Leutgeb et al., 2007; McHugh et al., 2007), and CA3 completes a partial cue to a stored pattern (Nakazawa et al., 2002). CA1 receives the recalled pattern from CA3 and the present one from entorhinal cortex, in separate gamma rhythms (Colgin et al., 2009), and is thought to compare them (Lisman & Otmakhova, 2001). Its response in people grows with the number of changes between what is recalled and what is seen (Duncan et al., 2012), and the human left hippocampus signals novelty “only in conditions where one contextual component was novel and the other repeated” (Kumaran & Maguire, 2007a), the condition of a minimal pair. The theory reads the four functions as steps of one operation, whose result is the moment on which an answer depended.

**Figure 4.**
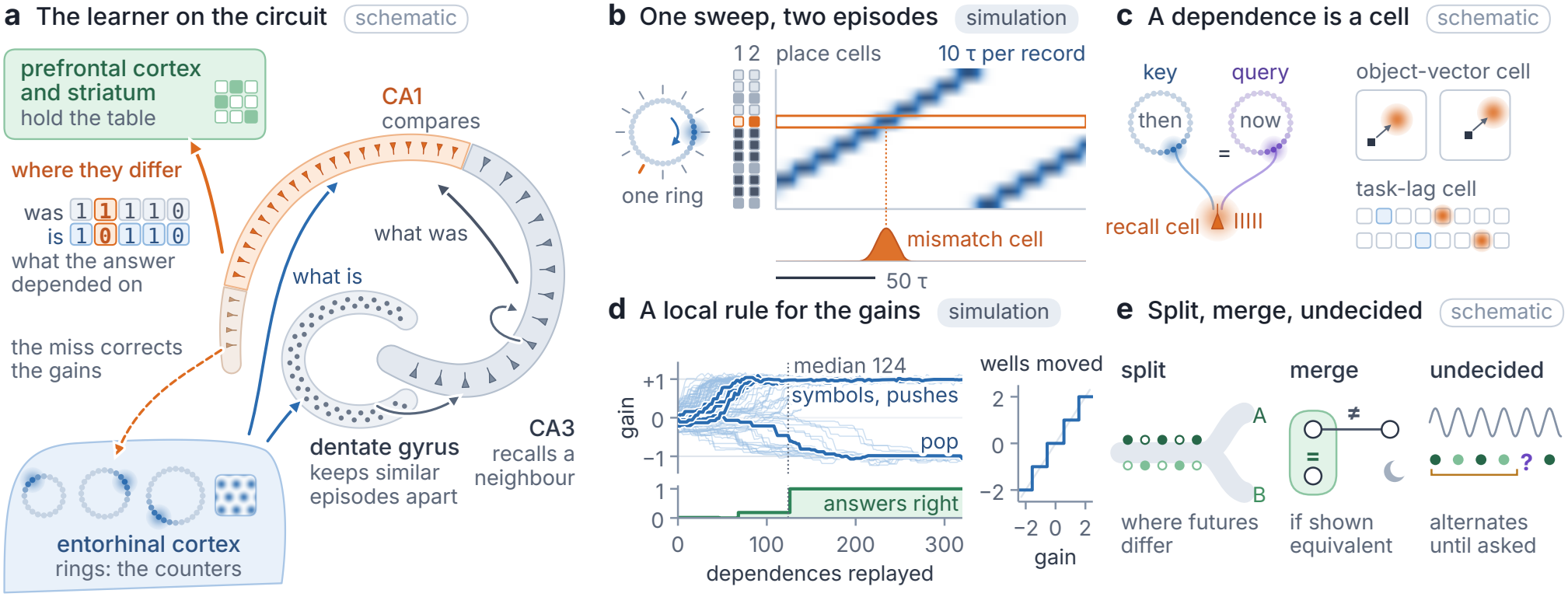
The difference engine in the brain. **(a)** The learner on the hippocampal circuit: rings in entorhinal cortex, grid modules among them, are the counters; the dentate gyrus keeps similar episodes apart, CA3 recalls a neighbour, and CA1 compares what was (from CA3) with what is (from entorhinal cortex); where they differ is what the answer depended on, and the miss for that dependence corrects the gains (dashed). Prefrontal cortex and striatum hold the table. **(b)** Experiment 7: two episodes stored along one ring and replayed in one sweep; a cell fires only where they differ. **(c)** A recall cell fires when the query ring stands where the key ring stood; in the theory, object-vector and task-lag cells hold dependences in this way. **(d)** Experiment 7: from all its dependences, unsorted, the local rule learns the stack’s counter (20 of 20 runs); the wells make each gain whole. **(e)** State codes split where futures differ, merge in sleep when shown equivalent (P6) and, where undecided, alternate until a question settles them (P7).

In the theory, writing a program whose answers depend on a recalled moment needs this operation, and running the program does not: a program runs on its counters and its table, and on the index if it recalls records; a regularity among events that are seen together, which needs no recall of an earlier moment, is learned by counting how often they occur together, with no comparison (Appendix E.5). The dentate gyrus, the stage that keeps neighbours apart, is needed to encode a spatial or contextual memory and not to retrieve it (Lee & Kesner, 2004; Kheirbek et al., 2013); habits and an overtrained maze response are acquired or run without the hippocampus (Knowlton et al., 1996; Packard & McGaugh, 1996); and amnesic patients classify strings of an artificial grammar as well as controls, except when told to judge them by comparison with the strings studied (Knowlton et al., 1992).

Theories of the hippocampus have been pursued largely apart (Eichenbaum & Cohen, 2014), each built on one finding; the theory accounts for these findings in turn. Memory for facts and events needs the hippocampus, and learning skills does not (Cohen & Squire, 1980; Squire, 1992): in the theory facts and events are records in the index, and a skill is a table with its gains, learned by counting. Configural association theory held the hippocampus necessary for learning combinations of cues, and later held that it speeds that learning (Sutherland & Rudy, 1989; Rudy & Sutherland, 1995); relational theory holds it necessary for binding items into relations that can be used flexibly (Cohen & Eichenbaum, 1993). In the theory a combination of cues is a set of causes that act jointly, invisible to statistics of single cues and visible to one comparison (Theorem 1), so that comparison learns it in a step where statistics of the conjunctions learn it slowly. In complementary learning systems the hippocampus replays episodes to a cortex that learns slowly (McClelland et al., 1995; Kumaran et al., 2016), and with a schema in place a new association is learned in one trial, with the hippocampus, and soon no longer needs it (Tse et al., 2007). In the theory replay pairs episodes to compare them, and a schema is a written program, which runs in cortex and striatum: a new association, stored in the index after one trial, enters its table with no comparison and soon runs without the index.

Comparator theories read the hippocampal mismatch signal as novelty that gates storage (Vinogradova, 2001; Lisman & Grace, 2005); in the theory it also locates a cause, so that CA1 reports credit. Maps of space and of tasks (O’Keefe & Nadel, 1978; Behrens et al., 2018), which models learn from the statistics of experience (Stachenfeld et al., 2017; Whittington et al., 2020; George et al., 2021; Gershman & Niv, 2010), are in the theory a program’s counters and its states, found from differences and split where futures differ. Replay, taken to carry value back from a reward and to plan (Foster & Wilson, 2006; Pfeiffer & Foster, 2013; Mattar & Daw, 2018), also shows, in the theory, the neighbour with the other outcome, whose difference assigns credit at once. Table 2 gives predictions that separate the theory from each account (Appendix E.3).

### 5.2. Two moments in one cycle

In the theory each counter is a ring of neurons, a continuous attractor network (Zhang, 1996) of the kind observed in grid modules and in the heading system of the fly (Gardner et al., 2022; Kim et al., 2017). Its reading is where a bump of activity sits. Wells in the otherwise continuous attractor hold the bump in one of a few stable positions and so make the reading a whole number, which rings that count discrete events, such as laps, should show (P5). An event moves the bump by a pulse whose strength is its gain. Besides distance, hippocampal codes count laps, elapsed time and accumulated evidence (Sun et al., 2020; Pastalkova et al., 2008; Nieh et al., 2021). On this account a place cell detects a coincidence of rings, and a recall cell fires when one ring stands where another stood when its record was stored (Figure 4c). Cells in entorhinal cortex fire at a fixed distance and direction from an object (Høydal et al., 2019), and cells in medial frontal cortex at a fixed lag after a step of a task (El-Gaby et al., 2024); on this account each holds a dependence as a relation between two counters (P4). Experiment 7 compiles the programs that the learner wrote into such circuits and runs them (Section 5.5).

A comparison needs the two experiences of a minimal pair in one cycle of activity, even when they happened days apart. When the neurons of a ring adapt, the bump travels along the ring by itself, and the place cells on its path reactivate their records in order, those of earlier episodes stored at the same readings among them. We take these sequences to be theta sweeps and replay (Skaggs et al., 1996; Foster & Wilson, 2007), which adaptation produces in models (Chu et al., 2024; Ji et al., 2026), and a cell that receives both episodes of one sweep can then report where they differ (Figure 4b; Experiment 7). Replay can run backward from a reward, at a rate that follows the change in reward (Foster & Wilson, 2006; Ambrose et al., 2016), and often shows the other experience that a comparison needs: another environment (Karlsson & Frank, 2009), a path never run, or the arm not chosen (Gupta et al., 2010; Carey et al., 2019). The theory predicts that a surprise brings the present episode and its neighbour of the other outcome into one sweep (P2).

### 5.3. A local rule for the gains

A circuit meets the equations of Theorem 2 one at a time, each a constraint that a coincidence occur (He & Wu, 2026b). The events between a recalled record and the answer are replayed in a sweep, faster than the wells can catch the bump. The bump adds their pulses unrounded and arrives at some distance from the record’s reading: the *miss* of Eq. (2),

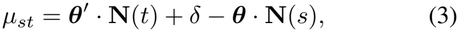

which is graded although the running counter is not.

#### Theorem 6

(A local rule). *(i) For a counter recalled against itself, the force of a dependence on the gain of a kind of event, the negative gradient of half its squared miss, is the miss times the number of events of that kind between the record and the answer, with the sign that reduces the miss. (ii) If some gains serve every dependence, the rule that changes the gains at each miss by this force, divided by the sum of the squared counts, converges to the solution nearest its start when the dependences recur, in turn or at random, as the method of Kaczmarz (1937) does; with wells, if that solution is whole, the counter becomes exact after finitely many dependences and stays exact. (iii) Backpropagation through time computes the same force (Werbos, 1990); the rule computes it when the miss occurs, from the miss and the counts, with no backward pass*.

No error has to travel back from the answer to the record, because the counters have already summed the events between the two: an event turns a counter by an amount that depends on its kind alone. Each event between the record and the answer sends its pulse once in a sweep, so a trace at the synapses of its kind holds the count when the miss arrives: the rule has three factors, as rules with eligibility traces do (Gerstner et al., 2018; Bellec et al., 2020), and the miss is an instructive signal of the kind single neurons receive (Francioni et al., 2026). Landmarks moved in proportion to an animal’s motion change the gain of its path integration, in part and lastingly (Jayakumar et al., 2019), which the rule reproduces with nothing fitted (Experiment 7).

The miss acts through the firing field of the record’s place cell, about one well wide, so a dependence that the counter cannot serve misses by whole wells and is ignored. A ring fed with all dependences, unsorted, thus learns the counter on which most of them agree (Figure 4d; Experiment 7).

Learning is gradual, and what is learned is exact: the synapses that set a gain move by small steps, the wells turn a gain within half a well of a whole number into that number, and the counter becomes exact at the step that brings its gains into that range, and stays exact (Figure 4d; P5). Graded synapses can thus learn and hold an exact program.

### 5.4. What is carried, merged, or left undecided

A program carries only what its continuations distinguish. Cells on a common stretch of a maze fire according to where the animal came from or will go (Wood et al., 2000; Frank et al., 2000), and when mice run two corridors identical except for one cue, the codes of the two move apart over days where the corridors look alike and the task differs (Sun et al., 2025). In the theory these are states, split where a common continuation is answered differently (Theorem 3).

Sleep helps people infer relations between items learned apart (Ellenbogen et al., 2007). In the theory, consolidation, which we place in sleep, merges what experience has shown equivalent and never distinguished: situations met in one context with one outcome, and not situations that only share an outcome (P6). It needs no search: threshold units, one for each candidate, excite the units of the candidates shown equivalent to their own, every distinguished pair vetoes, and the classes are what the spreading activity reaches (Appendix B.7). A consolidated memory becomes labile again after recall (Nader et al., 2000) when the recall meets a prediction error (Sevenster et al., 2013), the condition under which the learner compares a merge again.

In the learner, a moment that experience has not decided stays undecided until the learner asks. We read deliberation and exploration in this way: at an unlearned choice a rat pauses while its sweeps run down one arm and then the other (Johnson & Redish, 2007; Redish, 2016), and children explore more when the evidence they were shown is confounded (Schulz & Bonawitz, 2007). Theta sequences of trained rats alternate between paths or directions on every cycle (Kay et al., 2020; Vollan et al., 2025); the theory predicts alternation between states where the paths are the same, at an undecided moment (P7; Figure 4e).

### 5.5. Experiment 7: the engine in circuits of neurons

#### Why

The account of this section asks four things of neurons: that the programs written by comparison run on rings and place cells, that a sweep brings two episodes together and reports where they differ, that a ring learns the gains of a counter from dependences that nobody has sorted, and that consolidation merges without search.

#### Design

Each counter is held by rings of rate neurons, continuous attractor networks with adaptation in the form used in the first paper (He & Wu, 2026a; Mi et al., 2014). The neuron with preferred phase *θ* has synaptic input *U*, adaptation current *V* and firing rate *r*:

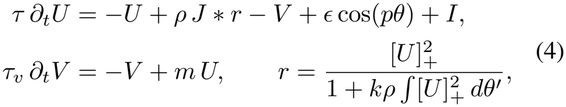

where *J*, a Gaussian kernel of the distance on the ring, holds a bump of activity at any phase, the weak input *ϵ* cos(*pθ*) makes *p* wells, *k* sets the divisive inhibition and *I* is the input from outside the ring. An event turns the ring by a brief asymmetric input whose duration is proportional to its gain. Without adaptation (*m* = 0) the bump stays where it is put; with *m > τ/τ*_*v*_ it travels along the ring by itself. A counter is a set of rings of coprime periods, read by the Chinese remainder theorem, as the modules of the grid system together encode a position (Fiete et al., 2008; Stensola et al., 2012). Five tests follow the parts of this section (Appendix D.8). (a) One event of each gain from *−* 2.5 to 2.5. (b) Each program in step that the learner wrote for the benchmark, compiled into rings, place cells that file each record under the wells of its key counter, latches that hold the side of a threshold, and the program’s table, run on fresh strings of 10 to 120 symbols and then with noise in every neuron. (c) The key counter of the stack, started from random gains and fed all 172 pairs of its dependences in random order, each replayed in a sweep and corrected by the rule of Theorem 6 through the firing field of the record’s place cell; and the same without the field. (d) Two episodes that differ in one record, stored along one ring with adaptation and replayed by its travelling bump to a cell that receives both. (e) The consolidation circuit of Section 5.4, on the tables that the learner wrote. The rule is also run alone, on the dependences of reversal and of the stack, against a backward pass, and on the protocol of Jayakumar et al. (2019) (Appendix D.9).

#### Result

(a) Each event moved the counter by the whole number nearest to its gain. (b) Seven of the eight programs compile into rings, place cells and latches alone; the recall of the eighth, the missing duplicate, needs a counter set to where another stands. On 420 strings of up to 120 symbols, ten times the longest heard, the circuits answered as the programs on all, and with noise in every neuron, on every string tried up to *σ* = 1 (Figure 5a,b). (c) From random gains the ring came to act as (+1, +1, *−*1, +1, +1), the solution that elimination finds, in 20 of 20 trials, after a median of 124 dependences, although 26% of the pairs belong to other recalls; without the field, in 0 of 20 (Figure 4d). (d) The bump passed the stored records in order, 10 time constants a record, and the cell fired only at the record where the episodes differ (Figure 4b). (e) The circuit sorted the candidates into the learner’s classes in 46 of 46 cases, within 4 sweeps of spreading activity. Run alone, the rule reached the program in every run. Its correction equals the gradient of a backward pass to 2 *×*10^*−*13^, and parts from it once the turn that an event gives depends on the counter’s reading (Figure 5c). In the recalibration protocol the gain kept after the landmarks are switched off lies between one and the landmarks’ gain, the more so for a faster rule, and it stays there, as in rats, with nothing fitted (Figure 5d).

**Figure 5.**
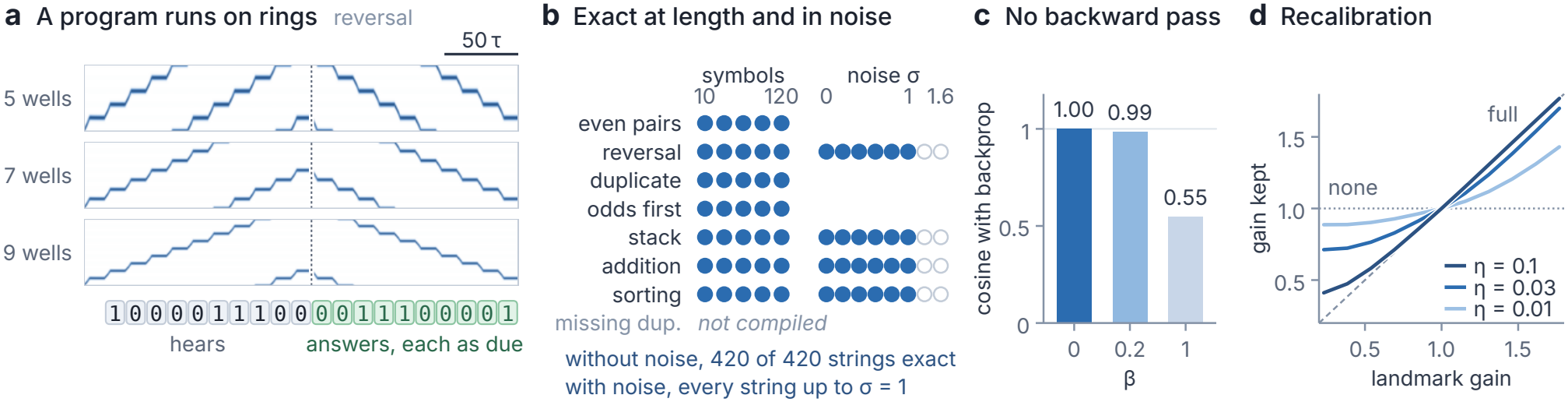
The engine in circuits of neurons. (Experiment 7; every panel is a simulation). **(a)** The program for reversal that the learner wrote, compiled into rings, place cells and its table, reverses a fresh string of ten symbols. Shown is the rate of every neuron of the query counter’s three rings, of five, seven and nine wells: the bump moves one well forward for each symbol heard and one back for each answer, and each ring wraps at its own period, as the modules of the grid system do. **(b)** Each program in step, compiled into rings, place cells and latches, answers as the program does on fresh strings of 10 to 120 symbols (filled: on every string), and with noise of strength *σ* in every neuron (four programs). The recall of the missing duplicate needs a counter set to where another stands. **(c)** Mean cosine between the correction that the rule computes from the miss and the counts, and the gradient of a backward pass; *β* sets how strongly the turn that an event gives depends on the counter’s reading (0 in the memory of Eq. (1)). **(d)** The protocol of Jayakumar et al. (2019): the gain kept after landmarks moved in proportion to the animal’s motion are switched off, for three rates *η* of the rule; dashed, full recalibration; dotted, none.

#### Reading

Marr’s three levels (Marr, 1982) are here one system. What elimination solves, the ring learns by a rule that uses only the miss at a recalled record and the counts of events; what the ring learns acts as whole numbers because of its wells; and the programs written by comparison run on the rings unchanged, at lengths never heard. The ring also supplies what the rule needs. The miss is graded because a sweep outruns the wells, and the dependences need no sorting by recall, because those that a counter cannot serve miss by whole wells, beyond the field of a place cell.

## 6. Discussion

A gradient and a difference are two ways of asking what an answer depended on. The gradient asks the computation, by changing it a little and following the change back through every step. The difference asks experience, by finding two experiences that differ in one place and reading off where their answers part. The temporal difference of reinforcement learning compares two moments of one experience (Sutton, 1988); the difference engine compares one moment of two experiences. The theory reads the hippocampal system’s costly memory of whole episodes as the brain’s solution to the problem of credit. Babbage’s difference engine was designed to tabulate functions without multiplying, because differences reduce the work to addition (Babbage, 1864); the engine proposed here writes programs without backpropagation, because differences reduce the work to solving and counting.

Hume held that causes are known by custom (Hume, 1748/2000), and a network trained on examples learns by custom: what it acquires is a habit, strong where the examples were many. Among his rules for judging causes is also the one used here: “the difference in the effects of two resembling objects must proceed from that particular, in which they differ” (Hume, 1739–1740). Custom needs many instances, and the rule needs two that differ in one place. Mill observed that nature seldom offers the pair of instances this rule requires, and that an experiment makes one (Mill, 1843). A memory of episodes keeps nature’s rare pairs, and a question is the experiment.

The two modern forms of the problem of induction also have an answer for this learner. Goodman asked which predicates can be projected beyond the evidence, since *grue* fits every emerald seen so far as well as *green* does; Kripke asked what fact about a speaker’s past use fixes that she meant *plus*, since *quus* agrees with it on every sum she has computed (Goodman, 1955; Kripke, 1982). The answer lies in the format of the learner’s memory. A rule that agrees with experience up to some point and parts from it there has to know how far a count has gone, and a table that receives only coincidences knows this only within the bounds on its parts. Within those bounds, a rule that parts from the task does so at or before a length that the bounds fix (Theorem 4), and the test on the longest strings can refute it, as a test refutes a conjecture (Popper, 1959).

The theory also bears on an old question of cognitive science: whether a network of neurons can hold rules that apply to every input, as systematic thought seems to require (Fodor & Pylyshyn, 1988; Smolensky, 1988; Marcus, 2001). In the difference engine the learning is graded and the program is exact. Generality comes from a table that receives only coincidences (Theorem 4); exactness comes from the wells of the attractors, which round graded gains to whole numbers (Section 5.3). The symbolic and the connectionist descriptions then apply to one circuit at two time scales: the program it runs, and the slow change of its gains.

For machine learning, the learner opens a way between two traditions. Networks trained by gradient descent extrapolate when they are given a stack, a tape or a grid, or positions encoded for the task, after millions of examples (Joulin & Mikolov, 2015; Kaiser & Sutskever, 2016; Delétang et al., 2023; McLeish et al., 2024); program synthesis writes exact programs by searching spaces that grow exponentially with program size (Solar-Lezama, 2008; Ellis et al., 2021), and Summers’ method builds recursive programs from the differences between successive examples (Summers, 1977). Like a network, the learner learns from examples, and like program synthesis it writes exact programs, which it finds by solving equations. Its two ingredients, kept episodes and counters, mark its place between the two (Proposition 8, Appendix B.11): without kept episodes a learner credits causes that act jointly only in sets or along a gradient, and without counters it carries what it learned to other lengths only by search.

Its questions are the membership queries of automata learning (Angluin, 1987), and a minimal pair is an intervention on one variable (Pearl, 2009). Rules proposed for learning in the brain approximate the backward pass by feedback, local traces or the contrast of two phases of one network (Lillicrap et al., 2016; Scellier & Bengio, 2017; Bellec et al., 2020), and agents carry value across long delays by recalling stored episodes (Hung et al., 2019). Such agents recall one past moment and carry value back to it; the difference engine recalls a second experience, like the first except at the cause, and finds the cause where the two differ. Interpretability finds the circuits of trained networks by patching activations between inputs that differ in one word (Vig et al., 2020; Geiger et al., 2021; Wang et al., 2023); in the theory, the brain writes its circuits by the same comparison.

The first paper showed that this memory can run everything a Turing machine computes. Experience writes less: programs in step, which composition, trial and recursion extend to procedures of quadratic time such as long multiplication. It writes neither a procedure whose parts were never shown, such as the square root, nor the stored program of Turing’s universal machine, which keeps its own table in memory and recalls it as it recalls the world. The theory treats memory, tasks and kinds of event as exact; with noise a single difference becomes a statistic over many pairs, and an animal takes days where the learner takes seconds (Sun et al., 2025). Its predictions concern what is computed and in what order, and the first of them is the most direct test of the principle: a rule with joint causes should be learned in a step from minimal pairs, and slowly from experiences that offer the same statistics without them.

In the *Meno*, true opinions become knowledge when “fastened by the tie of the cause”, and “this fastening of them, friend Meno, is recollection” (Plato, 1892). In the difference engine what is recollected is another experience, and the tie is the difference between the two.

## Impact Statement

This paper presents work whose goal is to advance theoretical neuroscience and machine learning: a principle for how neural systems turn remembered experience into procedures without backpropagation, tested in simulation on a standard benchmark and on fifteen tasks withheld from the design. No human participants or animal experiments were involved. There are many potential societal consequences of our work, none of which we feel must be specifically highlighted here.

## AI Use Statement

## Appendix

The appendix has three parts, on the theory, the experiments and the brain, followed by the related work and the code. Every entry of the contents below is a link.

### The theory

Appendix A defines the terms of the paper, with their symbols and their names in the code, and works the main examples by hand, from a single minimal pair to long multiplication. Appendix B states each result of Section 3, and the local rule of Section 5.3, in full and proves it. Appendix C gives the learner as an algorithm, its four constants, and the programs it writes for the benchmark.

### The experiments

Appendix D gives the protocol that the experiments share, then each experiment of Section 4 in full (why it was run, its design, its result and what it means), then Experiment 7 of Section 5, the engine in circuits of neurons and the local rule of Theorem 6.

### The brain

Appendix E sets each part of the learner beside the structure of the brain that the theory takes to carry it, with the evidence, and gives a protocol for each of the eleven predictions.

### Related work and code

Appendix F places the theory in the work on learning procedures, credit assignment, causal inference, development and the hippocampus, and Appendix G gives the code and the commands that reproduce every number, table and figure.

*To follow the proofs*, read the glossary (Appendix A.1) and the counter of the stack worked by hand (Appendix A.4), then Appendix B in order. Table 6 at its start lists every result with the places where it is stated and proved, and Appendix C.2 shows where the learner uses each.

*To reproduce an experiment*, start from Table 10, which names each experiment in the code with the command that runs it, and from the protocol (Appendix D.1). Appendix G gives the commands in order, what each writes and how long it takes; make demo shows the learner at work in about two minutes.

*For the predictions*, start from Table 2 in Section 5, then read the protocols (Appendix E.4), each with the result it follows from, what the competing account expects and the outcome that would refute the theory. Appendix E.1 gives the evidence for the correspondence, Appendix E.3 how one requirement accounts for findings on which theories of the hippocampus divide, Appendix E.6 the results that would refute the theory, and Appendix F.6 the theories themselves.

### A. Glossary and worked examples

This appendix is for a reader who wants the terms of the paper defined in one place, or who wants to follow the theory by hand on small cases before reading the proofs. Appendix A.1 collects the terms with their symbols and their names in the code. Seven examples follow, in the order of Section 3: a minimal pair and the dependence it shows (Appendix A.2), the program for reversal (A.3), the counter of the stack solved from its dependences (A.4), the states of parity (A.5), binary addition with its recalls, threshold and states (A.6), a question that settles an undecided moment in arithmetic modulo five (A.7), and multiplication built from differences (A.8). Every number in the examples is the one that the learner’s code gives. The programs are those that the learner writes from the standard experience with seed 0 (Appendix C.6), unless another experience is named.

#### A.1. Glossary

The code has names of its own for several terms: a counter is a *clock* there, a recall a *read*, a minimal pair *twins*, a dependence a *need*, a program a *law*, and consolidation *sleep*. Tables 3, 4 and 5 give, for each term of the paper, its meaning, its symbol where it has one, and its name in the code. Table 25 in Appendix G goes the other way, from the names of the code to the terms of the paper.

**Table 3.**
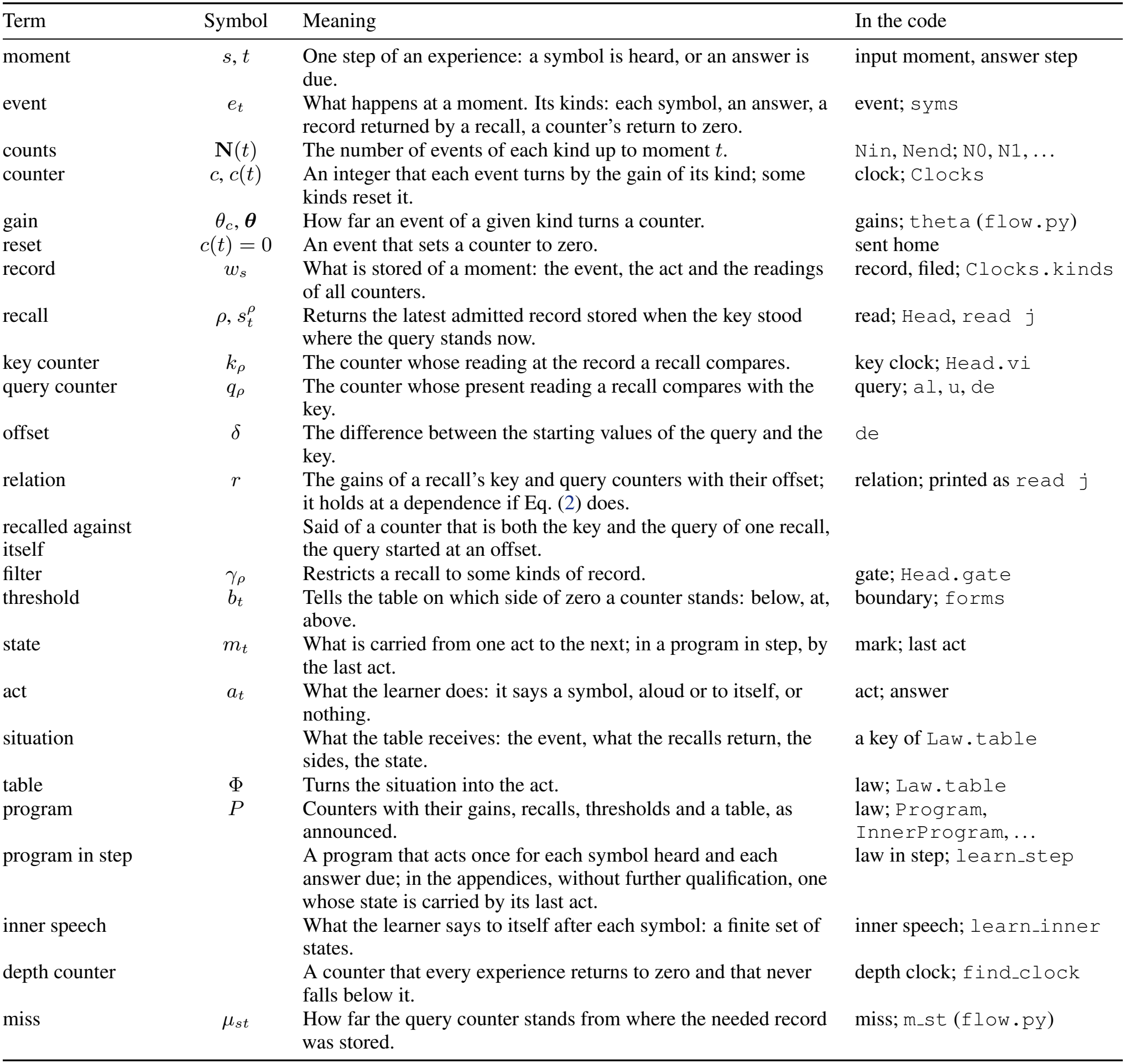
Terms of the memory and its programs. (Section 2).

| Term | Symbol | Meaning | In the code |
|---|---|---|---|
| moment | s, t | One step of an experience: a symbol is heard, or an answer is due. | input moment, answer step |
| event | e<sub>t</sub> | What happens at a moment. Its kinds: each symbol, an answer, a record returned by a recall, a counter's return to zero. | event; syms |
| counts | **N**(t) | The number of events of each kind up to moment t. | Nin, Nend; N0, N1, ... |
| counter | c, c(t) | An integer that each event turns by the gain of its kind; some kinds reset it. | clock; Clocks |
| gain | θ<sub>c</sub>, **θ** | How far an event of a given kind turns a counter. | gains; theta (flow.py) |
| reset | c(t) = 0 | An event that sets a counter to zero. | sent home |
| record | w<sub>s</sub> | What is stored of a moment: the event, the act and the readings of all counters. | record, filed; Clocks.kinds |
| recall | ρ, s<sub>t</sub><sup>ρ</sup> | Returns the latest admitted record stored when the key stood where the query stands now. | read; Head, read j |
| key counter | k<sub>ρ</sub> | The counter whose reading at the record a recall compares. | key clock; Head.vi |
| query counter | q<sub>ρ</sub> | The counter whose present reading a recall compares with the key. | query; al, u, de |
| offset | δ | The difference between the starting values of the query and the key. | de |
| relation | r | The gains of a recall's key and query counters with their offset; it holds at a dependence if Eq. (2) does. | relation; printed as read j |
| recalled against itself |  | Said of a counter that is both the key and the query of one recall, the query started at an offset. |  |
| filter | γ<sub>ρ</sub> | Restricts a recall to some kinds of record. | gate; Head.gate |
| threshold | b<sub>t</sub> | Tells the table on which side of zero a counter stands: below, at, above. | boundary; forms |
| state | m<sub>t</sub> | What is carried from one act to the next; in a program in step, by the last act. | mark; last act |
| act | a<sub>t</sub> | What the learner does: it says a symbol, aloud or to itself, or nothing. | act; answer |
| situation |  | What the table receives: the event, what the recalls return, the sides, the state. | a key of Law.table |
| table | Φ | Turns the situation into the act. | law; Law.table |
| program | P | Counters with their gains, recalls, thresholds and a table, as announced. | law; Program, InnerProgram, ... |
| program in step |  | A program that acts once for each symbol heard and each answer due; in the appendices, without further qualification, one whose state is carried by its last act. | law in step; learn_step |
| inner speech |  | What the learner says to itself after each symbol: a finite set of states. | inner speech; learn_inner |
| depth counter |  | A counter that every experience returns to zero and that never falls below it. | depth clock; find_clock |
| miss | μ<sub>st</sub> | How far the query counter stands from where the needed record was stored. | miss; m_st (flow.py) |

**Table 4.**
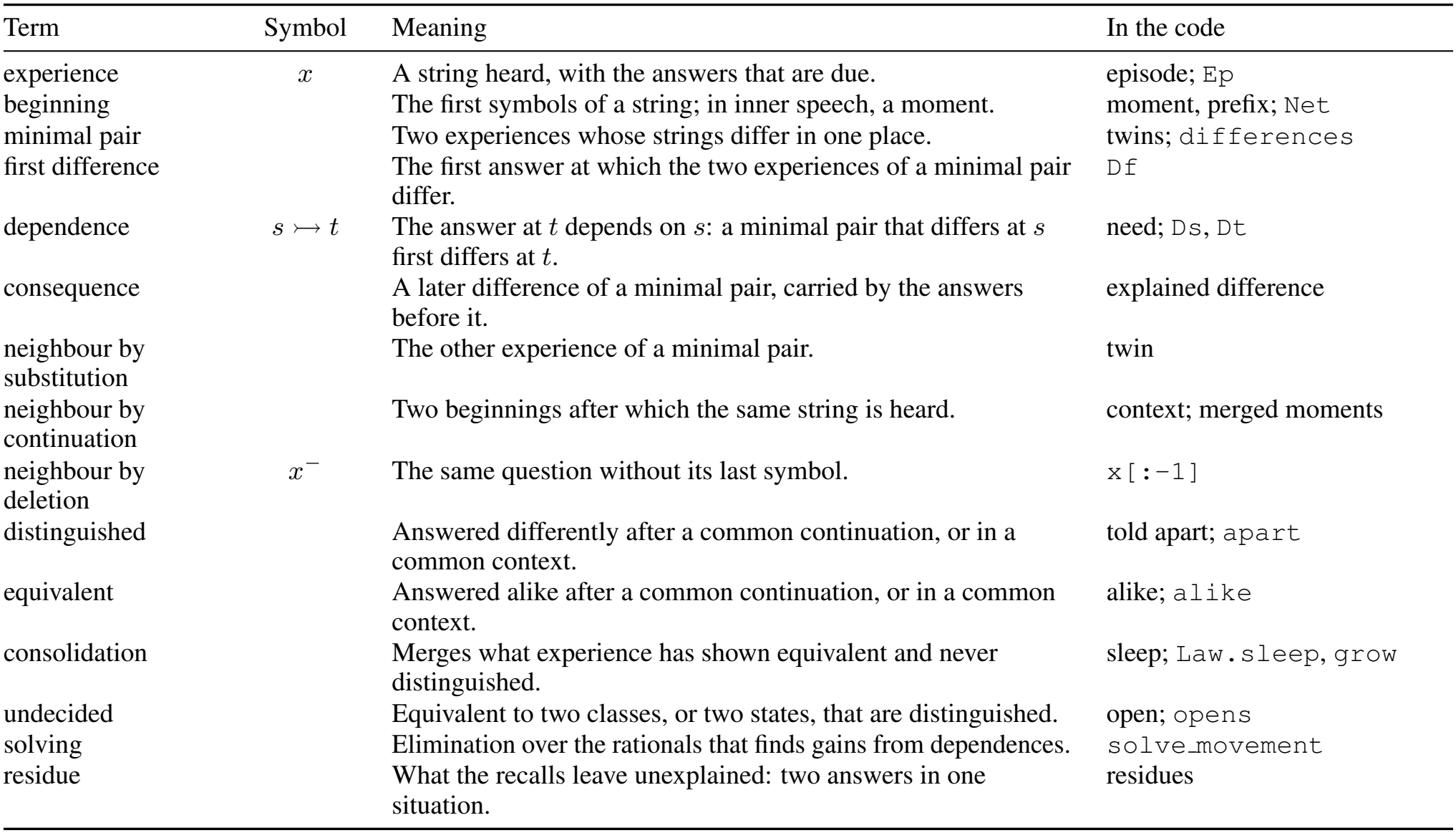
Terms of comparison. (Section 3).

**Table 5.** Terms of the outcomes and of experience. (Sections 3 and 4).

| Term | Symbol | Meaning | In the code |
|---|---|---|---|
| test on the longest strings | *L*, *m* | A program is announced only if it answers the *m* longest strings, of length *L*. | prediction; predicted |
| exact |  | A run whose program answers every test string correctly. | verdict 'exact' |
| says that it does not know |  | The program meets a situation that is not in its table, and says so. A program that does so on some test string and errs on none *abstains*. | silent; 'silent' |
| false program |  | A program that answers some test string wrongly, without warning. | false law; 'false' |
| no program |  | The learner announces nothing, and gives the reason. | no law; log['why'] |
| solved |  | A task is solved if each of ten seeds gives an exact program. | all_exact |
| question |  | A string that can be heard, with the answers due; asking is choosing one and being told its answers. | play; ask, asked |
| composition | *P* | Running a learned program and learning from what it answers. | thinking; learn_thinking |
| trial |  | Trying each candidate answer in the place of the unknown. | trial; learn_trial |
| recursion | *D*(*x*), ⊕ | Adding the differences between the answers to successive beginnings. | recursion; learn_recursion |
| library |  | The programs already learned, which composition and recursion use. | library, Library |
| experience of size *N* | *N* | 64 random strings of each length from 1 to *N*, repeated strings removed. | episodes(task, N, 64, seed) |
| standard experience |  | The experience of size 12: a median of 457 distinct strings (363 to 653). | STANDARD = (12, 64) |
| seed |  | Decides which strings are heard. | seed |
| learner |  | The memory of Eq. (1) with learning by comparison. | difference machine; dm.learn |

#### A.2. A minimal pair and a dependence

Reversal answers a string with its symbols in the opposite order. A string of *n* symbols is heard at the moments 1 to *n*, and its answers are due at the moments *n* + 1 to 2*n*. The learner remembers two experiences:

~~~
heard 10110, answered 01101;
heard 11110, answered 01111.
~~~

The strings differ in the second symbol and nowhere else, so the two experiences form a minimal pair. Their answers agree in the first three places and differ in the fourth, 0 against 1. Up to that answer the two experiences differ at moment 2 alone: the other four symbols heard are the same, and so are the three answers said before. Whatever procedure gave these answers, something of moment 2 reached the fourth one. The fourth answer, at moment 5 + 4 = 9, depends on moment 2: 2 ↣ 9.

The pair gives this one dependence, and other pairs give the others. Changing symbol *i* of a string of *n* symbols changes answer *n* + 1*− i* and no other, so the minimal pairs of reversal show that answer *j* depends on symbol *n* + 1 *−j*. Pairs are common in short experience: each of the 32 strings of five symbols has five neighbours by substitution, which makes 32 *·* 5*/*2 = 80 minimal pairs, the number 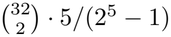 of Theorem 1(iii).

#### A.3. Reversal as a program

The dependences of reversal are served by one counter *c*, turned +1 by each symbol heard and *−*1 by each answer, and one recall that uses *c* as its key counter and *c* + 1, the same counter started one reading higher, as its query counter. The recall admits the records of symbols heard, and the table says the symbol recalled. On the string 110, whose reversal is 011:

| moment | event | reading of *c* | record stored at | query asks for | record returned |
|---|---|---|---|---|---|
| 1 | hears 1 | 1 | 1 |  |  |
| 2 | hears 1 | 2 | 2 |  |  |
| 3 | hears 0 | 3 | 3 |  |  |
| 4 | answer 1 | 2 |  | 3 | moment 3: says 0 |
| 5 | answer 2 | 1 |  | 2 | moment 2: says 1 |
| 6 | answer 3 | 0 |  | 1 | moment 1: says 1 |

The records of the symbols are stored at the readings 1, 2, 3 and recalled at 3, 2, 1. Every moment is stored, answers included (Eq. (1)), and the filter keeps the record of the first answer, stored at reading 2, from hiding the record of moment 2. Nothing in the program mentions the length of the string: the same counter, recall and table reverse a string of a million symbols, and Theorem 2(iv) shows that this is the solution that the dependences of reversal allow.

The learner prints this recall as I(s) = I -A +1 (Appendix C.6). The key counter I counts the symbols heard. The right side, read as a counter, starts at 1 and is turned +1 by each symbol and *−*1 by each answer, the present one included; it is the query counter *c* + 1 (Lemma 2).

#### A.4. The counter of the stack, solved from dependences

##### The task

The stack task of the benchmark hears a stack of the digits 0 and 1, bottom first, and then actions: 2 pops, 3 pushes a 0, 4 pushes a 1. Its answers are the final stack read from the top, followed by the end symbol 2. There are five kinds of symbol: two that fill the initial stack, a pop and two pushes.

##### Dependences

Take the experience 013424: the stack 01, then a push of 0, a push of 1, a pop and a push of 1. The final stack is 0101, bottom first, so the answers are 1, 0, 1, 0 and then 2. The minimal pairs that exchange a symbol for another of its kind (0 for 1, one push for the other) show four dependences: the first answer depends on symbol 6, the second on symbol 3, the third on symbol 2 and the fourth on symbol 1. A change of symbol 4 changes no answer, since its push is popped. A change of kind, a pop for a push, moves the depth of every later record, and the dependences it shows belong to other recalls; they make up the share of the pairs of dependences drawn in grey in Figure 2a.

##### Equations

Let **N**(*s*) count the symbols of each kind among the first *s*, and let ***θ*** = (*θ*_0_, *θ*_1_, *θ*_2_, *θ*_3_, *θ*_4_) be the gains of a key counter for the symbols 0 to 4. Let an answer depend on the record of moment *s*, and the answer *d* places later on the record of moment *s*^*′*^. A recall whose query falls by one at each answer serves both only if the key counter stands *d* readings lower at *s*^*′*^ than at *s*:

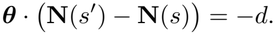

Each pair of dependences of the experience gives one such equation:

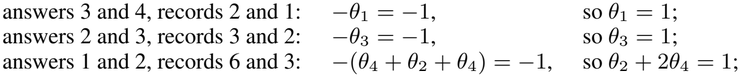

and the pairs two answers apart give sums of these. Two short experiences fix the rest. In 103 (the stack 10 and a push of 0, answered 0, 0, 1, 2) the second and third answers depend on symbols 2 and 1, which gives *−θ*_0_ = *−*1. In 044 (the stack 0 and two pushes of 1, answered 1, 1, 0, 2) the first and second answers depend on symbols 3 and 2, which gives *−θ*_4_ = *−*1, and so *θ*_2_ = 1*−* 2*θ*_4_ = *−*1. The solution is ***θ*** = (+1, +1, *−*1, +1, +1): each digit of the initial stack and each push raises the counter by one, and a pop lowers it by one. It is the depth of the stack. From the standard experience the learner solves all such equations together, 172 of them, of which 26% come from dependences of other recalls, and keeps the solution that more than half of them agree in (Theorem 2(ii)).

##### The recall

The learner prints its recall for the stack as N[+0+1-2+3+4](s) = N[+0+1-2+3+4](end) -A +1, with a filter that admits the symbols 0, 1, 3 and 4. The key counter is the depth, and the query is the depth at the end of the string, started one higher and lowered by one at each answer. On 013424 the depth after each symbol is 1, 2, 3, 4, 3, 4, and at the five answers the query asks for 4, 3, 2, 1, 0. Two records stand at depth 4, those of moments 4 and 6, and the latest, moment 6, is returned. Two stand at depth 3: moment 3, the push of 0, and moment 5, the pop, which stands at the depth of the record it uncovers. The filter is closed to pops, so moment 3 is returned. Moments 2 and 1 follow; at depth 0 nothing is found, and the table says the end symbol. The table has three situations, 0 recalled, 1 recalled and nothing recalled: consolidation has merged the digit 0 with the push of 0, and the digit 1 with the push of 1, which experience showed equivalent and never distinguished.

#### A.5. The states of parity

Parity hears a string of 0s and 1s and answers once, at the end, the number of 1s modulo two. A single answer at the end carries no state from one symbol to the next, so the learner has to say something to itself after each symbol, and what it says is fixed by which beginnings their continuations distinguish (Section 3.4).

The beginnings 1 and 111 are equivalent on every continuation *w*: the strings 1*w* and 111*w* differ by two 1s and have the same answer. The beginnings 1 and 11 are distinguished by the empty continuation, which they answer 1 and 0. Every beginning is equivalent either to the empty beginning, if it holds an even number of 1s, or to 1, if it holds an odd number. Two states are therefore enough, and one is too few: these are the classes of Myhill and Nerode for parity.

The learner sees only the continuations it has heard. The experiences 1 and 111, both answered 1, show the two beginnings equivalent on the empty continuation, the experiences 10 and 1110 show them equivalent on 0, and no experience distinguishes them; consolidation merges them. From the standard experience the learner prints

~~~
state 0 after (start) answer 0 steps 0>0 1>1
state 1 after 1 answer 1 steps 0>1 1>0
~~~

A 1 moves the learner from one state to the other, and a 0 leaves it where it is. Each step of this table was taken in at least 300 of the 461 experiences, so that experience decides each of them. Without inner speech parity has no program: one answer comes at the end, so no last act carries a state, and a threshold tells the side of zero of a counter, which says nothing of the parity of the count of 1s. Several thresholds on one counter at different offsets would read the counter in pieces, and the learner keeps at most one threshold of a counter (Appendix C.2).

#### A.6. Binary addition: two recalls, a threshold and a state

##### The task

Binary addition, as the benchmark gives it, hears the first number, the symbol 2 and the second number, each written lowest digit first. Each number is at least one and may carry leading zeros, which come last. The answers are the digits of the sum, lowest first and without leading zeros, followed by the end symbol 2. For 3 + 5 the learner hears 11 2 101, six symbols, and answers 0, 0, 0, 1 and then 2, since 8 is 1000 in binary.

##### The program

The learner’s program has two recalls, one threshold and a state carried by the last act.

- Recall 0 has the key counter I, the symbols heard, and the query counter A, the answers given, the present one included. Its filter admits the records stored before any 2 was heard. At answer *j* it returns symbol *j* while that lies in the first number, then the separator 2 once, and then nothing.
- Recall 1 has the key counter S|2, the symbols heard since the last 2, which the 2 resets, and the query counter A. Its filter admits the digits heard after the 2. At answer *j* it returns digit *j* of the second number.
- The threshold counter is the number of 1s heard less the number of 1s that the two recalls have returned so far. It stands at zero when nothing is left to add.
- The state is carried by the last act: the last answer and the two digits recalled at the last step. The digit said is the sum of the two digits recalled and of the carry received, modulo two, so the last act fixes the carry received and with it the carry passed on. Consolidation sorts the 23 last acts that experience holds into three states, the beginning, carry and no carry, and leaves 3 of them undecided.

On 11 2 101:

| answer | recall 0 | recall 1 | 1s heard less 1s recalled | state | says |
|---|---|---|---|---|---|
| 1 | 1 | 1 | 4 − 2 = 2 | beginning | 0 |
| 2 | 1 | 0 | 4 − 3 = 1 | carry | 0 |
| 3 | 2 | 1 | 4 − 4 = 0 | carry | 0 |
| 4 | nothing | nothing | 0 | carry | 1 |
| 5 | nothing | nothing | 0 | no carry | 2, the end |

At the fourth answer both recalls return nothing and the threshold is at zero, but the state carries a one, which is said. At the fifth nothing is carried and nothing is left to add, and the sum ends.

##### The threshold is a residue

The experiences 1001 2 10 and 1000 2 10, that is 9 + 1 and 1 + 1 with the first number written with three leading zeros, form a minimal pair. Their answers, 0101 and 01, each followed by 2, first differ at the third answer, which therefore depends on symbol 4. No recall returns symbol 4 there. At the third answer the recalls return 0 and nothing in both experiences, and the state is no carry in both, so the two moments are in one situation with two answers, a 0 and the end symbol. They differ in a count: symbol 4 is a *−*1 in the first experience, so one 1 heard is still to be recalled, and the threshold counter stands at 1 against 0. At every moment at which a sum ends, the 1s heard equal the 1s recalled, and elimination on the counts of events at those moments returns the gains of the threshold counter: +1 for a 1 heard, and 1 for each 1 that a recall returns (Theorem 3(i)). The third answer depends on symbol 4 through a counter whose side of zero it reads, the second of the three ways of Lemma 1.

#### A.7. A question in arithmetic modulo five

##### The task

Arithmetic modulo five hears a string of odd length: the digits 0 to 4 at the odd places, and the signs +,*−* and *×*, heard as the symbols 5, 6 and 7, at the even places. Multiplication goes before addition and subtraction, and the single answer, at the end, is the value modulo five.

##### The states

After a digit the program has to keep two numbers modulo five: the sum *s* of the terms already finished, and the product in progress *p*, with its sign. The value is *s* + *p*. After a sign it keeps a pair (*s, p*) that the next digit *d* turns into (*s, pd*): after *×* this is the pair it had, after + it is (*s* + *p*, 1), and after *−*it is (*s* + *p*, 4), since *−d ≡*4*d*. There are thus 25 states after a digit and 25 after a sign, the 50 states of the program that the learner writes. Its table has to tell each of them from every other, and this takes 5,843 strings when the learner may ask and 11,135 when it may not (Experiment 4).

##### The undecided beginning

In one experience (seed 8, 512 strings drawn at each length from 1 to 16, 5,834 distinct strings) nothing heard continues the beginning 1+1 *×*2 *×*2 with a product. After it the learner has heard only continuations that begin with + or *−*, and the end of the string; after these only the value of the beginning matters, and it is 1 + 4 0. The beginning is therefore shown equivalent to each state of value zero: the states after 0, 1+4, 2+3, 0 *−*2+2 and 4+1, whose pairs (*s, p*) are (0, 0), (1, 4), (2, 3), (3, 2) and (4, 1). Continuations that multiply distinguish these five from one another, so the beginning is undecided. Its true pair is (1, 4), and experience has not shown it.

##### The question

The learner looks for a remembered experience in which one candidate stood and which goes on from there to end differently for another; one such experience goes on with *×*2. It puts the undecided beginning in that place and asks for 1+1*×*2*×*2*×*2. Under the five candidates this question would be answered

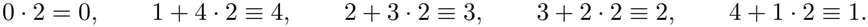

The world answers 1 + 1*·* 2*·* 2*·* 2 = 9 *≡*4, which leaves one candidate, the state after 1+4. One question settles five candidates here because one continuation gives each of them a different answer; Theorem 3(iii) bounds the number of questions by four. Made to guess, the learner merges the beginning with one of the five, and four of the five choices answer this question wrongly (Appendix D.3).

#### A.8. Multiplication by differences

##### The task

Binary multiplication, as the benchmark gives it, hears the multiplicand, the symbol 2 and the multiplier, each lowest digit first, and answers the product lowest digit first, followed by 2. For 13 *×*5 the learner hears 1011 2 101: 13 is 1101 in binary and is heard as 1011, and 5 is 101 either way.

##### Beginnings and differences

The beginnings of this question that are themselves questions are 1011 2 1, 1011 2 10 and 1011 2 101: 13*×* 1, 13*×* 1 (the multiplier 10, lowest digit first, is one) and 13*×* 5. The beginnings 101, 1011 and 1011 2 hold no multiplier and are no questions of the task; their answer is *nothing*, the end symbol alone. (The strings of the two shortest lengths, here 1 and 10, the learner leaves to memory: Appendix C.4.) The difference between the answer to a beginning and the answer to the beginning one symbol shorter is taken by trial with the learner’s own addition: it is the string *d* for which the addition, asked for the earlier answer followed by *d*, gives the later answer (Theorem 5(iii)). From nothing, the difference is the answer itself; between equal answers it is nothing.

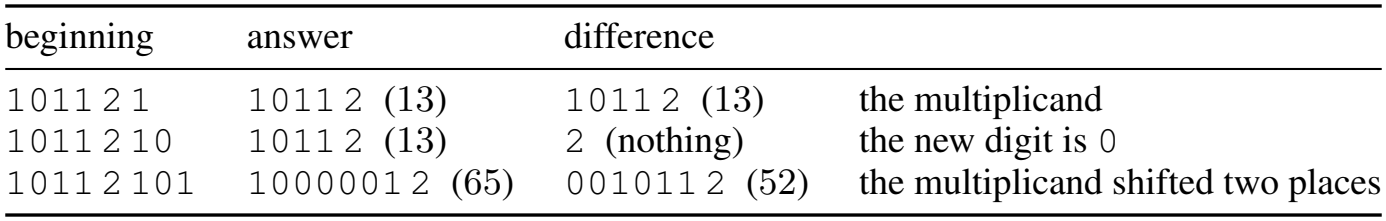

The differences 13, 0 and 52 are 13 *·*1, nothing and 13 *·*4: the multiplicand shifted to the place of the new digit of the multiplier when that digit is 1. In the benchmark’s format the shift puts zeros in front, and 52 is 001011. Figure 2d follows the same steps for 7 *×* 5.

##### The program

The differences have a program in step of 11 situations (Appendix C.6), and the answer to a question is the sum of the differences along its way, added by the learned addition. Here the addition is asked 1011 2 001011 and answers 1000001 2, which is 65; the end symbol of one answer serves as the separator of the next question to the addition. Each beginning of a question of *n* symbols costs one run of the program of the differences and at most one addition, each of order *n* steps, so a product takes a number of steps of order *n*^2^ (Theorem 5(iii)); the learned multiplication needs a number of moments that grows as the power 2.02 of the length (Experiment 6). To find the program of the differences the learner needs the answers to the beginnings of its questions, their neighbours by deletion, which passive experience seldom holds; from the standard experience it asks a median of 990 questions for them.

### B. Proofs

The main text gives each theorem in a short form. This appendix states each one in full, with all its conditions, and proves it: the five theorems of Section 3, the local rule of Section 5.3, and the lemmas and propositions on which they rest. A reader who wants to know exactly when a claim of the main text holds, or to check it, will find it here. Appendix B.1 sets out the memory of Eq. (1) and the notation, and Appendix B.2 shows how a difference between two experiences travels through a program. The theorems then follow in the order of the main text, each with its number there, its intuition in a few sentences, its proof and, where it applies, the way the learner uses it. Table 6 lists the results with the places where they are stated and proved; Table 7 lists the symbols.

**Table 6.**
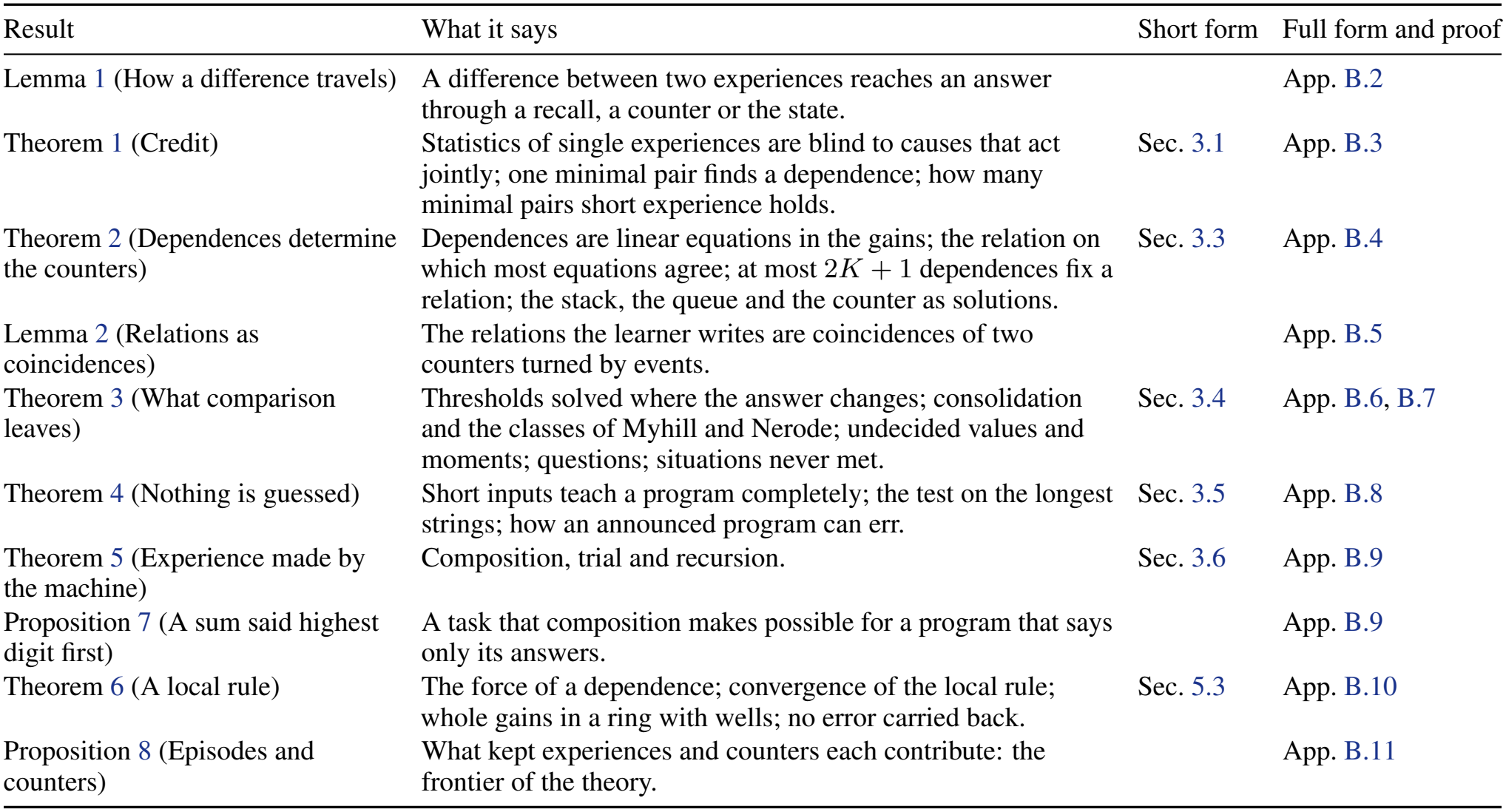
The results of the theory: where the main text states each in short form, and where it is stated in full and proved. Every entry is a link.

| Result | What it says | Short form | Full form and proof |
|---|---|---|---|
| Lemma 1 (How a difference travels) | A difference between two experiences reaches an answer through a recall, a counter or the state. |  | App. B.2 |
| Theorem 1 (Credit) | Statistics of single experiences are blind to causes that act jointly; one minimal pair finds a dependence; how many minimal pairs short experience holds. | Sec. 3.1 | App. B.3 |
| Theorem 2 (Dependences determine the counters) | Dependences are linear equations in the gains; the relation on which most equations agree; at most 2*K* + 1 dependences fix a relation; the stack, the queue and the counter as solutions. | Sec. 3.3 | App. B.4 |
| Lemma 2 (Relations as coincidences) | The relations the learner writes are coincidences of two counters turned by events. |  | App. B.5 |
| Theorem 3 (What comparison leaves) | Thresholds solved where the answer changes; consolidation and the classes of Myhill and Nerode; undecided values and moments; questions; situations never met. | Sec. 3.4 | App. B.6, B.7 |
| Theorem 4 (Nothing is guessed) | Short inputs teach a program completely; the test on the longest strings; how an announced program can err. | Sec. 3.5 | App. B.8 |
| Theorem 5 (Experience made by the machine) | Composition, trial and recursion. | Sec. 3.6 | App. B.9 |
| Proposition 7 (A sum said highest digit first) | A task that composition makes possible for a program that says only its answers. |  | App. B.9 |
| Theorem 6 (A local rule) | The force of a dependence; convergence of the local rule; whole gains in a ring with wells; no error carried back. | Sec. 5.3 | App. B.10 |
| Proposition 8 (Episodes and counters) | What kept experiences and counters each contribute: the frontier of the theory. |  | App. B.11 |

**Table 7.**
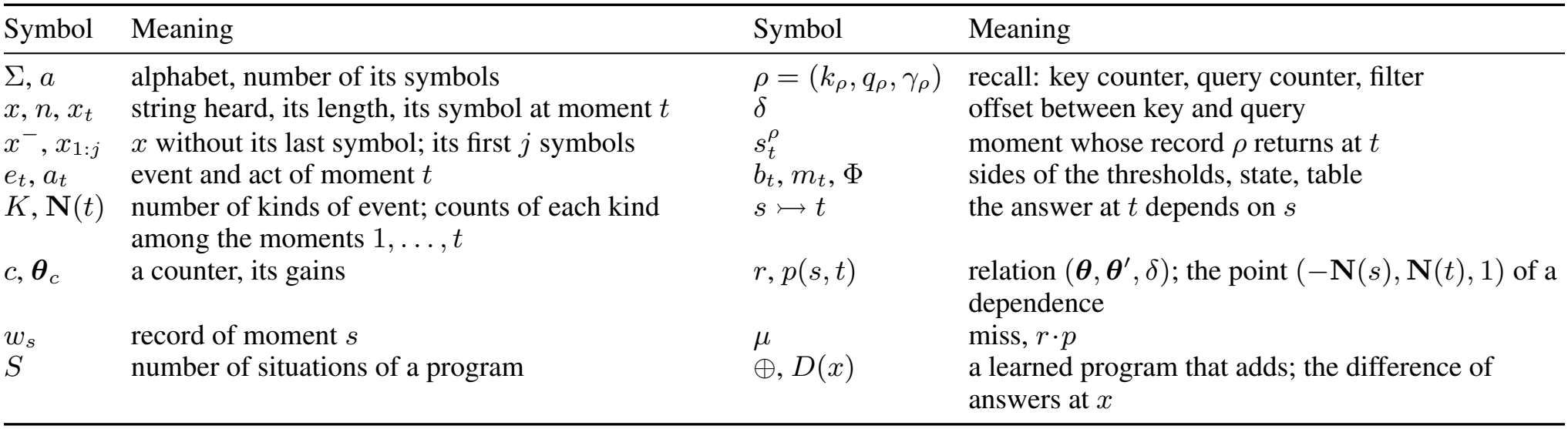
Symbols used in this appendix.

| Symbol | Meaning | Symbol | Meaning |
|---|---|---|---|
| Σ, a | alphabet, number of its symbols | ρ = (k<sub>ρ</sub>, q<sub>ρ</sub>, γ<sub>ρ</sub>) | recall: key counter, query counter, filter |
| x, n, x<sub>t</sub> | string heard, its length, its symbol at moment t | δ | offset between key and query |
| x<sup>−</sup>, x<sub>1:j</sub> | x without its last symbol; its first j symbols | s<sub>t</sub><sup>ρ</sup> | moment whose record ρ returns at t |
| e<sub>t</sub>, a<sub>t</sub> | event and act of moment t | b<sub>t</sub>, m<sub>t</sub>, Φ | sides of the thresholds, state, table |
| K, **N**(t) | number of kinds of event; counts of each kind among the moments 1, . . . , t | s ↣ t | the answer at t depends on s |
| c, **θ**<sub>c</sub> | a counter, its gains | r, p(s, t) | relation (**θ**, **θ**′, δ); the point (−**N**(s), **N**(t), 1) of a dependence |
| w<sub>s</sub> | record of moment s | μ | miss, r·p |
| S | number of situations of a program | ⊕, D(x) | a learned program that adds; the difference of answers at x |

**Table 8.** The constants and limits of the learner, with their names in the code (dm.PARAMS and dm/machine.py, dm/inner.py). Experiment 5 varies the four constants (Table 19).

|  | in the code | value | role |
|---|---|---|---|
| *Constants* |  |  |  |
| margin | margin | 20 bits | what is kept explains more than it costs to state, by at least this much |
| luck | luck | 4 standard deviations | a count exceeds chance if it exceeds it by this much |
| idle | idle | 0.10 | a kind of record needed in a smaller share of its returns may be closed out of a recall |
| major | major | 0.15 | a recall that meets this share of the differences is part of every table |
| *Limits of size* |  |  |  |
| recalls of a program | max_reads | 8 |  |
| thresholds offered | max_forms | 12 |  |
| recalls compared | top_relations | 40 | of the 160 best counted, in each round |
| dependences sampled | max_pairs | 3,000 | for each count of the queries |
| *Vocabulary and search* |  |  |  |
| gains solved for |  | up to 3 | also the gains of threshold forms |
| queries | ALS, DELTAS, anchors | α ∈ {0, ±1}, δ ∈ {−2, ..., 2} | at most two other constants, each ±1 or ±2 |
| solutions kept | KEEP | 3 | those that more than half of the pairs agree in |
| random draws | trials | 200 | when no gains solve every equation |
| questions for a moment | tries | 8 | before the learner turns to the next undecided moment |
| states of inner speech | max_states | a quarter of the strings, at least 16 | more states than the experience can bear: no program |
| time | time_budget | 600 s (120 for inner speech) | processor seconds of one core; 120 s in the light runs |

#### B.1. Setting and notation

##### Experiences and tasks

An experience of size *n* is a string *x∈* Σ^*n*^, heard one symbol at a moment, followed by the moments at which answers are due. A task assigns to every *x* its answers *y*_1_(*x*), …, *y*_*m*_(*x*), symbols of a finite set; *m* is a function of *n*, or the answers end with a symbol that says so, the *end symbol*. The moments are *t* = 1, …, *n* + *m*, and the event *e*_*t*_ of moment *t* is *x*_*t*_ for *t≤ n* and an *answer step*, at which an answer is said, for *t > n*. We write *a* = |Σ|, *x*^*−*^ for *x* without its last symbol, and *x*_1:*j*_ for its first *j* symbols; these are the *beginnings* of *x*.

##### Kinds of event and counters

The kinds of event are the symbols of Σand the answer step and, where a program uses them, the return of a record of a given content by a given recall (Appendix B.6), the return to zero of a ring of two states

(Lemma 2) and the things said by a program run first (Appendix B.9). *K* is their number, and **N**(*t*) *∈ ℕ* ^*K*^ counts the events of each kind among the moments 1, …, *t*. A *counter c* has a whole *gain θ*_*c*_(*k*) for each kind *k*, and possibly a kind of event that resets it to zero. Its reading is

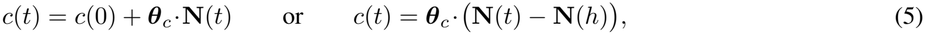

the second if *h ≤t* is the last moment that reset it. A counter that is not reset is thus a weighted count of the events so far, and this is why the equations of Theorem 2 are linear.

##### Records and recalls

The *record* of moment *s* is *w*_*s*_ = (*e*_*s*_, *a*_*s*_). It is stored with the readings of all counters at *s* and with its *kind*: the symbol and, for each symbol *b*, whether *b* had been heard before *s*, each of which is a coincidence in its own right. A *recall ρ* = (*k*_*ρ*_, *q*_*ρ*_, *γ*_*ρ*_) has a *key counter k*_*ρ*_, a *query counter q*_*ρ*_ and a *filter γ*_*ρ*_ on kinds of record. At moment *t* it returns the record of

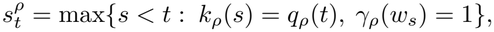

or nothing if no such moment exists. An offset *δ* between key and query, as in Eq. (2), is the reading at which the query counter starts; for a query counter that is reset, the offset is a fixed displacement in the comparison, *k*_*ρ*_(*s*) = *q*_*ρ*_(*t*) + *δ*, which a coincidence detector tests as easily as equality.

##### Thresholds, states and the table

A *threshold* is a counter of which the table is told the *side*: below zero, at zero or above. The *state m*_*t*_, from a finite set, is a function of what was recalled and done at *t*. The act is

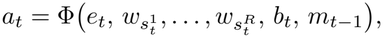

with Φ a finite *table* and *b*_*t*_ the sides of the thresholds. The argument of Φ is the *situation* of the moment. The table receives what the recalls return, or that they return nothing, the sides of the thresholds and the state, and no reading.

##### Programs

A *program* of Eq. (1) consists of counters with their gains, recalls, thresholds, a finite set of states and a table. It *answers* a task if at every answer step its act is the answer due. It *says only its answers* if it acts at answer steps alone, so that nothing has been said before the first answer. It runs *in step* with its input if it acts once for each symbol it hears, so that a string of *n* symbols with *m* answers takes *n* + *m* moments; a memory of this kind computes in step whatever a multitape

Turing machine computes in step (He & Wu, 2026a). The learner writes three kinds of program. Two run in step: programs whose state is carried by the last act, that is, by the last answer and what the recalls returned at the last step, and programs with *inner speech*, whose state is what the learner says to itself after each symbol. The third are programs composed of programs already learned (Appendix B.9). In the appendices, a program in step without further qualification is of the first kind, the kind the code learns in learn step; a program with inner speech is named as such.

##### Minimal pairs and dependences

Two experiences *x, x*^*′*^ *∈* Σ^*n*^ form a *minimal pair* at *s* if they differ at the moment *s* and nowhere else. In *x* the answer at *t depends* on *s*, written *s* ↣ *t*, if *x* forms a minimal pair at *s* with an *x*^*′*^ whose answers agree with those of *x* before *t* and differ at *t* (an answer that has ended counts as a symbol of its own); *t* is the *first difference* of the pair. A dependence is a property of the task, and the learner finds those whose minimal pairs its experience holds. Later differences of the same pair are *consequences* if the answers just before them differ, and are treated as dependences otherwise (Appendix C.2). Two beginnings of experiences are neighbours by *continuation* when the same string is heard after both, and the neighbour by *deletion* of *x* is *x*^*−*^. Appendix B.6 defines when continuations distinguish two beginnings and when they show them equivalent.

#### B.2. How an earlier moment reaches an answer

An earlier moment can affect a later answer only through what the table receives. The lemma makes this precise for the two runs of a minimal pair.

##### Lemma 1

(How a difference travels). *Let a program answer a task, and let s* ↣ *t in x, shown by the minimal pair x, x*^*′*^. *In the runs of the program on x and on x*^*′*^ *there are moments s* = *s*_0_ *< s*_1_ *<· · · < s*_*k*_ = *t such that each s*_*i*+1_ *depends on s*_*i*_ *in one of three ways:*

a. content: *a recall at s*_*i*+1_ *returns the record of s*_*i*_ *in one of the two runs;*
b. count: *the event or the act of s*_*i*_, *or a record returned there, turned a counter or reset it, and at s*_*i*+1_ *a recall compares a reading of that counter or a threshold reports its side;*
c. state: *s*_*i*+1_ = *s*_*i*_ + 1, *and the state carried from s*_*i*_ *enters the table*.

##### Intuition

Before *s* the two runs are the same. After *s* the table sees the event, what the recalls return, the sides of the thresholds and the state, so a difference that reaches the answer at *t* has travelled along recalls, counters and the state, and along nothing else.

*Proof*. Let *D* be the set of moments at which the two runs differ in the event, in the act, in what a recall returns (the moment or its record) or in the side of a threshold. The runs agree before *s*, where the strings agree; *s∈ D*, because the events differ there, and *t∈ D*, because the acts do. Let *u ∈ D* with *u > s*. The event at *u* is the same in both runs, so the runs differ at *u* in what a recall returns, in the side of a threshold, or, failing both, in the act alone; and an act that differs while its event, returns and sides agree has a different state *m*_*u−*1_ among its arguments. In each case there is an earlier moment of *D* on which *u* depends.

If a recall at *u* returns the same moment *v* in both runs with different records, then *v ∈ D*, and *u* depends on *v* by content. If it returns different moments, or a moment in one run only, let *v*^*′*^ be the later of the moments returned. In the run that does not return *v*^*′*^, either the filter refuses *v*^*′*^, so that its record differs between the runs, *v*^*′*^ *∈ D*, and *u* depends on *v*^*′*^ by content; or the key reading at *v*^*′*^ or the query reading at *u* differs between the runs. A reading is a sum of gains over what happened since the last reset, and the event at *u* is the same in both runs, so some moment *v < u* at which an event, an act or a return differs, hence *v ∈ D*, turned or reset one of the two counters, and *u* depends on *v* by count. If a threshold reports different sides at *u*, the reading of its counter differs, and the same argument gives a moment *v ∈ D, v < u*, on which *u* depends by count, or a return at *u* that differs, which is the case already treated. If the states *m*_*u−*1_ differ, what was recalled or done at *u −* 1 differs, so *u −* 1 *∈ D*, and *u* depends on *u −* 1 by state.

Every moment of *D* after *s* thus depends on an earlier moment of *D*. Following these dependences back from *t* gives a decreasing chain in *D*, which can end only at a moment of *D* with no earlier one, and that moment is *s*.

##### The three ways and what the learner learns

The method of difference rests on this lemma: in a program whose table receives records, coincidences and a state, a difference between two experiences travels along recalls, counters and states. The three ways are the three things the learner learns. Content is a recall that returns a record, which minimal pairs find (Theorem 2). Count is a threshold, as in sorting, where each symbol moves the place at which the answer changes (Theorem 3(i)). State is what the last act or inner speech carries from one moment to the next (Theorem 3(ii)). The last two leave no record that a recall could return, and they show up as what the recalls leave unexplained.

#### B.3. Credit Theorem 1 **(Credit)**

i. *Let the answer at a step be y* = *r*_1_ + *· · ·* + *r*_*m*_ + *c* (mod *b*), *where r*_1_, …, *r*_*m*_ *are what m≥* 2 *recalls return, independent and uniform on {*0, …, *b−* 1 *}, and where c and what every other candidate recall returns are independent of each r*_*i*_ *given the others. Then for every set of candidate recalls that omits at least one of the m, what the set returns is independent of y, and has with y the same joint distribution as the returns of a set of useless recalls with the same marginals. No statistic of single experiences that involves fewer than all m recalls therefore distinguishes a useful recall from a useless one*.
ii. *Let x, x*^*′*^ *form a minimal pair at s with first difference t. For every procedure that maps what was heard and what was said so far to the next answer and gives the answers of x and x*^*′*^, *the answer at t is not a function of what was heard at the moments other than s and of what was said before t, however many moments lie between s and t. Conversely, if for some x the answer at t changes when the symbol at s is changed and nothing said before t changes, then s* ↣ *t in x, and the pair that shows it is x and the changed string*.
iii. *Two different strings drawn uniformly from* Σ^*n*^ *form a minimal pair* (*wit h probability n*(*a −* 1)*/*(*a*^*n*^ *−* 1), *so among N different strings so drawn the expected number of minimal pairs is* 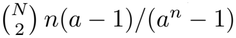.

##### Intuition

When an answer is the sum of several returns modulo some base, every incomplete set of them is independent of the answer, so a learner that counts how often a record and an answer go together sees nothing until it looks at all the causes at once. A minimal pair holds everything fixed but one place, and the first answer at which its two experiences part shows a dependence at once, whatever happened in between. Short experience is full of minimal pairs.

*Proof*. (i) Let the set omit *r*_*i*_, and let *Z* collect what the set returns together with *c* and the *r*_*j*_, *j* ≠ *i*. Then *y* = *r*_*i*_ + *g*(*Z*) (mod *b*) for a function *g*. Since *r*_*i*_ is independent of the *r*_*j*_, *j* ≠ *i*, and the rest of *Z* is independent of *r*_*i*_ given them, *r*_*i*_ is uniform and independent of *Z*, so *y* is uniform given *Z*: it is uniform and independent of *Z*, hence of what the set returns. Their joint distribution is the product of the marginals, which is also the joint distribution for recalls on which *y* does not depend. A statistic computed from the returns of fewer than all *m* recalls and from the answers, over any number of experiences taken one at a time, therefore has the same distribution whether its recalls are useful or not. The first sets of recalls that such statistics can tell apart contain all *m*, and among *R* candidates there are 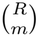 sets of that size.

(ii) What was heard at the moments other than *s* is the same in *x* and *x*^*′*^, and so is what was said before *t*, by the choice of *t*. A function of these alone gives the same answer at *t* in both, while the answers at *t* differ. The number of moments between *s* and *t* does not enter. The converse is the definition of a dependence.
(iii) Two different strings of Σ^*n*^ drawn uniformly form a minimal pair with pr(ob ability *n*(*a −* 1)*/*(*a*^*n*^ *−* 1): of the *a*^*n*^ *–* 1 strings other than a given one, *n*(*a −* 1) differ from it in one place. There are 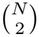 pairs of strings.

##### Worked cases

In binary addition the answer at step *k* is 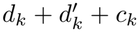 (mod 2), where *d*_*k*_ and 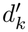 are the two digits of column *k* and *c*_*k*_ is the carry into it, a function of the lower columns. The assumptions of part (i) hold with *m* = 2, and they hold for the sum of two numbers in any base, for a difference and for a bitwise exclusive or. For part (iii), the 32 strings of five bits, all of them, give 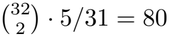 minimal pairs, the number in Section 3.1.

##### Why the first difference

Part (ii) asks nothing about the procedure and nothing about the other causes of the answer, and this matters twice. First, a cause may act only together with others. In the conjunction of two strings each digit matters only where the other is one, and in a product a digit of the multiplicand matters only if the digit of the multiplier is one. A test that asked whether a change of the record always changes the answer would reject such recalls; the definition asks for one minimal pair, and minimal pairs whose answers agree count against nothing. Second, a difference between the two experiences of a minimal pair need not be a dependence. Changing a low digit of a sum can change every higher digit of the answer, through the carry. These later differences are consequences, carried by the program’s own acts, and a learner that took them for dependences would look for recalls that do not exist. The first difference is the one place at which the two experiences are certain to differ in nothing but the place that was changed. The learner applies the same reasoning to the later differences (Appendix C.2): a later difference is explained if a recall already found returns different records in the two experiences at that step or at the step before, or if the answers just before it differ, and only what is left unexplained is a dependence.

##### Returns on which nothing depends

A relation is judged by its dependences and also by what it returns when nothing depends on the record. Of the returns of a candidate recall, some are records that a minimal pair changes. The share of these whose change makes a difference at the step of the return says how much of what the recall does is of use, and the information that “this record is returned at this step” carries about “changing it makes a difference there” is counted in bits (Appendix C.2). This is Mill’s joint method of agreement and difference: the relation holds where the phenomenon is present and fails where it is absent. A kind of record on which the answer depends in fewer than a tenth of its returns, and less often, by four standard deviations, than it would if the answer depended on it as often as on the other kinds, is excluded by the filter of the recall.

#### B.4. Dependences determine the counters

For counters that are not reset between the two moments, a *relation r* = (***θ, θ***^*′*^, *δ*) *∈* ℤ ^2*K*+1^, made of the gains of a key counter, the gains of a query counter and an offset, *holds* at a pair of moments (*s, t*) if

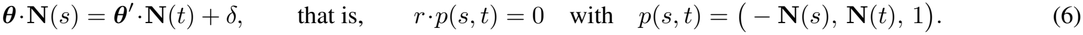

This is Eq. (2). For a counter that is reset, **N** is counted from the last reset, and what follows is unchanged. A relation that holds at a dependence *s* ↣ *t serves* it if no later record that passes the filter has the reading of *s*.

**Theorem 2 (Dependences determine the counters)**

i. *The relations that hold at every dependence of a set F are the whole points of the orthogonal complement of the span of {p*(*s, t*) : (*s, t*) *∈ F}. For two dependences* (*s, t*) *and* (*s*^*′*^, *t*^*′*^) *of one experience the condition* ***θ*** *·*(**N**(*s*^*′*^) *−* **N**(*s*)) = ***θ***^*′*^ *·*(**N**(*t*^*′*^) *−* **N**(*t*)) *does not contain δ. A basis with whole entries of the solutions is computed by elimination over the rationals in O*(|*F*|*K*^2^) *operations, whereas the counters with gains in {−g*, …, *g} number* (2*g* + 1)^*K*^.
ii. *Let ν such equations in d* + 1 *unknowns be given, and let a share π of them be met by a relation r*^*\**^, *any d different ones of which determine r*^*\**^ *up to scale. Then d different equations drawn at random have r*^*\**^ *as their solution with probability* Π_*i<d*_(*πν − i*)*/*(*ν − i*), *close to π*^*d*^ *when ν is large compared with d. If* 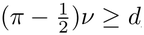, every relation that more than half of the equations meet is *r*^*\**^ *up to scale, whatever the other equations are*.
iii. *Let F*_*n*_ *be the dependences of a task, served by one recall, in experiences of size at most n. There are a size n*_0_ *and a set of at most* 2*K* + 1 *dependences in* 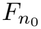 *such that every relation that holds at them holds at every dependence of F*_*n*_, *for every n*.
iv. *Let n symbols be followed by answers, let answer j depend on one symbol, and let the filter admit the records of symbols heard. If answer j depends on symbol n* + 1*− j (nested dependences), the dependences are served by the counter with gain* +1 *for every symbol and −*1 *for an answer step, recalled against itself with offset* +1: *a stack. If it depends on symbol j (serial dependences), no counter that is not reset serves the dependences when recalled against itself; they are served by a counter with gain* +1 *for an answer step recalled against a counter with gain* +1 *for every symbol, a queue, and by one counter with these two gains that the first answer step resets, recalled against itself with offset* +1. *If the answer changes at the step at which one count of events reaches another, as when the number of answers reaches the number of symbols of some kinds, the change is at a coincidence of two counters, each turned* +1 *by the events it counts; their difference is the counter of computer science, a register that events raise and lower and that is tested for zero*.

##### Intuition

A dependence says that a counter stood at the record where another stands at the answer. A counter that is not reset is a weighted count of the events so far, so the dependence is one linear equation in the gains, and the dependences of a recall form a linear system that elimination solves. The solution mentions no length, so the dependences of short strings fix it for every length, and the stack, the queue and the counter of computer science are the solutions for nested dependences, for serial ones, and for an answer that changes when one count reaches another.

*Proof*. (i) The condition (6) is one linear equation in *r* for each dependence. Subtracting the equations of two dependences of one experience removes *δ*, which is common to them. The whole solutions of a homogeneous system with whole coefficients are rational combinations of the basis that Gaussian elimination over the rationals gives, cleared of denominators, and eliminating |*F* | equations in at most 2*K* + 1 unknowns takes *O*(|*F*|*K*^2^) operations.

(ii) If the *d* equations drawn are all met by *r*^*\**^, they determine it up to scale, and it is their solution; if one of them is not met by *r*^*\**^, *r*^*\**^ does not solve them. Of the 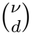 draws, 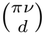 consist of equations met by *r*^*\**^, and their ratio is the product given. Now let a relation *r* be met by more than half of the equations. The equations met by both *r* and *r*^*\**^ number more than 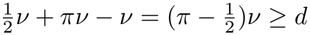. They are among the equations met by *r*^*\**^, so any *d* of them determine *r*^*\**^ up to scale, and *r* satisfies them: *r* is a multiple of *r*^*\**^. The other equations do not enter.
(iii) The spans of the sets *{p*(*s, t*) : (*s, t*) *∈ F*_*n*_ *}*increase with *n* and have dimension at most 2*K* + 1, so they are the same from some *n*_0_ on, and equal the span for the dependences of all sizes. Choose a basis of it among the points of 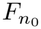. A relation orthogonal to the basis is orthogonal to the span, and so holds at every dependence of every size.
(iv) Let *c* have gain +1 for a symbol and *−*1 for an answer step. Symbol *i* is stored under *c* = *i*, and at answer *j* the counter shows *n j*, so *c*(*s*) = *c*(*t*) + 1 holds for *s* = *n* + 1 *j*, and no later record of a symbol has that reading. Now let answer *j* depend on symbol *j*, and let a counter that is not reset, with gains *θ*(*σ*) for the symbols and *β* for an answer step, serve the dependences against itself at offset *δ*. Then for every string and every *j ≤ n*, Σ _*i≤j*_ *θ*(*x*_*i*_) = Σ _*i≤n*_ *θ*(*x*_*i*_) + *βj* + *δ*, that is, *− Σ*_*j<i≤n*_ *θ*(*x*_*i*_) = *βj* + *δ*. The left side is independent of the symbols only if all *θ*(*σ*) are equal, to *θ* say; then *−θ*(*n− j*) = *βj* + *δ* for all *j≤ n* and all *n*, and *j* = *n* gives *β* = *δ* = 0, and then *θ* = 0. This counter does not turn: every record has the same reading, the latest symbol is returned at every answer, and only the last answer is served. The counter *A* of the answers shows *j* at answer *j*, and the counter *I* of the symbols stored symbol *j* under *j*, so *I*(*s*) = *A*(*t*), and no later record of a symbol has that reading. A counter that counts symbols and answers and is reset by the first answer step stores symbol *j* under *j* and shows *j−* 1 at answer *j*, which is offset +1. For the last claim, two counts of events are the readings of two counters, each turned +1 by the events it counts, and one count reaches the other at the step at which the readings coincide; the number of answers and the number of symbols of given kinds are the readings of the counter of answers and of a counter turned by those kinds. The difference of the two readings rises and falls with the events and is zero exactly at the coincidence. □

##### How the learner uses the theorem

The learner solves for the key counter first (Appendix C.2; solve movement in dm/machine.py). Two dependences of one experience whose answers are *d* steps apart ask that the key counter stand, at the two records, as far apart as the query moves in *d* answers. With the query’s gain for an answer fixed at *−*1, which sets the scale and the sign, this is one equation in the *K* gains of the key counter, whose coefficients are the numbers of symbols of each kind between the two records. The learner solves all these equations together; if no vector solves them all, it solves draws of *K−* 1 of them at random and keeps the solutions on which more than half of the equations agree (part (ii)). Where the equations met by *r*^*\**^ do not determine it, because a kind of event seldom lies between the two records of a pair of dependences, there are several such solutions, and all are kept for the dependences to choose among (Appendix C.2). For the stack of the benchmark, with two symbols that fill the initial stack, a pop and two pushes, the solution is (+1, +1, *−*1, +1, +1): a counter of depth (Figure 2a). Nothing about stacks was given. A list of the counters with gains up to one over five kinds of symbol would have 243 entries, and with gains up to three, 16,807.

Where the records on which successive answers depend lie a fixed number of symbols apart whatever the symbols are, the solution has equal gains: it is the counter *I* of all symbols, which the learner has from the start, together with the counters that a symbol resets. The query is then found by counting: among the queries turned by symbols, by every second symbol, by answers and by every second answer, with gains up to two, the learner takes the one at which the most dependences are served (each is tested by inner products, part (i); Lemma 2 says what these queries are). The sizes in part (iii) are small: the dependences that each recall of Appendix C.6 accounts for span their space on strings of at most two symbols for reversal and duplication, three for the stack, four for odds first and five for binary addition. Part (iv) derives the disciplines of the first paper from the kinds of dependence: a dependence on the latest unanswered symbol is served by a counter that the answers turn back, a dependence on the earliest by two counters or by one that is reset, and a dependence on a count by a threshold.

#### B.5. Relations as coincidences of counters

The learner writes a relation in the form

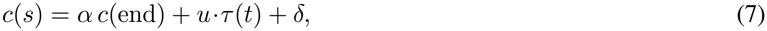

with *c* a key counter, *c*(end) its reading when the question ends, *α ∈{* 0, *±* 1 }, *δ∈ {−*2, …, 2 }, and *τ* = (*I, W, A, E, H*) the numbers of symbols heard, of symbols heard from the world (the others being what a program run first has said), of answers given, of pairs of symbols heard and of pairs of answers given; at most two entries of the whole vector *u* differ from zero, each *±*1 or *±*2 (Appendix C.1). The form is compact for counting. The lemma shows that each such relation is a coincidence between the key counter at the record and a query counter turned by events, so that the programs the learner writes run on coincidences, as Eq. (1) requires.

##### Lemma 2

(Relations as coincidences). *The terms of Eq*. (7) *and the forms of thresholds are readings of counters of Eq*. (1).

i. *I, W and A are counters of Eq*. (1) *with gain one for the events they count. E and H are counters turned by an event of their own: the return to zero of a ring of two states that every symbol, or every answer, turns. A counter “since the last b” is I reset by b*.
ii. *Let c be a counter that is not reset, with gains* ***θ***. *The relation* (7) *holds exactly when the key counter c coincides, at the record, with the query counter q that starts at δ and is turned by a symbol of kind σ by αθ*(*σ*) + *u*_*I*_, *and by u*_*W*_ *more if the symbol comes from the world, by every second symbol by u*_*E*_, *by an answer by u*_*A*_ *and by every second answer by u*_*H*_ . *For a counter c since the last b, the same holds with q reset by b and δ a displacement of the comparison when α≠* 0 *and u*_*I*_ = *u*_*W*_ = *u*_*E*_ = 0, *and with the gains above when α* = 0.
iii. *The side of a threshold form f* = Σ_*k*_ *ϕ*_*k*_*N*_*k*_ + *ϕ*_0_, *with N*_*k*_ *the numbers of events of each kind (symbols heard, answers given, records of each content returned by each recall), is the side of the counter with gains ϕ that starts at ϕ*_0_. *On a ring, where a reading has no sign, the side is kept by a latch that is set to zero when the counter reaches zero and to the sign of the turn when it leaves zero*.

*Every program that the learner writes with such recalls and thresholds is therefore a program of Eq. (1): its table receives records, sides and its state, and its counters are compared for equality alone*.

##### Intuition

Each term of the right side of Eq. (7) grows by a fixed amount at each event of some kind, so the right side is itself the reading of a counter, the query, and the relation is the coincidence of key and query. A sum of counts is likewise the reading of a counter, and its side of zero is a threshold.

*Proof*. (i) This is the definition of the counts: *E* = *⌊I/*2*⌋* increases exactly when the ring of two states returns to its first state, and likewise *H* = *⌊A/*2*⌋*.

i. During the question every term of the right side except those in *A* and *H* grows with the symbols: *αc* by *αθ*(*σ*) at a symbol *σ, u*_*I*_ *I* + *u*_*W*_ *W* by *u*_*I*_, or by *u*_*I*_ + *u*_*W*_ at a symbol from the world, and *u*_*E*_*E* by *u*_*E*_ at every second symbol. After the question only *A* and *H* change. The right side is therefore the reading of the counter *q* described, and the relation says *c*(*s*) = *q*(*t*). For a counter since the last *b* with *α* ≠ 0 and no term in *I, W* or *E*, the right side during the question is *α* times a counter that *b* resets, plus *δ*: the query reset by *b*, compared at the displacement *δ*. For *α* = 0 the right side does not involve *c*, and the query is the counter with the gains above.
ii. A sum of counts with whole coefficients is the reading of the counter with those gains, and it is at zero exactly when the form vanishes. The counter changes sign only by passing through zero or by a turn that carries it across zero, and the latch holds the sign as long as no turn carries the counter across zero in one step. This holds for the thresholds of the programs of this paper, and it is checked when a program is compiled into a circuit (Appendix D.8).

##### The one form outside the lemma

The form that part (ii) leaves out, a counter since the last *b* with *α* ≠ 0 and a term in *I, W* or *E*, is the sum of a counter that *b* resets and one that it does not. A query for it needs one operation beyond Eq. (1): a counter set, at an event, to where another counter stands. One recall of the benchmark has this form, one of the two recalls of the missing duplicate (Appendix C.6). The other programs in step have the form of the lemma as they stand, and they run as circuits of rings and place cells (Appendix D.8). What is linear in the learner is the way a counter adds up its turns: the learner solves linear equations in counts, and the programs it writes compare readings for equality and read sides of zero.

#### B.6. What comparison leaves

The recalls that minimal pairs find may leave moments in one situation with different answers. This is the place for Mill’s method of residues: what the known causes leave unexplained is the effect of the causes still unknown (Mill, 1843). To state what remains, fix the recalls and thresholds of a program *P* .

##### Views, values and classes

The *view* of a moment is its event together with what the recalls return there and the sides of the thresholds; with these fixed, the table and the state form a finite automaton that reads views and writes acts. A *continuation* of a moment is a sequence of views that follows it. A continuation that follows two moments *distinguishes* them if the acts it calls for after them differ at some step, and *shows them equivalent* if they agree. In an experience, two moments are *equivalent* if some continuation in the experience shows them equivalent and none distinguishes them, and a moment is *undecided* if it is equivalent to two moments that are distinguished. The same words apply to the *values* of one feature of the situation: what a recall returns, the side of a threshold, or a candidate for the state, which is a last act in a program in step and a moment in inner speech. The *context* of a value at a moment is the values of the other features there. A context in which two values both occur distinguishes them if their answers there differ, or if one of them has two answers there, and shows them equivalent if both have one and the same answer. *Consolidation* merges two values if they are equivalent and neither is undecided. Its classes are those that merged pairs connect, a class that would hold two distinguished values being left unmerged. It then compares again: with the classes of every feature at hand, a value that is equivalent to a class other than its own, while that class and its own are distinguished, is taken out of its class and merged with nothing, and so on until no value is taken out (Algorithm B.1).

**Theorem 3 (What comparison leaves)**. *Let a task have a program P of Eq*. (1).

i. *Besides the records its recalls return, the table of P receives whether a recall returns a record at all, the present event, the sides of its thresholds and the state. Two moments that the recalls leave in one situation with different answers therefore differ in the side of a threshold or in the state. Let a threshold of P have gains ϕ over the kinds of event and start ϕ*_0_, *and let T be a set of moments at which it is at zero. Then* (*ϕ, ϕ*_0_) *is orthogonal to the vectors* (**N**(*t*), 1), *t ∈T: it lies in their null space, which elimination computes. If the answer of the task changes exactly where the threshold reaches zero, the moments of change determine ϕ up to the null space of their counts*.
ii. *Every program with the recalls and thresholds of P gives different states to moments that are distinguished. If every continuation that can follow one moment can follow every other, not being distinguished is an equivalence, and its classes are the states of the program with the fewest states. If the state of P is a function of the last view and act, the classes are classes of values of the last view and act, and two such values lie in different classes exactly when some present view calls for different acts after them. In an experience, no class of consolidation holds two distinguished values; consolidation merges no undecided value, and when it stops comparing, no merged value is equivalent to a class other than its own that is distinguished from its own; the classes formed in one pass do not depend on the order in which pairs are taken; and from complete experience, in which every two values have met in every context, the classes are those of the equivalence, the classes of Myhill and Nerode*.
iii. *Let a value be undecided, and let it be given, in each context in which it was not met and in which every class it is equivalent to has one answer, the same for all, that answer, and no answer elsewhere, where the program says that it does not know. If the value belongs to one of those classes, every answer so given is right; and if a moment is one of several states that take the same step after every view, it takes those steps. Let a moment u be equivalent to each of k≥* 2 *states, any two of which are distinguished: an undecided moment. For any two of them a continuation w in the experience follows both and is answered differently, and the world’s answer to the string heard up to u followed by the symbols of w contradicts at least one of the two; so at most k−* 1 *such questions leave at most one candidate. A program in which u is identified with no state answers, after u, what experience has followed u with, and otherwise says that it does not know. Let experiences be drawn independently, and let S be the number of situations of a program. The probability that a new experience contains a situation that none of N earlier ones contained is at most S/*(*eN* ). *If moments are exchangeable, the probability that a further moment falls in a situation not met among n earlier ones is the expected number of situations met exactly once among n* + 1 *moments, divided by n* + 1.

##### Intuition

Once the recalls are found, what remains of the dependences is a count or a state (Lemma 1). A count changes the answer where a counter reaches zero, and the counts at the moments of change are equations for its gains. A state is fixed by which beginnings their continuations distinguish, as Myhill and Nerode showed for complete experience; from partial experience consolidation merges only what has been shown equivalent and never distinguished, compares again until nothing changes, and leaves the rest undecided, to be asked about or answered only where every candidate agrees.

*Proof*. (i) The argument of Φ holds nothing else. If the threshold is at zero at *t∈ T*, then *ϕ·* **N**(*t*) + *ϕ*_0_ = 0, one linear equation for each moment.

(ii) If two moments had the same state, the automaton would answer every continuation that follows both in the same way, so moments that are distinguished have different states. When continuations are shared, not being distinguished is reflexive, symmetric and transitive. Giving every class one state defines an automaton, because two moments of one class that receive the same view are followed by moments of one class (a continuation that distinguished the later two would, preceded by that view, distinguish the earlier two) and call for the same act. It has the fewest states, since every automaton has at least one state for each class. This is the theorem of Myhill (1957) and Nerode (1958). Now let the state be a function of the last view *ξ* and act. Two moments with the same last view and act have the same state and are never distinguished. For a table Φ whose state *χ* is the pair itself, the next state is the pair of the present view and of the act Φ(*ξ, χ*). So if Φ(*ξ, χ*) = Φ(*ξ, χ*^*′*^) for every *ξ*, the next states are equal and nothing later distinguishes the two; if not, one step does. Distinguishing by a continuation thus reduces to distinguishing in one step, which is what the learner counts.

In an experience, a class that would hold two distinguished values is left unmerged, so no class holds such a pair, and a value that is undecided is merged with nothing. Whether a pair is merged depends on the evidence alone, and the classes are the connected components of the merged pairs, so no order enters. Comparing again only takes values out of classes, at least one in each round that changes anything, so it stops; and when it stops, its condition holds for no value, which is the second claim. If every two values have met in every context of the experience, every pair is either distinguished or equivalent. Being equivalent is then the equivalence above, so no value is equivalent to two values that are distinguished: nothing is undecided, comparing again takes nothing out, and consolidation merges exactly the classes of the equivalence.

(iii) An answer on which all the classes agree is the answer of whichever of them the value belongs to, and a step that all the states take is the step of whichever of them the moment is; nothing that experience left undecided enters either. Two states that are distinguished have, by definition, a continuation *w* that follows both in the experience and is answered differently at some step. The world gives one answer to *u* followed by *w*; at that step it differs from the answer after one of the two states at least, which is thereby distinguished from *u*. Each question removes one candidate or more. A program that identifies *u* with no state takes, after *u*, the steps that experience took after *u*, and no others. For a situation *σ* let *q*_*σ*_ be the probability that an experience contains it. The probability that a new experience contains *σ* and *N* earlier ones do not is

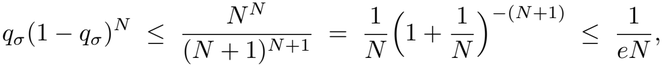

since the left side is largest at *q*_*σ*_ = 1*/*(*N* + 1) and (1 + 1*/N* )^*N*+1^ *≥e*. Summing over the *S* situations gives the bound. For the last claim, by exchangeability the probability that the last of *n* + 1 moments is the only one in its situation is the expected number of situations met exactly once among the *n* + 1, divided by *n* + 1; this is the estimate of Good (1953).

##### How the learner uses part (i): thresholds are solved

The learner finds a threshold by elimination. It takes the moments at which the answer changes from one step to the next, grouped by the new answer, and the moments at which minimal pairs show a dependence that no recall accounts for. For each group it eliminates on the counts of events at those moments: symbols heard of each kind, answers given, records of each content returned by each recall (the counter *D*^*ρ*^ is turned whenever recall *ρ* returns *σ*) and, for a recall with a filter, the records of each content that the filter admits. Where no form vanishes at all of the moments, it takes the forms on which the largest shares of them agree (Theorem 2(ii)). A form is kept if it is within the vocabulary of the recalls, gains up to three and start up to two, and if its side removes more contradictions than it costs to state (Appendix C.2). In binary addition the counters “ones returned by the first recall” and “ones returned by the second” are as good as “ones heard”, and the form the learner solves for is 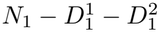: the ones heard less the ones returned so far. It is zero when nothing is left to add, which is where the sum ends (Figure 2b). No minimal pair shows this threshold as a relation between two records: every digit of both numbers moves it, a dependence by count in the sense of Lemma 1. The learner keeps at most one threshold of a counter, since two at different starts would read the counter in pieces, which the memory of Eq. (1) does not allow (Appendix C.2 shows what parity becomes without this rule).

###### Algorithm B.1

Consolidation of a program in step (Law.sleep in dm/law.py)

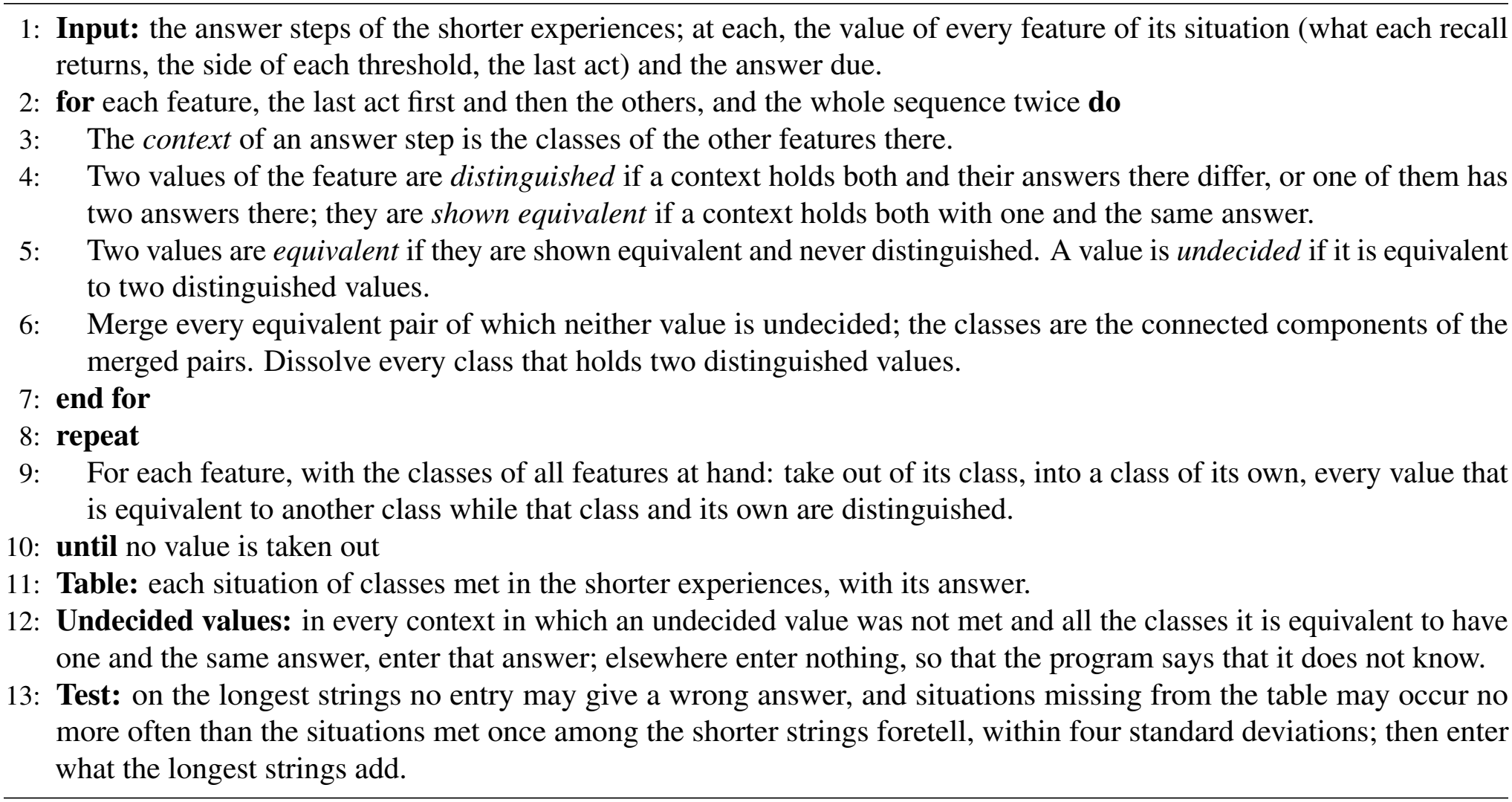

##### How the learner uses part (ii)

For a program in step the values are those of the recalls, the thresholds and the last act, and the classes of the last act are the states. In binary addition the last acts of the standard experience take 23 values, which consolidation sorts into three states, leaving 3 undecided; the states hold the carry from one column to the next (Figure 2c). For inner speech the values are the beginnings of experiences. Appendix B.7 gives consolidation for both as an algorithm.

#### B.7. Consolidation as an algorithm

Consolidation is the rule of Theorem 3(ii) applied to the evidence of an experience. Algorithm B.1 gives it for a program in step as dm/law.py computes it, in the method Law.sleep (the code calls consolidation sleep). It merges only what experience has shown equivalent and never distinguished, leaves undecided what is equivalent to two distinguished things, and compares again what it merged early until nothing changes; from complete experience its classes are those of Myhill and Nerode.

##### Why it compares again

A value is merged while the other features are still unmerged, and it can then be compared with few. When the classes of every feature stand, more contexts coincide, and a value may turn out to be as equivalent to another class as to its own, while the two classes are distinguished. To let the early merge stand would be to let the order of merging decide, which is a guess like any other. For the same reason the features are gone through twice before values are compared again one by one. In binary addition comparing again takes out two last acts that had been merged with the state of no carry and are equivalent to the beginning of a sum as well. The learner that compares only once lets such merges stand (Experiment 2, Section 4.2).

##### Undecided values, and situations never met

An undecided value keeps the entries of the table that it has met, and receives in addition an entry in every context in which all the classes it is equivalent to have one and the same answer. Such entries are inferred, are not counted in the length of the program, and are right if the value belongs to one of those classes (Theorem 3(iii)). Where the classes differ, or one of them has no answer, the table has no entry, and the program says that it does not know. A situation never met is treated in the same way. Theorem 3(iii) bounds how often a new experience meets one: for a program of *S* situations, at most *S/*(*eN* ) after *N* experiences. The learner uses the same fact, counted moment by moment, when it tests a program on the longest strings: the program may say that it does not know on them as often as the situations met once among the shorter strings foretell (Good, 1953), within four standard deviations, and it may give no wrong answer (Appendix B.8).

##### Consolidation of moments: inner speech

For inner speech the values are the beginnings of experiences, and a merge has consequences, since a program is deterministic: if two moments are one, so are the moments that follow them after the same symbol. A merge is therefore tried, its consequences are followed, and it is taken back if two different answers meet (grow in dm/inner.py; Appendix C.3). A merge is supported by each pair of equal answers that it brings together, one from each side. The states are grown from the first moment. A moment that follows a state is compared with every state found so far: if exactly one state supports it, it is that state; if two states support it and do not contradict each other, they are one; if none supports it, it is a new state. Where several states support it and contradict each other, the moment is undecided: it is identified with no state, what follows it is followed like what follows a state, and it is compared again after every change. If the learner can act, it asks (Theorem 3(iii)). The question is a minimal pair made on purpose, by substitution of a beginning: the undecided moment is put in the place of one of two candidate states, in an experience that would have ended otherwise for the other, and the world’s answer is a first difference between the moment and one of its candidates.

##### Comparing the finished table again

Comparing again matters most for inner speech, because its table is grown state by state. A moment at the edge is compared with the states found until then, and if one of them alone supports it the two are identified; a state found later may support the moment as well and be distinguished from the first. So the finished table is compared with itself, every step with every state (undecided in dm/inner.py). A step whose moments the whole table shows to be equivalent to two distinguished states is undecided. If the two states go on alike after every symbol and differ only in what they answer, the moments keep their steps and lose only their answer; otherwise the table is grown again with the moments left undecided, and what is then undecided is asked about or left undecided as before. The false program of the learner that compares only once, announced in 1 of its 300 runs, is a decision of this kind: a counter for the majority of a string that stops counting, decided when its table was half grown (Appendix D.3).

##### Why consolidation seeks no fewest states

A learner that wants the fewest states has to colour the graph of distinguished pairs with the fewest colours, as learners of automata that merge states do (Oncina & Garćıa, 1992; Lang et al., 1998; Heule & Verwer, 2010). The problem is hard in general (Gold, 1978; Angluin, 1978; Pitt & Warmuth, 1993), and where experience is incomplete it has many solutions: a value equivalent to two distinguished values can take the colour of either. The network of the second paper would solve such a problem by its own flow, a multiplier growing on every prohibited pattern that is active until none is (He & Wu, 2026b). The state in which it comes to rest is one of the colourings, and the order of its updates decides which; experience leaves the choice open, and the choice is a guess. The false programs of the learner made to guess (Experiment 2, Appendix D.3) are guesses of this kind and of no other: a last act that could be the carry or the end of a sum, and a moment of an arithmetic expression that could be any of five states of equal value. Consolidation gives up the fewest states for the states that experience has decided. It may keep apart two values that a complete experience would merge, at the price of strings on which the program says that it does not know, and it merges nothing that experience has left undecided.

##### A circuit for consolidation

The rule is local. Whether two values are merged depends on the evidence about the two, that they were shown equivalent and never distinguished, and on the evidence one step further, whether either is equivalent to two values that were distinguished. A circuit computes it in two layers (code/neural/sleep.py). Give each value, of a recall or of the last act, a unit; let two equivalent values excite each other; and let every distinguished pair, a prohibited pattern of the second paper (He & Wu, 2026b), drive an interneuron that silences every unit that both of its members excite: such a unit is undecided. Activity that starts in one unit spreads through the units that are not silenced, and the units it reaches are the class of that value; a class in which a prohibited pair is active is no class, and its units fall back to themselves. The veto then acts once more, on the assemblies. A value was merged while the other recalls were unmerged; when their classes stand, more contexts coincide, and the value is excited by more. A unit that an assembly other than its own excites, and that is in no prohibited pair with it, while a prohibited pair lies between the two assemblies, was undecided: it falls back to itself, and so on until nothing changes. One motif does both: a prohibited pair silences what is equivalent to both of its members, and then what is equivalent to both of their assemblies. On the evidence of every recall of the programs in step learned for the benchmark and the withheld tasks (21 programs), this circuit gives the classes of the learner in 46 of 46 cases, after at most 4 sweeps of spreading (Appendix D.8).

#### B.8. Nothing is guessed

**Theorem 4 (Nothing is guessed)**.

i. *Fix bounds on the gains and starts of counters and on the numbers of recalls, of thresholds and of states. The programs of Eq*. (1) *within the bounds are finitely many, and for every task there is a size N*_teach_ *such that each of them that answers the task on all inputs up to N*_teach_ *answers it on every input, or, with periodic counters as in the brain, on every input within their range*.
ii. *Let a program be chosen as a function of experiences of length less than L, and let it be announced only if it answers each of m experiences of length L drawn independently of the choice. If the program errs on a fraction ε of the strings of length L, it is announced with probability at most* (1*− ε*)^*m*^ *≤e*^*−εm*^; *if k programs chosen in this way are tested on the same experiences, one that errs on a fraction ε is announced with probability at most k*(1*− ε*)^*m*^, *the sum of their bounds*.
iii. *On every string an announced program gives the answer due, or meets a situation that is not in its table and says that it does not know, or gives a wrong answer. It gives a wrong answer only in a situation in which it has taken for one two moments with different answers due. A program whose recalls, thresholds and table agree with a program of the task on all situations of its table gives no wrong answer*.

##### Intuition

A program that uses coincidences alone is the same finite object at every length, because its table never sees a reading; short inputs therefore teach it completely. The learner cannot know how short is short enough, and the test on the longest strings bounds that risk. A wrong answer needs two moments taken for one, and the learner takes for one only what experience has shown equivalent and never distinguished.

*Proof*. (i) A program within the bounds is a choice among finitely many relations and filters for each recall and a table over finitely many situations. Each program of the class that fails on some input fails on one of smallest size, and *N*_teach_ is the largest of these finitely many sizes. This is Theorem 1(ii) of He & Wu (2026a): the situation of a program holds records, coincidences and a state, never a reading, so the class is finite and the same at every size.

i. The program is fixed before the *m* strings are drawn, and each is answered correctly with probability 1 *−ε*, independently of the others. For *k* programs tested on the same strings, the probability that one that errs on a fraction *ε* passes is at most the sum of the *k* probabilities, by the union bound.
ii. The three cases are exhaustive. Every entry of the table was learned from moments of experience with their answers, or, for an undecided value, from the common answer of the classes it is equivalent to. A wrong answer at a moment whose situation is in the table therefore means that the entry came from moments with another answer due, which the program has placed in the same situation. If the recalls and thresholds are those of a program of the task and the table agrees with it wherever it is defined, then on a string all of whose situations are in the table the two programs act alike.

##### Three sizes, and the test

*N*_teach_ bounds the worst program of the class. For the program the learner writes three smaller sizes matter, one for each thing learned: the size *n*_0_ at which the dependences span their space (Theorem 2(iii)), the size *N*^*\**^ at which the program has met its situations (He & Wu, 2026a), and the size at which the experience contains the minimal pairs that show its dependences (Theorem 1(iii)). The learner does not know these sizes, and part (ii) takes their place. Minimal pairs, dependences, counters, recalls, thresholds, the table and its merges are all found from the experiences shorter than the longest, and the longest are only asked. The program has to be right wherever it speaks on them; it may say that it does not know on them as often as the situations it has met once foretell (Theorem 3(iii)), within four standard deviations; and what the longest strings add is then entered in the table. With *m* = 64 strings, a program that errs on a tenth of the strings of the longest length passes with probability below 0.002.

##### What the test reaches

The test bounds the errors of a program on strings as long as the longest it has heard. Against errors that only a longer string would show, a state reached only after many symbols or a sum that ends in a way no short sum does, the learner relies on part (i), since a relation between readings mentions no length, and on the rule that it takes for one only what experience has decided (Theorem 3(ii) and (iii)): an entry of its table that no experience has decided is absent, and the program says that it does not know where it would have been used. The test is on length, and not on a random share of the experience, because a relation fitted to the range of the experience, such as a counter whose gain matches the longest operand heard, fails first at the next length. Setting the longest strings aside costs few minimal pairs: among 64 random binary strings of twelve symbols Theorem 1(iii) expects about six, while the short strings, which the experience samples densely, hold most of its pairs. Without the test the learner announces false programs where experience is thin: in Experiment 2 the learner that announces untested programs, found from all its strings, announced 14 false programs in 1,500 runs, and the complete learner 0 on the same experiences (Section 4.2, Appendix D.3).

#### B.9. Experience made by the machine

Let *P* be a program the learner has learned. To *compose* with *P* is to run it on what was heard before answering: the experience of the new task is *x* followed by what *P* says to *x*, each thing said being an event of a kind of its own, and then the answer steps. An answer ends with the end symbol; *nothing* is the answer that consists of the end symbol alone, and it is the answer to a string that is no question.

**Theorem 5 (Experience made by the machine)**. *Let the learner have learned programs P and ⊕*.

i. Composition. *Let x, x*^*′*^ *form a minimal pair at s, and let their answers to a new task differ first at a step t. In every program for the new task that composes with P, the step t depends, in the sense of Lemma 1, on s or on a moment at which what P says to x and to x*^*′*^ *differs, and the comparison of the two experiences finds these dependences however many moments P took. Learning the new task examines the relations between the counters of the answer steps and the counters under which x and what P said were stored, and counts a table over what they return; no relation, filter or entry of P is examined again. There are tasks that no program within given bounds answers while it says only its answers, and that have a program of three situations over what a learned program says; the sum of two numbers said highest digit first is one (Proposition 7*).
ii. Trial. *Let⊕ be a program whose jth answer is fixed once its question is fixed up to a place l*(*j*) *that does not decrease with j, as the jth digit of a sum said lowest digit first is fixed by the first j digits of each number. Given the beginning u of a question and an answer v, a continuation d with ⊕* (*u d*) = *v, if there is one in which each symbol is determined by the answers up to it, is found by trying, for each place of d in turn, each of the a symbols, and keeping the one under which the answer of ⊕ agrees with v wherever it already speaks: at most a* |*v*| *runs of ⊕. If a symbol χ of a question stands for its answer and a program predicts the last symbol of every true statement, the answer is found with one run of that program for each possible answer*.
iii. Recursion. *Let f be a task, with f* (*x*) *nothing where x is no question, and let D*(*x*) *be the answer d such that ⊕, given f* (*x*^*−*^) *followed by d, answers f* (*x*); *⊕ answers d to nothing followed by d, and v to v followed by nothing. Then f* (*x*) = *D*(*x*_1_) *⊕ D*(*x*_1:2_) *⊕ · · · ⊕ D*(*x*), *the sums taken from the left. If D and ⊕ have programs in step, f* (*x*) *is computed with* |*x*| *runs of the program of D and at most* |*x*| *runs of ⊕, a number of steps of Eq*. (1) *of order* |*x*| ^2^ *when answers are no longer than questions; f itself need not have a program in step*.

##### Intuition

What the learner says is stored and counted like what it hears, so a program already learned can produce experience. Its answers become events on which later answers depend; its runs on candidate answers become imagined experiences that differ in one place; and its answers to the successive beginnings of a question become a table of differences, to be added up as Babbage’s engine was designed to add them.

*Proof*. (i) Lemma 1 applies to the experience made of what was heard and what *P* said, with the set of moments at which the two experiences differ in place of the single moment *s*: every chain of dependences ends at one of them. What *P* says is fixed once *x* is; it is stored and turns counters like any event, and the learner of Appendix C.2 is applied to the longer experience, with minimal pairs taken among the strings heard. The comparison sees the two ends of each chain, so the moments that *P* took do not enter. Counters that time the world, those reset by a symbol, stand still while the learner speaks to itself. The last claim is Proposition 7.

(ii) By assumption the answers of *⊕* up to the *j*th are fixed once the question is fixed up to place *l*(*j*). Suppose the first *i* symbols of *d* are right. Under the right symbol at place *i* + 1, the answers that the question so far fixes agree with *v*; under a wrong one, by the assumption on *d*, some answer that is already fixed differs from *v*. An answer may end early while the question is incomplete; the learner then waits, taking a symbol under which the answer does not yet reach the place in question, and checks the complete *d* against all of *v*. Each place costs at most *a* runs. For a statement, the possible answers are tried in the place of *χ*, and those under which the prediction of the last symbol is wrong are refuted.

(iii) By the definition of *D, f* (*x*) = *f* (*x*^*−*^) *⊕ D*(*x*) for every *x*, also where *x*^*−*^ or *x* is no question. Unfolding along the beginnings of *x* down to the empty one, whose answer is nothing, gives the sum. Each term needs one run of the program of *D* on a beginning of *x* and one run of *⊕* on two answers, and a run of a program in step on a string of *n* symbols with *m* answers takes *n* + *m* steps. □

###### Proposition 7

(A sum said highest digit first). *Let the task be to say the sum of two numbers of L digits, each heard lowest digit first, in L* + 1 *digits, highest digit first. For every bound of Theorem 4(i) there is an L such that no program within the bound that says only its answers and recalls at answer steps gives the first answer on all inputs. Composed with a program for addition, the task has a program with one recall and a table of three situations*.

*Proof*. After addition, answer *j* is the *j*th thing said counted from the last: a counter turned forward by each thing said and back by each answer, recalled against itself at offset +1 (Theorem 2(iv)), and the table says the digit returned, or ends when nothing is returned.

For the first claim, fix a program within the bound. At the first answer step it has said nothing, and its situation consists of the records that its recalls return at that step and of coincidences between readings that depend on the numbers of symbols of each kind in the input, the *totals*. Each recall returns the latest record of some kinds at which a counter stood at a reading fixed by the totals. Take two inputs that agree except in two neighbouring columns, which hold (0, 0) below (1, 1) in the first input and (1, 1) below (0, 0) in the second, and in which every higher column holds a one and a zero. The totals are the same. The top digit of the sum is the carry out of the upper of the two columns, which every higher column passes on: one in the first input, zero in the second. The two inputs have the same records, under the same readings, outside a window of at most four records around the two columns in each number (counters reset by a digit agree again once each digit has occurred after the window). A recall therefore returns the same record in both inputs unless, in one of them, it returns a record of the window, and this requires that the reading of its key counter at that record equal its target and that no later record of its kinds have that reading. For a relation whose two sides do not depend on the counts, the latest record with the reading lies above the window if the higher columns contain every arrangement of four records that the window can contain. For every other relation the requirement is a linear equation between the counts of the symbols below the window and the totals. Finitely many relations give finitely many such equations, and for *L* large there are inputs of the form described that satisfy none of them. On such a pair of inputs the program is in the same situation at the first answer, and the answers due differ.

##### What composition is for

The restriction to programs that say only their answers is the point. A program whose acts are its answers has no state but its counters and its last act (Theorem 3(ii)); any other state would have to be an act that the task does not ask for, something said to oneself. Lemma 1 explains why minimal pairs do not teach such acts directly: a dependence is found by comparing what is heard with what is answered, and a chain that passes through moments at which nothing is answered leaves no trace there. Once the acts said to oneself are the answers of a program that was learned as a task of its own, the chain passes through events again, and each link is a dependence like any other. This is the order in which procedures are taught, the partial result first written down and later only thought (Vygotsky, 1978). The first paper showed that with steps to think the memory computes whatever a computer computes (He & Wu, 2026a); the theorem says of what the steps are made.

##### Trial is difference in imagination

The trials of part (ii) differ in one place, the place being tried: they are minimal pairs that the learner makes for itself, and the answer that tells them apart is its own prediction. For equations the learner first has to find which symbol stands for the answer. It tries each symbol that occurs exactly once in every question except the shortest; for each, it puts the answers in that place, which turns its experience into statements, and looks for a program that predicts the last symbol of a statement from the rest (learn trial in dm/closure.py). For one symbol such a program exists, 15 states of inner speech with a counter of depth for the brackets (Appendix C.3); for the others the statements contradict one another. Nothing told the learner what an equation is.

##### Recursion is the method of differences

Babbage’s difference engine was designed to build a table of a function from its own last entry by adding differences that follow a simpler rule than the function (Babbage, 1864). Part (iii) is that method with two changes: the differences are taken along the beginnings of a question, where Babbage took them along the integers, and both the taking of differences and their adding are done with programs the learner has learned (learn recursion and invert in dm/closure.py). For multiplication, heard as two numbers lowest digit first, the neighbour by deletion of a question lacks the highest digit of the multiplier, so the difference of the two products is the multiplicand shifted by the place of that digit if the digit is one, and nothing if it is zero. This has a program in step with 11 situations: a recall that walks the multiplicand at a lag equal to the number of digits of the multiplier so far, a recall that returns the last digit heard, and thresholds, one of which ends the answer when the ones of the multiplicand have all been returned (Appendix C.6; Figure 2d). The learner does not know that it is multiplying; it finds that the differences of its answers have a program, and that suffices. Where a beginning is no question, a number without a multiplier, its answer is nothing, and the program of the differences learns to say nothing there, so that the sum along the beginnings needs no rule for where to start. To take the differences the learner needs the answers to the beginnings of its questions. Those it does not remember it asks for, with a question of a directed kind: the same question with one symbol less.

#### B.10. A local rule

Elimination uses all dependences at once; a circuit meets them one at a time. Let gains and offsets be real numbers, write a relation as *r* = (***θ, θ***^*′*^, *δ*) and the point of the *i*th dependence *s*_*i*_ ↣ *t*_*i*_ as *p*_*i*_ = *p*(*s*_*i*_, *t*_*i*_) = (*−***N**(*s*_*i*_), **N**(*t*_*i*_), 1), so that the miss of Eq. (3) is *µ*_*i*_ = *r·p*_*i*_. For a counter recalled against itself, ***θ*** = ***θ***^*′*^ and *µ*_*i*_ = ***θ*** *·*(**N**(*t*_*i*_) *−* **N**(*s*_*i*_)) + *δ*.

**Theorem 6 (A local rule)**.

i. *The constraint µ*_*i*_ = 0 *acts on the relation with the force −µ*_*i*_*p*_*i*_, *the negative gradient of* 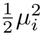. *Its component on the gain of a kind of event is the miss times a number of events of that kind, with the sign that reduces the miss: for the query counter the number up to the answer, for the key counter the number up to the record, with the opposite sign, and for a counter recalled against itself the number between the record and the answer*.
ii. *Let* 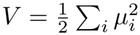. *The flow* 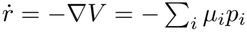 *converges to the relation that holds at all the dependences and is nearest to r*(0): *the projection of r*(0) *on the space of Theorem 2(i). So does the rule that, at each dependence met in experience, changes r by η −µ*_*i*_*p*_*i*_*/*| *p*_*i*_| ^2^ *with* 0 *< η <* 2, *when the dependences recur in turn or in random order; in random order each dependence multiplies the expected squared distance to the limit by at most* 1 *−η*(2 *−η*)*κ, where κ is the smallest positive eigenvalue of the mean of* 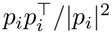. *The same holds with some components of r held fixed, whenever a relation with those components holds at the dependences; if the dependences then have one solution, it is the one that elimination finds. A counter that is run in a ring with wells moves by the whole number nearest to each gain: if the solution is whole, the counter is exact once its gains are within one half of their whole values, and it stays exact from the moment their distance to the solution is less than one half, while the flow or the rule carries them on to the solution; since they converge, this happens after finitely many dependences*.
iii. *Backpropagation through time on the counters, run one event at a time, computes the force of part (i). The correction for a dependence is a function of the miss and of the readings of the counters at the two moments; it is computed when the miss occurs, and no error is carried back through the moments between them*.

##### Intuition

Each dependence pulls the gains toward the plane on which its miss is zero, and that plane passes through every relation that holds at all the dependences; pulling toward one plane after another converges, as in the method of Kaczmarz, to the solution nearest the start. The pull on each gain is the miss times a count of events, which the counters already hold, so the correction is computed where the miss occurs. The wells of a ring make a gain that is nearly whole act as whole.

*Proof*. (i) *∂µ*_*i*_*/∂r* = *p*_*i*_, whose components are *−***N**(*s*_*i*_) on the key gains, **N**(*t*_*i*_) on the query gains and **N**(*t*_*i*_) *−***N**(*s*_*i*_) on the gains of a counter recalled against itself.

(ii) *V* is a convex quadratic form that vanishes exactly on the relations that hold at all the dependences: a linear space *U* by Theorem 2(i), or an affine one when components are held fixed. Each *p*_*i*_ is orthogonal to *U*, so the flow moves *r* within *r*(0) + *U*^*⊥*^, and on *U*^*⊥*^ the form is positive definite; *r* therefore converges, exponentially, to the one point of *r*(0) + *U*^*⊥*^ that lies in *U*, which is the projection of *r*(0).

For the rule, let *r*^*\**^ be that projection and *z* = *r − r*^*\**^, which lies in *U*^*⊥*^ at the start and stays there, since each step adds a multiple of some *p*_*i*_. As *µ*_*i*_ = *z·p*_*i*_, a step at dependence *i* turns *z* into *z*^*′*^ with

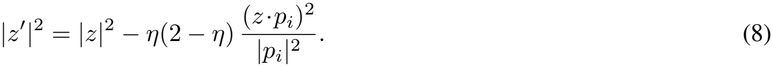

For 0 *< η <* 2 the distance to every point of *U* thus never increases, and for *η* = 1 the step is the projection of *r* on the plane *µ*_*i*_ = 0, which contains *U* : this is the method of Kaczmarz (1937). When the dependences recur in turn, a full round over them shortens every *z ≠*0 in *U*^*⊥*^, since a round that kept |*z*| would change nothing and leave *z* orthogonal to every *p*_*i*_, hence in *U* ; by compactness of the unit sphere of *U*^*⊥*^ a round shortens *z* by a fixed factor, and *z* tends to zero. When the dependence is drawn at random, the expectation of the last term of Eq. (8) is *η*(2 *η*) *z*^*⊤*^*Mz* with *M* the mean of 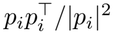, and *z*^*⊤*^*Mz κ≥* |*z*| ^2^ on *U*^*⊥*^, which gives the factor stated (Strohmer & Vershynin, 2009). The expected value of |*z*| ^2^ thus tends to zero, and since *z* never increases it tends to zero with probability one: *r* converges to the projection of its start. With one solution, the flow, the rule and elimination solve the same equations and end at the same point. A gain within one half of a whole number acts in a ring with wells as that number, so a counter whose gains are within one half of a whole solution moves as the solution does. The Euclidean distance of the gains to the solution does not increase, by Eq. (8) for the rule and because 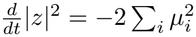 for the flow; once it is below one half, every gain is within one half of its whole value and stays so.

(iii) The components of *p*_*i*_ are the numbers of events of each kind up to the record and up to the answer. They are the readings, at the two moments, of the counters that count one kind each: the reading at the record was stored with the record, and the reading at the answer is present. A backward pass through *c*(*u*) = *c*(*u −* 1) + *θ*_*c*_(*e*_*u*_) carries the derivative of 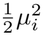 with respect to *c*(*u*) back unchanged, since *∂c*(*u*)*/∂c*(*u−* 1) = 1, and at each moment adds it to the derivative for the gain of the kind of event met there; it therefore returns *µ*_*i*_*p*_*i*_, the force with its sign reversed (Appendix D.9 checks this to 2 *×* 10^*−*13^).

##### A descent computed where the miss occurs

The rule is a descent on the squared miss: its correction is the derivative of 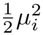 with respect to the gains, divided by |*p*_*i*_|^2^. The error it descends is a displacement in memory, the distance between where the query counter stands at the answer and where the record was stored, and it is computed where the miss occurs, from the miss and two readings. No error is carried back through the steps of the computation. Backpropagation through time would obtain the same derivative, for a parameter that acts at every moment between *s* and *t*, by carrying an error back through those moments (Werbos, 1990). For a counter the sum over the moments has already been formed, by the counter itself, because the turn that an event gives depends only on its kind; if the turn depended on the reading or on a state of the controller, the counts would no longer be the derivative. Appendix D.9 checks the equality numerically and shows how it fails in that case. The miss is an instructive signal delivered to the synapses that set the gains, of a kind that cortical neurons are known to receive (Francioni et al., 2026).

##### The offset and the scale

A relation and every multiple of it hold at the same pairs, so the dependences fix a direction and not a scale. The offset of a recall is a property of the recall, to be held fixed during the flow. With the offset of a reversal held at +1 the dependences have the one solution (+1, *−*1). With the offset left free the solutions are the line through zero spanned by (1, *−*1; 1), and the flow ends at its point nearest the start, 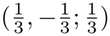 from gains (1, 1) and offset 1, a counter that every event turns forward. Between two counters the scale is set by the key counter, whose gains the memory of records already has; if both sides are left free and some dependences are wrong, the gains decay to zero, the one relation that holds everywhere (Appendix D.9).

##### Whole numbers, and where the graded miss comes from

A counter is a ring of attractor states, its wells (He & Wu, 2026a). When the counter is run, events are slower than the wells: a pulse of gain *g* carries the bump *g* wells, the wells catch it, and the counter moves by the whole number nearest to *g*. A real gain within one half of its whole value thus acts as that value, and every dependence of every length is served; without wells a gain in error by *ϵ* serves dependences only across spans of fewer than 1*/*(2*ϵ*) events. The number of dependences served therefore stays near zero while the gains travel and reaches all of them when the gains arrive: the flow is gradual and what it achieves is sudden (Figure 4d; Appendix D.9). The flow itself needs a graded miss, which a counter that rounds every event does not give, and a sweep gives it. When the stretch between the recalled record and the act is replayed, its events follow one another faster than the wells act, the bump adds their pulses as they are, and it arrives beside the record at a distance of ***θ****·* Δ**N** less the whole number due: the miss *µ*_*i*_ of the linear equation. In simulation the travel of the bump is the sum of the pulses to within a fraction of a well (Appendix D.8). The same ring is exact when it is run and linear when it replays, and the difference is one of pace.

##### Dependences that a counter cannot serve

The dependences of an experience do not all belong to one recall, and the convergence above assumes that they do. Let the rule be fed with all the dependences of an experience, and let the miss be felt through the field of the place cell of the record, as the response of a cell of width *w* to a displacement *µ* is: the step is *−η ψ*(*µ* ) *p /*|*p* |^2^ with 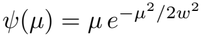. Let *r*^*\**^ be a whole relation that holds at a share 1 *− ϕ* of the dependences. At *r*^*\**^ the others miss by whole numbers, since gains and counts are whole, and so by at least one well. Near *r*^*\**^ the dependences at which it holds pull as before, with slope *ψ*^*′*^(0) = 1, and each of the others pulls with a force of size at most 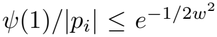 (for *w ≤* 1, and since |*p*_*i*_| *≥* 1), which is *e*^*−*2^ *≈* 0.14 for a field half a well wide. At a rest point *r*^*\**^ + *z* of the mean step, to first order in 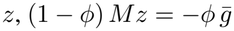, where *M* is the mean of 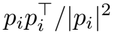 over the dependences at which *r*^*\**^ holds and 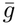, the mean of *ψ*(*µ*_*i*_) *p*_*i*_*/* |*p*_*i*_| ^2^ over the others, has size at most 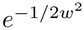. The rest point is therefore displaced from *r*^*\**^ by at most

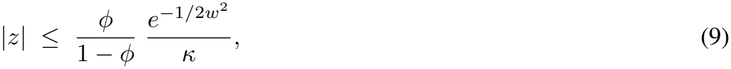

where *κ* is the smallest eigenvalue of *M* . Pulls of opposite sign cancel, and the displacement is usually far smaller. When it is below one half the wells round it away. With a miss felt in proportion at any distance, the factor 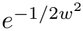 is replaced by the size of the misses, one or more, and the gains settle between the solutions of different recalls. The whole numbers thus serve twice: they make a learned gain exact, and they keep the dependences of other recalls a whole well away, where a field does not feel them. The algebra reaches the same end by taking the relation on which most pairs of dependences agree (Theorem 2(ii)). In the standard experience of the stack 26% of the pairs of dependences cannot be served by the counter of the stack; a ring fed with all pairs, unsorted, learned that counter in 20/20 trials with the field and in 0/20 without it (Figure 4d; Appendix D.8).

##### The multiplier

In the second paper a constraint acts through a multiplier *λ*_*i*_ that integrates its violation and leaks, 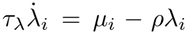,while the state moves under the force *−λ*_*i*_ *∂µ*_*i*_*/∂r*, so that 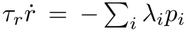 Eliminating the multipliers gives

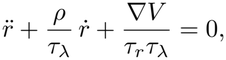

descent on *V* with the multipliers as momentum and their leak as friction. Each mode of the dependences, with singular value *σ*, obeys the rhythm law of the second paper, 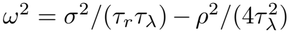. Without a leak the gains circle the solution indefinitely. With a leak they converge to the same limit as in part (ii), since 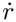 stays in *U*^*⊥*^; and when the leak is fast the multiplier is the miss itself, *λ*_*i*_ = *µ*_*i*_*/ρ*, which is the flow of part (ii). This is the sense in which the multiplier of a dependence is its miss.

#### B.11. What experience can write

##### The class reached

Programs in step compute, in the terms of the first paper, what a multitape machine computes in real time (He & Wu, 2026a). Composition composes such programs, trial inverts one at the cost of a factor *a n*, and recursion iterates one along the beginnings of the question, *n* times. Each multiplies the running time by a polynomial, so everything learned by a bounded number of these operations runs in polynomial time. A program in step takes a time proportional to the length of what it hears and says; a procedure learned by recursion takes a time that grows as its square, as the learned multiplication does, whose moments grow as the power 2.02 of the length (Appendix D.7). On-line, where each digit of a product must be said before the next digits of the factors are heard, multiplication has no program in step (Cook & Aanderaa, 1969); whether it has one when the factors are heard first we leave open. The learner finds none, and learns the product by recursion.

##### The frontier

The memory runs everything a Turing machine computes (He & Wu, 2026a), and experience writes less. The learner reaches a procedure through programs it has learned, so a procedure whose parts were never tasks of their own lies beyond it, as the square root does, which asks for the inverse of squaring. The stored program lies beyond it as well: Turing’s universal machine, which keeps its own table in memory and recalls it as it recalls the world. The theory takes memory, tasks and kinds of event as exact; with noise a single difference becomes a statistic over many pairs, which sets the pace at which an animal learns (Sun et al., 2025). The proposition below states what the two ingredients of the learner contribute, and so places it between two limits.

###### Proposition 8

(Episodes and counters). *Let a learner write programs of Eq*. (1) *from experience*.

i. *Kept experiences give credit to causes that act jointly: one comparison of a minimal pair finds a dependence whatever the other causes of the answer. A learner that keeps no experience credits a recall by statistics of single experiences, which credit only recalls that act alone; for causes that act jointly it has to credit recalls in sets, or move all parameters at once along a gradient of the error*.
ii. *Counters give generalization across lengths: a relation between their readings is fixed by the dependences of short strings and holds at every length. A learner that keeps experiences and has no counters finds the dependences of each string it has met as pairs of positions, and carries them to another length through a rule that generates them, which it has to find by search among the expressions of a language of positions*.

##### Intuition

Credit needs a second experience to compare with, and generalization needs a quantity that does not mention length. Without the first a learner is left with gradients, and without the second with search.

*Proof*. (i) The first sentence is Theorem 1(ii). A learner that keeps no experience can compute only statistics of experiences taken one at a time, and by Theorem 1(i) these do not credit a recall whose effect is joint with another’s; recalls then have to be credited in sets, or all parameters moved at once along a gradient of the error. (ii) The first sentence is Theorem 2(iii). A learner with experiences and no counters finds the dependences of every string it has met (Theorem 1(ii)), as pairs of positions. The pairs met at sizes up to *n* say nothing about size *n* + 1 unless they are instances of a relation that mentions no size, and without readings to relate, such a relation has to be found among the expressions of some language of positions.

A learner without episodes is connectionist learning, which assigns credit by gradients; a learner without counters is program synthesis, which generalizes by search (Solar-Lezama, 2008; Ellis et al., 2021). The learner of this paper keeps episodes and has counters. It takes its data from the first and its exactness from the second, and in place of the search it solves equations (Section 6).

### C. The learner

This appendix describes the learner of Section 3 step by step, for a reader who wants to see how each part of the postulate is carried out, or to read the programs that the learner writes. Appendix C.1 describes the memory for which it writes programs, Appendix C.2 the algorithm and its steps for a program in step, Appendix C.3 inner speech, Appendix C.4 composition, trial and recursion, Appendix C.5 its four constants, and Appendix C.6 the programs that it writes for the benchmark, with a key to their notation (Table 9). The code is code/dm/, about 2,500 lines of Python with NumPy (Appendix G). One function, dm.learn, is the learner, and every experiment on the learner calls it with the same constants. Appendix A works several of its steps by hand.

**Table 9.**
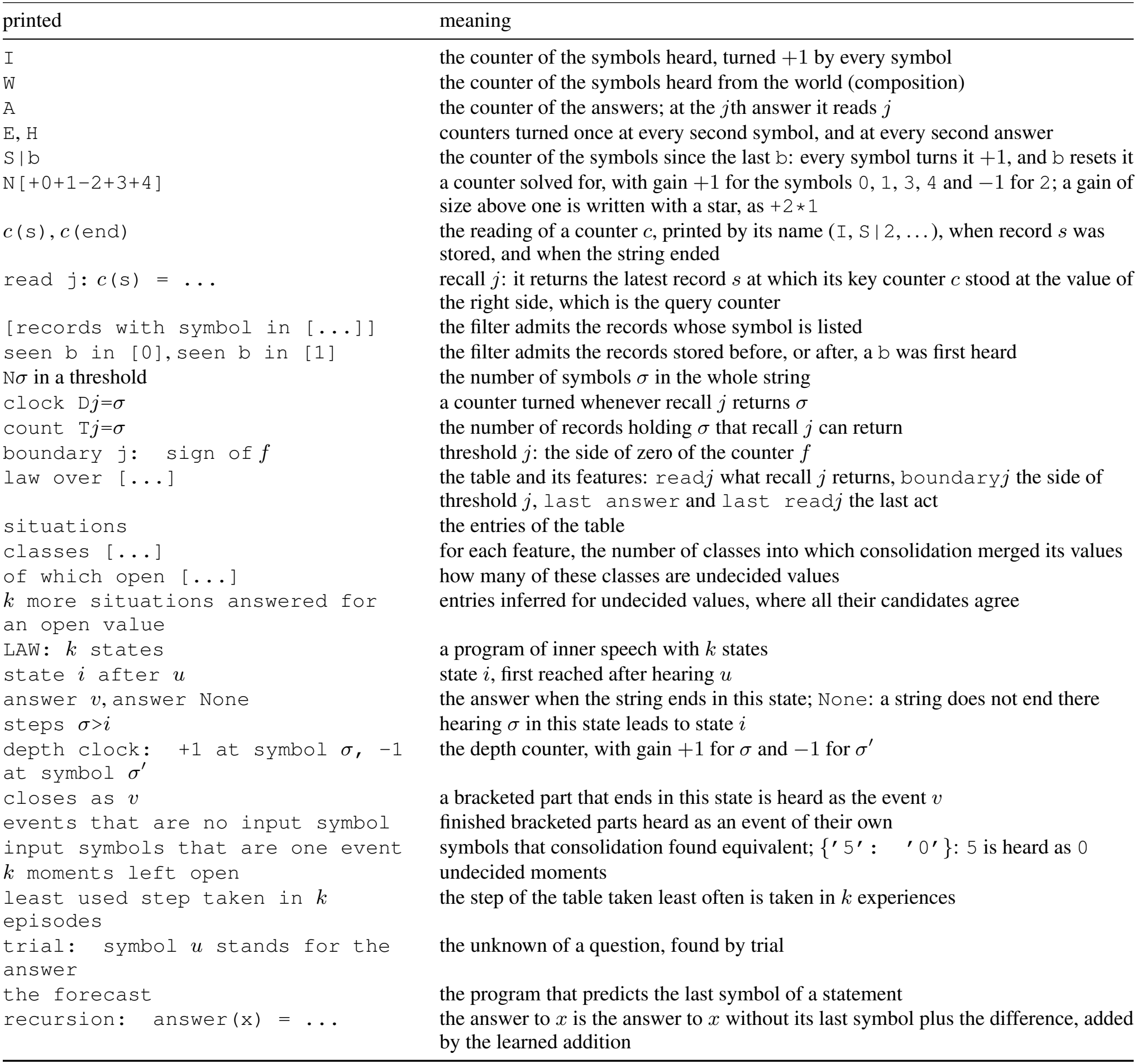
Key to the notation of the listing.

| printed | meaning |
|---|---|
| I | the counter of the symbols heard, turned +1 by every symbol |
| W | the counter of the symbols heard from the world (composition) |
| A | the counter of the answers; at the jth answer it reads j |
| E, H | counters turned once at every second symbol, and at every second answer |
| S\|b | the counter of the symbols since the last b: every symbol turns it +1, and b resets it |
| N[+0+1-2+3+4] | a counter solved for, with gain +1 for the symbols 0, 1, 3, 4 and −1 for 2; a gain of size above one is written with a star, as +2*1 |
| c(s), c(end) | the reading of a counter c, printed by its name (I, S\|2, ...), when record s was stored, and when the string ended |
| read j: c(s) = ... | recall j: it returns the latest record s at which its key counter c stood at the value of the right side, which is the query counter |
| [records with symbol in [...]] | the filter admits the records whose symbol is listed |
| seen b in [0], seen b in [1] | the filter admits the records stored before, or after, a b was first heard |
| Nσ in a threshold | the number of symbols σ in the whole string |
| clock Dj=σ | a counter turned whenever recall j returns σ |
| count Tj=σ | the number of records holding σ that recall j can return |
| boundary j: sign of f | threshold j: the side of zero of the counter f |
| law over [...] | the table and its features: readj what recall j returns, boundaryj the side of threshold j, last answer and last readj the last act |
| situations | the entries of the table |
| classes [...] | for each feature, the number of classes into which consolidation merged its values |
| of which open [...] | how many of these classes are undecided values |
| k more situations answered for an open value | entries inferred for undecided values, where all their candidates agree |
| LAW: k states | a program of inner speech with k states |
| state i after u | state i, first reached after hearing u |
| answer v, answer None | the answer when the string ends in this state; None: a string does not end there |
| steps σ>i | hearing σ in this state leads to state i |
| depth clock: +1 at symbol σ, -1 at symbol σ′ | the depth counter, with gain +1 for σ and −1 for σ′ |
| closes as v | a bracketed part that ends in this state is heard as the event v |
| events that are no input symbol | finished bracketed parts heard as an event of their own |
| input symbols that are one event | symbols that consolidation found equivalent; {′5′: ′0′}: 5 is heard as 0 |
| k moments left open | undecided moments |
| least used step taken in k episodes | the step of the table taken least often is taken in k experiences |
| trial: symbol u stands for the answer | the unknown of a question, found by trial |
| the forecast | the program that predicts the last symbol of a statement |
| recursion: answer(x) = ... | the answer to x is the answer to x without its last symbol plus the difference, added by the learned addition |

**Table 10.** The experiments. Code: the names of the experiments in the code and in the folder results/, and the commands that run them from the folder code/, with MODE=full. The baselines B1 and B2 are run by python3 -m baselines.gradient baselines and by python3 -m baselines.bptt first paper. Solved: exact for every seed.

| Experiment | Claim tested | Design | Main result | Code | Main text | Here |
|---|---|---|---|---|---|---|
| 1. The benchmark without backpropagation | Comparison alone writes exact programs for a standard benchmark from a few hundred short strings. | 15 tasks, *N* = 4 to 20; two gradient baselines. | 13 of 15 solved from the standard experience, 11 of them exact to 10<sup>6</sup> symbols; gradient descent on the same machine: at most five of 10. | E1<br>make e1;<br>B1, B2 | §4.1, Tab. 1, Fig. 3a | D.2 |
| 2. A learner that knows what it knows | The learner, which does not guess, announces no false program, even from thin experience. | 30 tasks, *N* = 3 to 20; three variants. | No false program in 5,680 runs; the variants: 3, 14 and 1. | E3<br>make e3;<br>E7 | §4.2, Fig. 3b | D.3 |
| 3. What each part is for | Each part of the theory carries tasks of its own. | 30 tasks; ten lesions, one at a time. | 28 of 30 solved by the complete learner; from 8 to 27 with one part of the theory removed. | E2<br>make e2 | §4.3, Fig. 3c | D.4 |
| 4. Minimal pairs and questions | Without minimal pairs no recall is learned, and questions make the pairs that experience lacks. | No minimal pairs; questions; more strings. | 6 of 30 without minimal pairs, 28 with questions; arithmetic modulo five from 5,843 strings (11,135 without questions). | E3, E7<br>make e3,<br>make e7 | §4.4, Fig. 3d,e | D.5 |
| 5. Withheld tasks and the constants | The learner solves tasks withheld from its design, and the tasks it solves stay the same when one of its constants at a time is set below or above its value. | 15 withheld tasks; each constant changed. | 15 of 15 withheld tasks solved; 28 of 30 tasks under each of 8 settings. | E4, E8<br>make e4,<br>make e8 | §4.5, Fig. 3a | D.6 |
| 6. Experience made by the machine | The learner extends its own experience, and builds long multiplication from addition. | Composition, trial, recursion. | all four tasks composed; long multiplication, exact to 3,200 symbols. | E5<br>make e5 | §4.6, Fig. 3f | D.7 |
| 7. The engine in neural circuits | The learner runs, compares and learns in rings of neurons. | Rings of rate neurons, Eq. (4). | 7 of 8 programs in step run as circuits; the stack's counter learned in 20/20 trials. | E9<br>make e9 | §5.5, Fig. 4b,d, Fig. 5a,b | D.8 |
| 7. The local rule alone | The rule of Theorem 6 finds the gains with no backward pass. | Reversal and the stack; noise; a backward pass. | Every run reaches the program; the correction equals that of backpropagation to 2 × 10<sup>−13</sup>. | E6<br>make e6 | §5.5, Fig. 5c,d | D.9 |

#### C.1. The memory for which the learner writes

The learner writes programs for the memory of Eq. (1), and nothing in this memory is chosen for a task.

##### Events and counters

The kinds of event are the symbols that can be heard, an answer said, a record of a given content returned by a given recall, and a counter’s return to zero. The learner starts with the counters that need no learning. The counter I is turned +1 by every symbol heard. For every symbol *b* the counter S|b is turned +1 by every symbol and reset by *b*: it counts the symbols since the last *b*. The counter A is turned +1 by every answer. The counters E and H are turned +1 whenever a ring of two states returns to its first state, the ring being turned by every symbol for E and by every answer for H, so that E advances at every second symbol and H at every second answer. When the learner hears its own answers (Appendix C.4), the counter W counts the symbols heard from the world. Every other counter is solved for: a key counter from pairs of dependences (Theorem 2), a threshold counter from the moments at which the answer changes (Theorem 3). The code holds no list of counters. While the learner speaks to itself the counters S|b stand still, since what it says does not turn the counters that time the world.

##### Records

The record of a moment of hearing holds the symbol and, for every symbol *b*, whether *b* had been heard before. These are the kinds of record, and a filter admits records by kind. The moments of hearing are the records that a recall can return; what the learner hears itself say is heard and stored like a symbol from the world.

##### Recalls

A recall has a key counter *k*, a query counter *q* and a filter, and it returns, at moment *t*, the latest admitted record *s* with *k*(*s*) = *q*(*t*). The learner writes the query counter in terms of what turns it. The query starts at an offset *δ ∈{−*2, …, 2 *}*; a symbol turns it by *α* times the gain of the key counter for that symbol, *α ∈ {*0, *±* 1*}*, plus a constant; every second symbol, an answer and every second answer turn it by constants; and at most two of these constants are not zero, each of them *±*1 or *±*2. With *α* = 1 the query is the key counter carried on into the answers, as in a stack; with *α* = 0 it is a fresh counter, as in a queue; with *α* = *−*1 it runs against the key. The code writes a recall whose key counter is *c* as

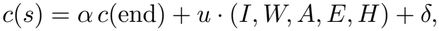

where *c*(end) is the reading of *c* when the string ends, *I, W, A, E, H* the readings of the counters above at the present moment, and *u* the constants; Lemma 2 shows that this is the coincidence of two counters just described. For each key counter there are 1,695 such queries, and 2,715 when the learner hears its own answers and W enters. The learner judges each of them by counting the dependences it meets.

##### Thresholds

A threshold is a counter whose side the table is told: at zero, or the sign with which it left zero. Its gains are over the symbols heard, the answers given, the returns of each recall (the counter *D*^*ρ*^ is turned whenever recall *ρ* returns *σ*) and, for a recall with a filter, the records of each content that the filter admits (the count *T* ^*ρ*^ of such records in the string).

##### The table

The table receives the symbols that the recalls return, or that nothing was returned, the sides of the thresholds, and the last act: the last answer and what the recalls returned at the last step, which carry the state. It never receives a reading. A program in step therefore consists of a few recalls, a few thresholds and a table. Running it takes one pass over the string for each counter used and one look into the table for each answer, and a string of a million symbols is answered in about a second on one processor core.

###### Algorithm 1

The learner (dm.learn). Lines 1 to 11 write a program in step (Appendix C.2); lines 12 to 16 are inner speech, the experience that the learner makes for itself, and its questions (Appendices C.3 and C.4).

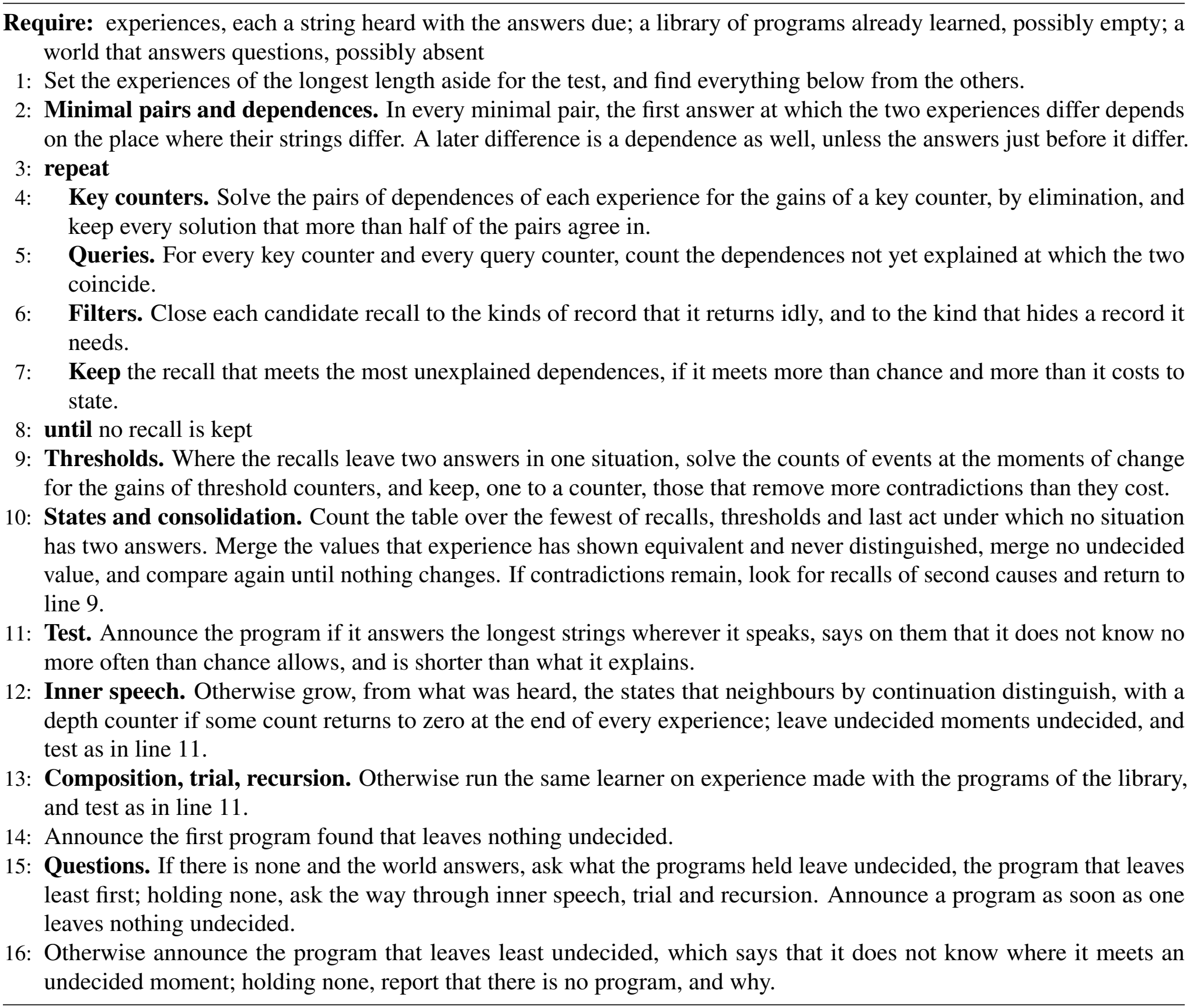

#### C.2. The algorithm

Algorithm 1 follows the order of the postulate of Section 3.2. Lines 2 to 8 are its parts (i) and (ii): minimal pairs give dependences, and the dependences are solved for key counters, queries and filters. Lines 9 and 10 are part (iii), thresholds, states and consolidation, and so is line 12, inner speech with its depth counter. Lines 1, 11 and 14 to 16 are part (iv): everything is found from the shorter strings, a program is announced only if it answers the longest, and what experience leaves undecided is asked about or stays undecided. Line 13 is the experience that the learner makes for itself (Section 3.6). The paragraphs after the algorithm take its steps in turn.

**The longest strings** (line 1). Everything is found from the experiences shorter than the longest, and the longest are kept for the test of line 11 (Theorem 4(ii)). An experience with fewer than three lengths of string leaves nothing to predict, and the learner reports that it has no program.

**Minimal pairs and dependences** (line 2). Minimal pairs are found by hashing every string with one place left open: two strings that agree everywhere but in one place share a key. For a pair that differs at *s*, every answer at which the two experiences differ is recorded, and the first is marked; it is a dependence (Section 3.1). A later difference is a consequence if the answers just before it differ, since the learner’s own last answer may then carry it, as a carry is carried in addition; otherwise it is a dependence as well. While the learner looks for recalls, a difference counts as *explained* if a recall already kept returns different records in the two experiences at that answer or at the one before, or if their last answers differ. The dependences that drive the next recall are the first differences not yet explained, and, once fewer than six of these remain, every difference not yet explained. This is the method of residues inside the method of difference. It lets the learner find the second recall of an addition, whose effect the first recall otherwise masks, and both recalls of the conjunction of two strings, which a test of “always or never” would reject (Appendix B.3).

**Key counters** (line 4). Take two dependences *s*_*i*_ ↣ *t*_*i*_ and *s*_*j*_ ↣ *t*_*j*_ of one experience whose answers are *d* steps apart. A key counter that serves both stands at the two records as far apart as the query moves in *d* answers. With the gain of the query for an answer set to *−*1, which fixes the scale and the sign, this is

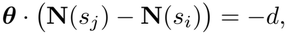

one linear equation in the gains ***θ*** of the key counter, whose coefficients are the numbers of symbols of each kind between the two records (Theorem 2(i); Appendix A.4 works one case). The learner forms these equations first from the answers one or two steps apart and then from any two answers, taking at most 24 dependences of an experience at a time, first for all minimal pairs together and then for each kind of change alone, a kind of change being a given symbol exchanged for a given other; a set of fewer than twelve equations is passed over. It solves the equations together by elimination over the rationals. If no gains solve them all, it solves two hundred random draws of just enough equations to fix a solution, each equation drawn as often as it occurs, and compares the solutions by the number of equations they satisfy; it keeps those that more than half of the equations agree in, the best three at most. There may be more than one. A pair of dependences constrains the gain of a kind of event only if such an event lies between its two records, and in short strings a pop seldom does, so that in some experiences a counter with gain *−*1 for a pop and one with gain *−*2 each satisfy three quarters of the pairs, one pair apart. The pairs do not decide between them, and the learner does not decide either. Both counters go on to the count of recalls, where each dependence counts every event from its record to its answer, and there the counter of the stack meets two thirds of the dependences and becomes the recall. A solution with all gains equal is the counter I, which the learner has from the start, and gains larger than three are refused. The step is repeated after each recall that is kept, on the dependences still unexplained.

**Queries** (line 5). For every key counter (I, the counters S|b and those solved for) and each of its queries, the learner counts the unexplained dependences at which key and query coincide, and the coincidences of the same query with a record drawn at random from the same experience. A candidate goes on only if it meets at least six dependences, and more than the random records do. The candidates are ranked by how many more they meet, the cheaper first among equals, and at most 3,000 dependences are sampled for this count. The 160 best candidates are then looked at in turn, until 40 that return different records have been compared.

**Filters** (line 6). A relation between counters says where to look, and a filter says at what. Two things close a filter. The first is idleness. Among the records that a candidate recall returns and that some minimal pair changes, a kind of record is closed out if it is needed in less than a tenth of its returns, and less often than chance would allow, by four standard deviations, were it needed as often as the other kinds. For a recall of the first number of a sum, these are the records after the separator. The second is hiding. If at some dependence the relation holds but a later record with the same reading is returned instead, the learner tries the filters that tell the two records apart by their kind: the kinds met at the needed records, all kinds but one, and one kind alone. In a stack a pop stands at the reading of the record it uncovers, and the filter is closed to pops (Appendix A.4). A recall is kept only if it returns every needed record at which its relation holds: a dependence is met or it is not (Section 3.3).

**Keeping a recall** (line 7). A recall taken at random returns one record of *n*, so of *k* dependences it would meet *k/n* by chance. A recall is kept only if it meets more than this by four standard deviations, and if it meets more dependences than the bits it takes to state it, counted at log_2_ *n* bits a dependence, with *n* the mean length of the strings: these bits are log_2_ of the number of relations counted so far, plus a margin of twenty bits. Among the recalls that pass, the one that meets the most unexplained dependences is kept. If several meet as many, up to chance, the learner keeps the one that tells most, in bits, about where a change of a record makes a difference at the answer that returns it, less its cost (Appendix B.3, idle returns). The cost of a recall is the sum of the sizes of its gains and constants. The search ends when no candidate passes, and after eight recalls at most.

**Thresholds** (line 9). The recalls that meet at least 15% of all differences are part of every table. If they leave two answers in one situation, the learner looks for thresholds (Theorem 3(i)). It takes the moments at which the answer changes from one answer to the next, grouped by the new answer, and the moments at which a minimal pair shows a dependence that no recall explains, grouped by the answer due. For each group with at least four distinct rows it eliminates on the counts of events at those moments: the symbols heard of each kind, the answers given, the records of each content that each recall has returned (the counters *D*) and, for a recall with a filter, the records of each content that it can return (the counts *T* ). A count that the others already give at every moment is left out. Where no form vanishes at all the moments of a group, the learner takes the forms that large shares of them agree in (Theorem 2(ii)), at most four from a group. A form is offered if it lies within the vocabulary of the recalls, with gains up to three and a starting value up to two, and if its side is not the same at every moment. The forms are then taken one at a time, the one that removes most contradictions first, as long as each removes more contradictions than it costs to state: log_2_ of the number of forms found, plus the margin, counted in units of log_2_ of the number of different answers. At most twelve are offered to the table.

##### One threshold to a counter

The learner keeps at most one threshold of a counter. Two thresholds of one counter at two starting values would tell the table whether the counter stands at one, at three or at five: a reading, read off in pieces, which the memory of Eq. (1) does not allow. A staircase of thresholds fits any finite experience and no longer one. Without this rule the learner fits parity, from strings of up to eight symbols, by three thresholds on the count of 1s, a program that is false from seven 1s on; with it, the learner finds no program in step and learns parity as two states of inner speech (Appendix A.5). In binary addition the threshold found is the number of 1s heard less the number of 1s returned by the two recalls, which is zero when nothing is left to add (Appendix A.6).

**The table and consolidation** (line 10). Call each thing that the table receives a *feature*: what a recall returns, the side of a threshold, the last act. The table is sought over the smallest sets of features first. The recalls that meet at least 15% of the differences are always in it; the last answer, what the recalls returned at the last step, the thresholds and the other recalls are added, up to four of them, and beyond four by adding the feature that removes most contradictions. A task with a single answer at the end has no last act. A set is accepted if, counted on the shorter strings, no situation has two answers, and if the program passes the test of the longest strings. Consolidation is the rule of Theorem 3(ii), applied to each feature in turn, the last act first. A *context* of a value is the rest of the situation, the values of the other features. Two values met in one context with one answer, and in no context with two, are merged, unless one of them is undecided: equivalent in this sense to two values that some context distinguishes. A class that would then hold two distinguished values is left unmerged. The features are gone through twice, since what is merged in one makes more situations comparable in the others.

Consolidation then compares again. A value was merged while the other features were not yet merged, and it could be compared with few. With the classes of every feature at hand it is compared with every class, and if it is equivalent to a class not its own and never distinguished from it, while that class and its own are distinguished, it was undecided: it is taken out of its class and merged with nothing. This is repeated until nothing changes. In binary addition it takes out two last acts that had been merged with the state of no carry and are as much like the beginning of a sum (Figure 2c). A value that is not merged keeps the entries of the table that it has met. An undecided value is given, besides, an entry in every context in which all the classes it is equivalent to have one and the same answer; such entries are inferred, and they are not counted in the length of the program. Where the classes differ, or one of them has no answer, the program says that it does not know. A recall whose values the table never needs to distinguish, one class, is dropped, and what the table does not use is forgotten.

**Second causes** (line 10). If no set of features leaves the shorter strings without contradictions, the last act may hide a second cause beside it. The learner then counts as a dependence every difference at which no recall returns the place that differs, looks for recalls for these as above, and returns to the thresholds. Times three, among the withheld tasks, needs such a second cause (Appendix D.6).

**The test** (line 11). A program is announced only if, found from the shorter strings, it answers the longest: it is right wherever it speaks on them, and it says on them that it does not know no more often than the situations it has met once foretell, by the estimate of Good (1953), plus four standard deviations. What the longest strings add is then entered in the table. Finally the program is shorter than what it explains: the bits of its table (log_2_ of the number of different answers for each entry stated) and of its features (log_2_ of the number of relations counted for each recall, two bits for each other feature), plus the margin, are fewer than the bits of the answers it accounts for.

##### Cost

Each step is a count or an elimination, and no step grows faster than the product of the number of dependences and the number of relations counted. From the standard experience a task that the learner solves takes between 0.1 and 56 seconds of one processor core, with a median of 0.5; the slowest is multiplication, for which the learner asks a median of 990 questions (Appendix G gives the run times of the experiments).

#### C.3. Inner speech

When the answers of a task do not show what must be carried, a single answer at the end being the extreme case, the last act carries no state, and the learner in step ends with two answers in one situation. The learner then has to say something to itself after every symbol (Theorem 3(ii)). The module dm/inner.py learns what, by the same consolidation as the table, applied to moments.

##### The tree of moments

Every experience shorter than the longest is filed as a path from a common root, one moment for each beginning, and a moment at which an experience ends holds its answer. To merge two moments is to merge all that follows from them, since the program is deterministic: the merge is carried through the two subtrees, and it fails if two different answers meet. A merge is *supported* by every pair of equal answers that it brings together, one from each side; this counts the continuations that show the two moments equivalent. Every change is written down, so that a merge that fails can be taken back.

##### Growing the states

States are grown from the root, breadth first. A moment that follows a state is compared with every state found so far by trying the merge and following its consequences: the state is refuted if two different answers meet, and supports the moment if an answer that followed the moment met an equal answer that did not. Five rules are applied in this order, and after each change everything is compared again.

1. A moment that exactly one state supports, more than once, is that state, the best supported first.
2. Two states that support the same moment and do not contradict each other are one.
3. A moment that no state supports is a new state, the shortest first.
4. A moment that exactly one state supports, once, is that state.
5. Otherwise several states support the moment and contradict each other. Experience does not decide, and the moment is left undecided: it is identified with no state, no moment is identified with it, what follows it is followed like what follows a state, and it is compared again after every change, since a later merge may refute all its candidates but one.

Growing stops, and the learner reports that it has no program, if the table needs more states than the experience can bear: a quarter of the number of strings it learns from, and at least sixteen.

##### Comparing again

A table grown in this way has compared each moment with the states found until then, and a state found later may be as much like it. So when the table is complete, every step of it is compared again, with every state. One sweep backward through each experience gives how the experience would have ended had a moment of it been any other state. A step is undecided if some other state answers a continuation of its moments as experience did, and none otherwise, while that state and the state the step leads to are distinguished; the same holds for the event as which a finished bracketed part is heard. The moments of an undecided step were undecided. If the states they could be go on alike after every symbol and differ only in their answers, nothing about what follows is undecided: the step is led to a new state that keeps their steps and has no answer of its own, so that the program says that it does not know only if a string ends there, and a learner that can ask asks how such a string ends. Otherwise the table is grown again with these moments left undecided, and so on, until nothing undecided is found. The table that is announced holds no step that rests on the order in which its states were found; a table in which such a step cannot be left undecided is not announced.

##### Questions

A moment still undecided at the end is a question (Theorem 3(iii)). If the learner may ask, it takes two of the candidate states and looks through its experiences for a place where one of them stood and from which the program, as far as it is known, would end differently for the other. It puts what it heard up to the undecided moment in that place, at the beginning of the experience or of the bracketed part in which the state stood, and asks the world how this ends (Appendix A.7 follows one question). A question is shorter than the longest strings of the experience, which are kept for the test: a program has not predicted what it asked. The world answers only what it could itself have posed, so the learner asks first what is most like what it has heard: the moment in a place of the kind in which it was met, within a bracketed part or not, and going on as it has been heard to go on. It asks about the undecided moments that follow a state before those that follow another undecided moment, which wait for its answer. The answer drops every candidate it contradicts. When one candidate is left, or none, or the world has answered none of eight such questions, the learner turns to the next moment; the answers are added to the experience, and the states are grown again. If the learner may not ask, the moment stays undecided. The states and their steps are the program: *step*[state][symbol] *→* state and *answer*[state]. The procedure is that of the state-merging learners of regular languages (Oncina & García, 1992; Lang et al., 1998), with evidence required for a merge and with the same test of prediction as every other part of the learner.

##### The depth counter

Some single answers need a memory that is not finite: an arithmetic expression with brackets has to set aside what it has computed when a bracket opens. The learner finds the counter for this as it finds every counter, by solving. A counter that every experience returns to zero has gains that annihilate the numbers of symbols at the end of every experience, their null space; if such a counter, with gain +1 for one symbol and *−*1 for another, never falls below zero and reaches two, it is a depth. For bracketed arithmetic the null space is one vector, +1 for the opening bracket and *−*1 for the closing one. At an opening bracket the present state is filed and a new experience begins in the first state. At the closing bracket the depth counter stands again where the state was filed, the state is recalled, and it hears the finished part as one event. Which event is learned by the same merging: an event is one with a symbol, or with another event, if experience shows them equivalent in a common context and never distinguished. In the program for bracketed arithmetic a part that ends in the state of value *v* is heard as the digit *v* (Appendix C.6). The store of states is a recall of the depth counter, of the same form as the recall of the stack. A machine whose stack moves as its input dictates is a visibly pushdown automaton (Alur & Madhusudan, 2004); here the learner solves for the symbols that push and pop.

##### Announcing, and the order of questions

A program of inner speech is found from the shorter experiences, and it is a candidate if it answers all of them and every one of the longest. A candidate that does so with moments still undecided is held back while the other ways are gone (Appendix C.4): of two candidates the learner announces the one that leaves nothing undecided. The learner does all that it can with what it has heard before it asks, since a question is an act. Every way to a program is first gone on what was heard, and only if none gives a program that leaves nothing undecided does a learner that can act ask. If it holds programs with moments undecided, it asks what they leave undecided, beginning with the program that leaves least, and announces a program as soon as one leaves nothing undecided; a program that does not survive what the world answered is dropped. If it holds no program, it asks its way through inner speech, trial and recursion in turn until it holds one. Of the programs it holds in the end it announces the one that leaves least undecided, and the program says, of a string that meets an undecided moment, that it does not know. At test a step never taken is a situation never met, and there too the program says that it does not know.

#### C.4. Composition, trial and recursion

The module dm/closure.py turns the same learner on experience that the learner makes for itself (Theorem 5; Appendix B.9). Each of the three ways runs dm.learn on a derived experience, and each composite program is kept only if, by itself, it answers every experience it was built from, the longest included.

##### Composition

For each program of the library that answers the questions of the experience, its answer to each question is heard as further input, as symbols of a kind of their own, and the learner in step is run on what was heard; the first program of the library after which a program in step is found is kept. The counters S|b stand still while the learner hears its own answers, and W counts the symbols heard from the world. A sum said highest digit first is learned in this way: after its own sum, said lowest digit first, the learner reads its answer backward (Proposition 7).

##### Trial

For each symbol that occurs exactly once in every question longer than the two shortest lengths, the answers are put in its place, which turns each question into a statement. The learner, in step and then by inner speech, is asked for a program that predicts the last symbol of a statement from the rest. A symbol for which there is one is the unknown; of several, the one with the smallest program. A question is then answered by trying each answer in the place of the unknown and keeping the one under which the prediction comes true; if no answer or more than one does, the program says that it does not know. Here too the learner first tries the symbols on what was heard, and then asks only about the program it holds. An undecided moment of the prediction is a question about a statement, which the world does not answer; the learner puts it as a question of the task, hiding one symbol of the statement behind the unknown and trying each ending until the answer is the symbol it hid.

##### Recursion

The learner walks from each question in its memory down through its beginnings, the same question without its last symbol, and so on, recalling their answers or asking for them; a beginning that is no question of the task has the answer *nothing*, the end symbol alone. The difference between the answer to a string and the answer to its neighbour by deletion is taken by trial with a program of the library: it is the continuation that, put after the earlier answer, makes the program give the later one, found symbol by symbol. A symbol is kept if the program’s answer then agrees with the later answer at that place; if no symbol does, one is taken under which the answer does not yet reach that place, and the learner waits; at most 4,000 runs of the program are spent on one difference. From nothing the difference is the answer itself, and between equal answers it is nothing. The learner in step is run on the differences. If it finds a program for them, every answer is the answer before it plus a difference, and the answer to a question is the sum of the differences along its way (Appendix A.8). At least twelve differences are needed, and a program of the library under which more than two differences cannot be taken is passed over.

##### The shortest strings

A program of these three kinds may fail on strings of the two shortest lengths of the experience, which are often no questions at all: an equation too short to hold an unknown, a product too short to hold a multiplier. These strings the learner keeps in memory as they are, and answers from memory. Every longer string the composite program answers by itself.

#### C.5. The constants

The learner has four constants (dm.PARAMS), set once and used for every task (Table 8). Each is a level of evidence in the sense of a statistical test: it fixes how strong the evidence has to be before the learner keeps something, and it says nothing about what the learner should find.

Two of them are significance levels in the strict sense. *Luck*, four standard deviations, is the level at which a count is taken to exceed chance; for a count that is normal under chance it is a one-sided level of about 3*×* 10^*−*5^. The learner applies it when it keeps a recall, when it closes a filter, when it judges that two recalls meet as many dependences up to chance, and when it judges how often a program may say on the longest strings that it does not know. The *margin*, twenty bits, is the level at which a description is taken to explain more than it costs. Of all the strings of a given length *L*, at most a share 2^*−k*^ can be written *k* bits shorter, since fewer than 2^*L−k*^ descriptions are shorter than *L− k* bits; a saving of twenty bits therefore has odds of about a million to one against chance. The cost of a recall includes the bits that name it among all the relations counted, so the margin is a level already corrected for the number of hypotheses examined.

The other two are levels of effect: the smallest shares that count as a use. *Idle*, a tenth, is the share of its returns below which a kind of record may be closed out of a recall, and only if the kind is also needed less often than chance, by four standard deviations. *Major*, 15%, is the share of all differences above which a recall is part of every table; the other recalls are offered to the table, which uses them where it needs them. Experiment 5 sets each constant below and above its value, one at a time: under all eight settings the learner solves 28 of the thirty tasks, and 0 of its 2,400 runs announce a false program (Appendix D.6.1).

Four limits of size bound the search: at most eight recalls and twelve thresholds offered to the table, forty recalls compared in a round after counting, and 3,000 dependences sampled for a count. The programs announced in the experiments have at most three recalls and four thresholds. The vocabulary of Appendix C.1 bounds the gains of the counters solved for at three and the constants of the queries at two. A run of the learner is given 600 seconds of one processor core, of which inner speech may use 120.

#### C.6. The programs written for the benchmark

The listing below gives the programs of the benchmark as the learner prints them from the standard experience with seed 0 (paper/programs.txt, written by experiments/make numbers.py). The square root, for which the learner finds no program, is left out. The printed form keeps the names of the code; Table 9 translates it, and the readings after the listing say what each program does. Recalls and thresholds are numbered from 0, as in the listing.

**The programs of the benchmark, as printed by the learner** (standard experience, seed 0).

|  |
|---|
| == Even pairs (in step; 461 strings of at most 12 symbols) |
| read 0: I(s) = 1 |
| read 1: I(s) = I |
| law over [’read0’, ’read1’]: 4 situations; classes [2, 2] |
| == Modular arithmetic (no program from the standard experience; 565 strings of at most 12 symbols) |
| == Parity (inner speech; 461 strings of at most 12 symbols) |
| LAW: 2 states, 0 events that are no input symbol; no depth clock; 461 episodes of length <= 12, 0 moments left open, least used step taken in 300 episodes |
| state 0 after (start) answer 0 steps 0>0 1>1 |
| state 1 after 1 answer 1 steps 0>1 1>0 |
| == Cycle navigation (inner speech; 583 strings of at most 12 symbols) |
| LAW: 5 states, 0 events that are no input symbol; no depth clock; 583 episodes of length <= 12, 0 moments left open, least used step taken in 98 episodes |
| state 0 after (start) answer 0 steps 0>1 1>0 2>2 |
| state 1 after 0 answer 4 steps 0>3 1>1 2>0 |
| state 2 after 2 answer 1 steps 0>0 1>2 2>4 |
| state 3 after 0 0 answer 3 steps 0>4 1>3 2>1 |
| state 4 after 2 2 answer 2 steps 0>2 1>4 2>3 |
| == Stack manipulation (in step; 587 strings of at most 12 symbols) |
| read 0: N[+0+1-2+3+4](s) = N[+0+1-2+3+4](end) -A +1 [records with symbol in [0, 1, 3, 4]] |
| law over [’read0’]: 3 situations; classes [3] |
| == Reverse string (in step; 461 strings of at most 12 symbols) |
| read 0: I(s) = I -A +1 |
| law over [’read0’]: 2 situations; classes [2] |
| == Bracketed arithmetic (inner speech; 449 strings of at most 12 symbols) |
| LAW: 10 states, 0 events that are no input symbol; depth clock: +1 at symbol 8, -1 at symbol 9; 449 episodes of length <= 12, 0 moments left open, least used step taken in 2 episodes |
| state 0 after (start) answer 0 closes as 0 steps 0>0 1>1 2>2 3>3 4>4 5>0 6>5 |
| state 1 after 1 answer 1 closes as 1 steps 0>1 1>2 2>3 3>4 4>0 5>1 6>6 |
| state 2 after 2 answer 2 closes as 2 steps 0>2 1>3 2>4 3>0 4>1 5>2 6>7 |
| state 3 after 3 answer 3 closes as 3 steps 0>3 1>4 2>0 3>1 4>2 5>3 6>8 |
| state 4 after 4 answer 4 closes as 4 steps 0>4 1>0 2>1 3>2 4>3 5>4 6>9 |
| state 5 after 6 answer None steps 0>0 1>4 2>3 3>2 4>1 6>0 |
| state 6 after 1 6 answer None steps 0>1 1>0 2>4 3>3 4>2 6>1 |
| state 7 after 2 6 answer None steps 0>2 1>1 2>0 3>4 4>3 6>2 |
| state 8 after 3 6 answer None steps 0>3 1>2 2>1 3>0 4>4 6>3 |
| state 9 after 4 6 answer None steps 0>4 1>3 2>2 3>1 4>0 6>4 |
| == Solve equation (trial; 373 strings of at most 12 symbols) |
| trial: symbol 9 stands for the answer; each of [0, 1, 2, 3, 4] is tried in its place and the one is kept under which the forecast of the last symbol comes true |
| the forecast: |
| LAW: 15 states, 0 events that are no input symbol; depth clock: +1 at symbol 7, -1 at symbol 8; input symbols that are one event: {’5’: ’0’}; 334 episodes of length <= 11, 0 moments left open, least used step taken in 3 episodes |
| state 0 after (start) answer None closes as 0 steps 0>0 1>1 2>2 3>3 4>4 6>5 10>6 |
| state 1 after 1 answer None closes as 1 steps 0>1 1>2 2>3 3>4 4>0 6>7 10>8 |
| state 2 after 2 answer None closes as 2 steps 0>2 1>3 2>4 3>0 4>1 6>9 10>10 |
| state 3 after 3 answer None closes as 3 steps 0>3 1>4 2>0 3>1 4>2 6>11 10>12 |
| state 4 after 4 answer None closes as 4 steps 0>4 1>0 2>1 3>2 4>3 6>13 10>14 |
| state 5 after 6 answer None steps 0>0 1>4 2>3 3>2 4>1 6>0 |
| state 6 after 10 answer 0 steps |
| state 7 after 1 6 answer None steps 0>1 1>0 2>4 3>3 4>2 6>1 |
| state 8 after 1 10 answer 1 steps |
| state 9 after 2 6 answer None steps 0>2 1>1 2>0 3>4 4>3 6>2 |
| state 10 after 2 10 answer 2 steps |
| state 11 after 3 6 answer None steps 0>3 1>2 2>1 3>0 4>4 6>3 |
| state 12 after 3 10 answer 3 steps |
| state 13 after 4 6 answer None steps 0>4 1>3 2>2 3>1 4>0 6>4 |
| state 14 after 4 10 answer 4 steps |
| == Duplicate string (in step; 461 strings of at most 12 symbols) |
| read 0: I(s) = A |
| read 1: I(s) = -I +A |
| law over [’read0’, ’read1’]: 4 situations; classes [3, 3] |
| == Missing duplicate (in step; 385 strings of at most 12 symbols) |
| read 0: S\|2(s) = -S\|2(end) +I -E [records with symbol in [0, 1], seen 2 in [0]] |
| read 1: S\|2(s) = E [records with symbol in [0, 1], seen 2 in [1]] |
| law over [’read0’, ’read1’]: 3 situations; classes [2, 2] |
| == Odds first (in step; 461 strings of at most 12 symbols) |
| read 0: I(s) = 2A -2E -1 |
| read 1: I(s) = 2A |
| law over [’read0’, ’read1’]: 4 situations; classes [3, 3] |
| == Binary addition (in step; 441 strings of at most 12 symbols) |
| read 0: I(s) = A [records with seen 2 in [0]] |
| read 1: S\|2(s) = A [records with symbol in [0, 1], seen 1 in [1], seen 2 in [1]] |
| clock D0=1: turned whenever read 0 returns 1 |
| clock D1=1: turned whenever read 1 returns 1 |
| boundary 0: sign of N1 -D0=1 -D1=1 |
| law over [’read0’, ’read1’, ’last answer’, ’last read0’, ’last read1’, ’boundary0’]: 36 situations; classes [2, 2, 2, 6], of which open [0, 0, 0, 3]; 8 more situations answered for an open value |
| == Binary multiplication (recursion; 441 strings of at most 12 symbols) |
| recursion: answer(x) = binary_addition( answer(x without its last symbol), difference(x) ); the law of the differences: |
| read 0: S\|2(s) = S\|2(end) -A +2H +1 [records with symbol in [0, 1], seen 2 in [1]] |
| read 1: S\|2(s) = -S\|2(end) +A +1 [records with symbol in [0, 1], seen 2 in [0]] |
| clock D0=0: turned whenever read 0 returns 0 |
| clock D1=0: turned whenever read 1 returns 0 |
| clock D1=1: turned whenever read 1 returns 1 |
| count T0=0: the records holding 0 that read 0 can return |
| count T0=1: the records holding 1 that read 0 can return |
| boundary 0: sign of N2 +A -D1=0 -D1=1 -T0=0 -T0=1 |
| boundary 1: sign of N1 -D1=1 -T0=1 |
| boundary 2: sign of D0=0 |
| law over [’boundary0’, ’boundary1’, ’boundary2’, ’read1’]: 11 situations; classes [2, 2, 2, 2] |
| == Bucket sort (in step; 640 strings of at most 12 symbols) |
| boundary 0: sign of -N0 -N1 -N2 -N3 +A |
| boundary 1: sign of -N0 +A -1 |
| boundary 2: sign of -N0 -N1 +A |
| boundary 3: sign of -N0 -N1 -N2 +A -1 |
| law over [’boundary0’, ’boundary1’, ’boundary2’, ’boundary3’]: 5 situations; classes [2, 2, 2, 2] |

##### Reading the programs

###### Even pairs

Recall 0, I(s) = 1, returns the first symbol and recall 1, I(s) = I, the last one, and the table of four situations says whether they differ. The number of places where neighbouring symbols differ is even exactly when the first and the last symbol are equal, so this task, which seems to need a count, needs two records.

###### Modular arithmetic

No program from the standard experience: its table of 50 states is not complete from so few strings (Appendix A.7; Experiment 4).

###### Parity

Two states of inner speech (Appendix A.5).

###### Cycle navigation

Five states, the positions on a cycle of five. The symbols 0, 1 and 2 move the position back by one, leave it, and move it on by one, and the answer is the position: state 1, first reached after 0, answers 4.

###### Stack manipulation

One recall on the counter of depth that the learner solved for, with a filter closed to pops (Appendix A.4).

###### Reverse string

One recall, I(s) = I -A +1 (Appendix A.3).

###### Bracketed arithmetic

Ten states of inner speech and the depth counter, +1 at the opening bracket (symbol 8) and *−*1 at the closing one (9). States 0 to 4 hold the value so far, from 0 to 4: a digit adds itself to it, a plus sign (5) leaves it as it is, and a bracketed part that ends in the state of value *v* is heard as the digit *v*. States 5 to 9 hold a value followed by a minus sign (6), and the next digit is subtracted from it.

###### SOLVE equation

Trial: symbol 9 stands for the unknown, and each digit from 0 to 4 is tried in its place. The program that predicts the last symbol of a statement has 15 states and a depth counter for the brackets (symbols 7 and 8). Its states 0 to 4 hold the value of the left side so far, as in bracketed arithmetic; the plus sign is heard as the same event as the digit 0, since both leave the value as it is; and the equals sign (10) leads from the state of value *v* to a state that answers *v*, the last symbol of a true statement.

###### Duplicate string

Recall 0, I(s) = A, returns symbol *j* at answer *j* for the first copy, and recall 1, I(s) = -I +A, returns symbol *j − n* for the second copy, *n* being the length of the string.

###### Missing duplicate

The string is a string of 0s and 1s written twice, with one symbol replaced by 2 and a 3 appended when the length is odd, and the answer is the symbol replaced. Recall 0, over the records before the 2, returns the symbol half a string before the 2, and recall 1, over the records after it, the symbol half a string after it; S|2(end) is the number of symbols after the 2, and E half the length. The query of recall 0 adds a counter that the 2 resets to counters that it does not reset. Among the programs in step of the benchmark it is the one recall that needs, beyond Eq. (1), a counter set at an event to where another counter stands (Lemma 2), and the one that does not compile into rings and place cells alone (Appendix D.8).

###### Odds first

Recall 1, I(s) = 2A, returns the symbols at the places 2, 4, 6, … for the first half of the answers, and recall 0, I(s) = 2A -2E -1, returns those at the places 1, 3, 5, … for the second half, E being half the length, rounded down. The places are counted from one; counted from zero, the first half holds the odd places that give the task its name.

###### Binary addition

Two recalls, one for each number; the threshold of the 1s heard and not yet recalled; and the state carried by the last answer and the last two digits recalled, three states and 3 undecided values, for which the table infers eight entries (Appendix A.6).

###### Binary multiplication

Recursion with the learned addition. The program of the differences has two recalls, three thresholds and a table of 11 situations. Recall 1, S|2(s) = -S|2(end) +A +1 over the records of the multiplicand, returns the multiplicand shifted by one place less than the number of digits of the multiplier. Recall 0 returns the last digit of the multiplier at every second answer, and threshold 2, the side of the number of 0s that recall 0 has returned, is above zero exactly when that digit is 0; the difference is then nothing. Threshold 1, the 1s heard less the 1s that recall 1 has returned and the 1s of the multiplier, reaches zero when the 1s of the multiplicand have all been said, and the difference ends. Threshold 0 is one plus the number of answers at which recall 1 has returned nothing, less the number of digits of the multiplier; it stays at or below zero until the shifted multiplicand has been said, and so tells the zeros in front of it from the places after it, where recall 1 returns nothing as well (Appendix A.8).

###### Bucket sort

No recall and four thresholds. For each of the kinds 0 to 3, a threshold reports the side of “answers given, less the symbols heard of that kind and the kinds below it” (two of them at a starting value of *−*1, which moves the side by one), and the answer at a step is the smallest kind whose count the answers have not yet passed. The table has five situations, one for each answer.

### D. Experiments

This appendix gives the six experiments of Section 4, and Experiment 7 of Section 5.5, in full, in the same order and in the same four parts: why the experiment was run, its design, its result, and what it means. The design of each says what is needed to reproduce it: the tasks, the sizes of experience, the seeds, the variants of the learner, and the name of the experiment in the code. Appendix D.1 gives what the experiments share, and Table 11 gives every task with one question and its answers. The last two sections give Experiment 7: the engine in circuits of neurons (Appendix D.8) and the local rule of Theorem 6 run alone (Appendix D.9).

**Table 11.**
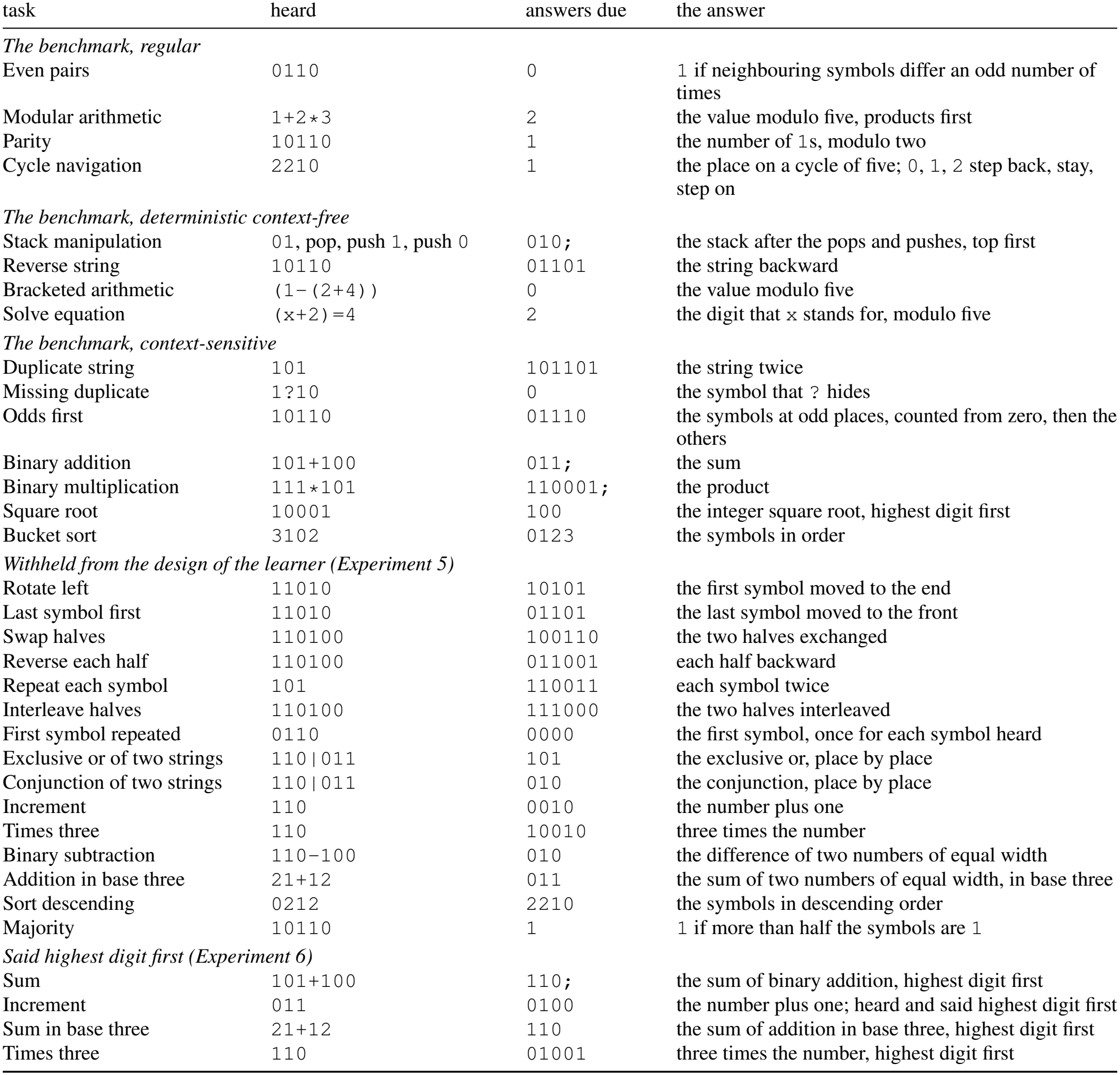
The tasks, each with one question and the answers due. Numbers are written lowest digit first unless the table says otherwise; ; is the end symbol of an answer whose length varies. The stack task hears an initial stack and then pops and pushes; in the missing duplicate, ? is the symbol that hides one place of a string written twice.

| task | heard | answers due | the answer |
|---|---|---|---|
| *The benchmark, regular* |  |  |  |
| Even pairs | 0110 | 0 | 1 if neighbouring symbols differ an odd number of times |
| Modular arithmetic | 1+2*3 | 2 | the value modulo five, products first |
| Parity | 10110 | 1 | the number of 1s, modulo two |
| Cycle navigation | 2210 | 1 | the place on a cycle of five; 0, 1, 2 step back, stay, step on |
| *The benchmark, deterministic context-free* |  |  |  |
| Stack manipulation | 01, pop, push 1, push 0 | 010; | the stack after the pops and pushes, top first |
| Reverse string | 10110 | 01101 | the string backward |
| Bracketed arithmetic | (1−(2+4)) | 0 | the value modulo five |
| Solve equation | (x+2)=4 | 2 | the digit that x stands for, modulo five |
| *The benchmark, context-sensitive* |  |  |  |
| Duplicate string | 101 | 101101 | the string twice |
| Missing duplicate | 1?10 | 0 | the symbol that ? hides |
| Odds first | 10110 | 01110 | the symbols at odd places, counted from zero, then the others |
| Binary addition | 101+100 | 011; | the sum |
| Binary multiplication | 111*101 | 110001; | the product |
| Square root | 10001 | 100 | the integer square root, highest digit first |
| Bucket sort | 3102 | 0123 | the symbols in order |
| *Withheld from the design of the learner (Experiment 5)* |  |  |  |
| Rotate left | 11010 | 10101 | the first symbol moved to the end |
| Last symbol first | 11010 | 01101 | the last symbol moved to the front |
| Swap halves | 110100 | 100110 | the two halves exchanged |
| Reverse each half | 110100 | 011001 | each half backward |
| Repeat each symbol | 101 | 110011 | each symbol twice |
| Interleave halves | 110100 | 111000 | the two halves interleaved |
| First symbol repeated | 0110 | 0000 | the first symbol, once for each symbol heard |
| Exclusive or of two strings | 110\|011 | 101 | the exclusive or, place by place |
| Conjunction of two strings | 110\|011 | 010 | the conjunction, place by place |
| Increment | 110 | 0010 | the number plus one |
| Times three | 110 | 10010 | three times the number |
| Binary subtraction | 110−100 | 010 | the difference of two numbers of equal width |
| Addition in base three | 21+12 | 011 | the sum of two numbers of equal width, in base three |
| Sort descending | 0212 | 2210 | the symbols in descending order |
| Majority | 10110 | 1 | 1 if more than half the symbols are 1 |
| *Said highest digit first (Experiment 6)* |  |  |  |
| Sum | 101+100 | 110; | the sum of binary addition, highest digit first |
| Increment | 011 | 0100 | the number plus one; heard and said highest digit first |
| Sum in base three | 21+12 | 110 | the sum of addition in base three, highest digit first |
| Times three | 110 | 01001 | three times the number, highest digit first |

**Table 12.**
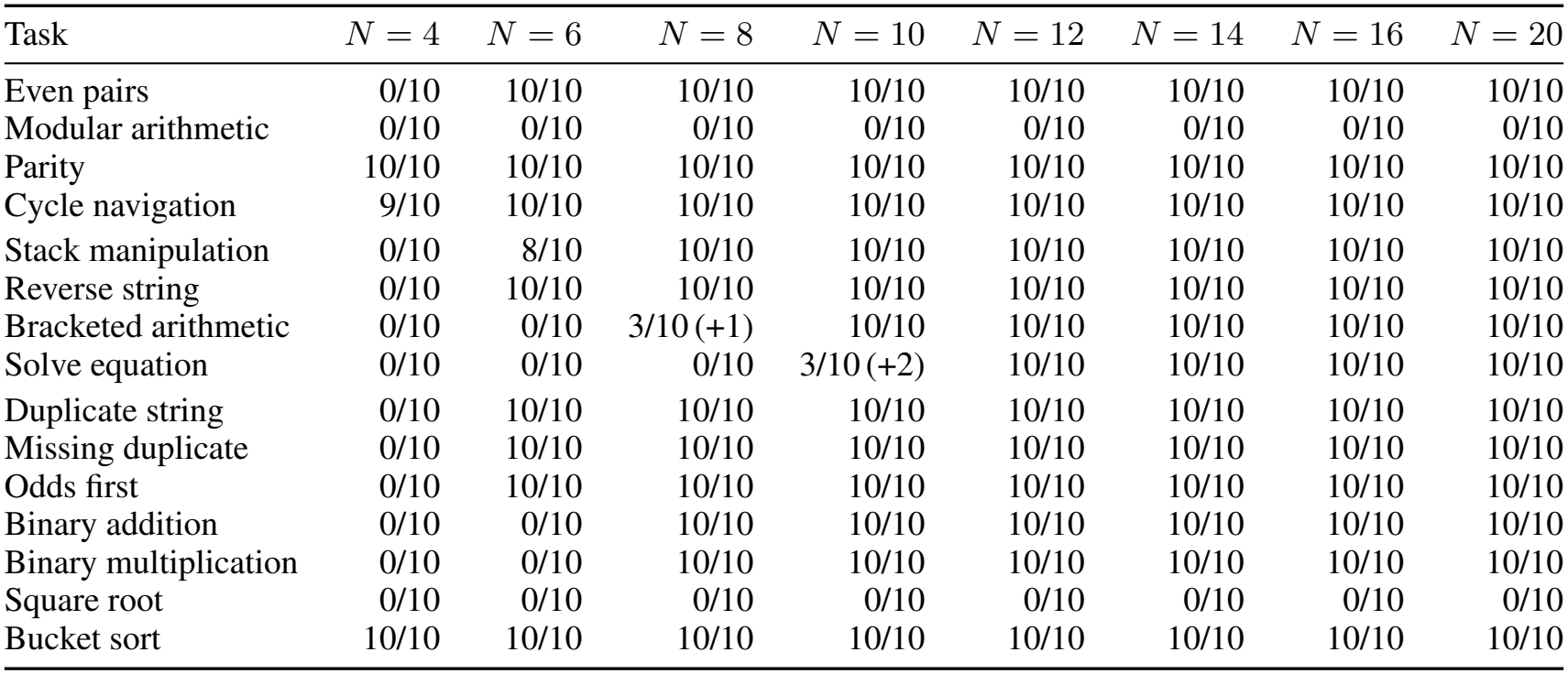
The benchmark at every size of experience: seeds exact of those run, for strings of at most *N* symbols (64 per length); (+*k*): *k* further seeds announced a program that was never wrong and abstained on some test string. Runs that announced a false program: 0.

| Task | *N* = 4 | *N* = 6 | *N* = 8 | *N* = 10 | *N* = 12 | *N* = 14 | *N* = 16 | *N* = 20 |
|---|---|---|---|---|---|---|---|---|
| Even pairs | 0/10 | 10/10 | 10/10 | 10/10 | 10/10 | 10/10 | 10/10 | 10/10 |
| Modular arithmetic | 0/10 | 0/10 | 0/10 | 0/10 | 0/10 | 0/10 | 0/10 | 0/10 |
| Parity | 10/10 | 10/10 | 10/10 | 10/10 | 10/10 | 10/10 | 10/10 | 10/10 |
| Cycle navigation | 9/10 | 10/10 | 10/10 | 10/10 | 10/10 | 10/10 | 10/10 | 10/10 |
| Stack manipulation | 0/10 | 8/10 | 10/10 | 10/10 | 10/10 | 10/10 | 10/10 | 10/10 |
| Reverse string | 0/10 | 10/10 | 10/10 | 10/10 | 10/10 | 10/10 | 10/10 | 10/10 |
| Bracketed arithmetic | 0/10 | 0/10 | 3/10 (+1) | 10/10 | 10/10 | 10/10 | 10/10 | 10/10 |
| Solve equation | 0/10 | 0/10 | 0/10 | 3/10 (+2) | 10/10 | 10/10 | 10/10 | 10/10 |
| Duplicate string | 0/10 | 10/10 | 10/10 | 10/10 | 10/10 | 10/10 | 10/10 | 10/10 |
| Missing duplicate | 0/10 | 10/10 | 10/10 | 10/10 | 10/10 | 10/10 | 10/10 | 10/10 |
| Odds first | 0/10 | 10/10 | 10/10 | 10/10 | 10/10 | 10/10 | 10/10 | 10/10 |
| Binary addition | 0/10 | 0/10 | 10/10 | 10/10 | 10/10 | 10/10 | 10/10 | 10/10 |
| Binary multiplication | 0/10 | 0/10 | 10/10 | 10/10 | 10/10 | 10/10 | 10/10 | 10/10 |
| Square root | 0/10 | 0/10 | 0/10 | 0/10 | 0/10 | 0/10 | 0/10 | 0/10 |
| Bucket sort | 10/10 | 10/10 | 10/10 | 10/10 | 10/10 | 10/10 | 10/10 | 10/10 |

Every experiment uses the one learner of Appendix C, dm.learn, with the same four constants for every task. It runs on one processor core, in Python with NumPy (Appendix G); the two gradient baselines are the only runs that use an accelerator. Every number in the text is read by its key from the results (paper/numbers.tex, written by code/experiments/make numbers.py; results/KEYS.md says which runs each key comes from). Table 10 lists the experiments with their names in the code, E1 to E9, B1 and B2, and the commands that run them.

#### D.1. Protocol

##### Tasks

A task gives, for each string it can be asked, the answers that are due. Table 11 gives the thirty tasks of the experiments, and the four of Experiment 6 said highest digit first, each with one question and its answers. The fifteen tasks of the benchmark are drawn by its generators as ported in the first paper (code/orrery p1/tasks.py). Their names follow the benchmark, with four exceptions: modular arithmetic is its Modular Arithmetic (Simple), bracketed arithmetic its Modular Arithmetic in the deterministic context-free group, parity its Parity Check, and the square root its Compute Sqrt. The tables group these tasks as the benchmark places them in the Chomsky hierarchy: regular, deterministic context-free, and context-sensitive. The fifteen withheld tasks are drawn by code/dm/tasks extra.py. Every task is also written as a function of the string (code/experiments/answers.py), which answers the learner’s questions; a test of the code checks it against the generator, symbol for symbol. A string is heard one symbol a moment, and its answers are due at the moments that follow it. Where an answer has no fixed length, as a sum has not, an end symbol follows it.

##### Experience

An experience of size *N* is drawn by the task’s generator: 64 strings of each length 1, …, *N*, with their answers, from which repeated strings are removed, so that all strings of the shortest lengths remain and a sample of the longer ones. The seed of a run decides which strings are drawn. The *standard experience* has *N* = 12: 768 strings are drawn, and after repetitions are removed a median of 457 distinct strings remain, from 363 to 653 over the tasks and seeds of Experiment 1. Experiments 1, 2 and 5 also use other sizes *N*, and Experiment 4 uses experiences with more strings of each length, written *N ×* strings per length.

##### Questions

Where the text allows it, the learner may ask a question: it chooses a string and is told its answers, or that the string is no question of the task. It asks in two places. A moment that consolidation leaves undecided is settled by a question (Section 3.4), and recursion needs the answers to the questions one symbol shorter than those it has heard (Theorem 5). The learner asks last: it first goes every way to a program on what it has heard, and asks only if none of them gives a program that leaves nothing undecided (Appendix C.3). Every question put to the world is counted, answered or not, and no question is as long as the longest strings of the experience, on which a program is tested before it is announced. Questions are allowed in every experiment except where the text withholds them: for the withheld tasks (Experiment 5); for the experience without minimal pairs (Experiment 4) and the learner of Experiment 2 that compares once, which receives the same experiences; under the lesion of questions (Experiment 3); and in the runs without questions of recursion (Experiment 6) and of arithmetic modulo five (Experiment 4).

##### The library

Composition and recursion run programs that the learner already has. A task of the benchmark is learned with a *library*: the programs in step that the same learner has written, from the same kind of experience (the same *N*, strings per length and seed), for the tasks before it in a fixed order (CURRICULUM in experiments/common.py, the tasks in step first). The library is built only when the learner reaches composition or recursion. A task said highest digit first (Experiment 6) has in its library the program of the same task said lowest digit first; the withheld tasks have none.

##### Time

A run of the learner has a budget of 600 seconds of processor time, of which each part of its search has a share. A part that reaches its share stops, and the record of the run says so. The record lists every way to a program that the learner went and why it gave none (log[‘tried’]): for instance, that the recalls leave two answers in one situation, or that no symbol of the question stands for its answer.

##### The test

An announced program is run on fresh strings of every length from 41 to 500, 32 strings of each length: 14,720 strings, drawn with the seed of the first paper’s test. In Experiments 1 and 5 a program that answers in step or by inner speech, and is exact on all of these, is run besides on four strings of each of 10^4^, 10^5^ and 10^6^ symbols; the other programs take a time that grows with the square of the length. The same program is often written again, from another experience or under another setting. It is tested once, and recognized by a fingerprint of all its counters, recalls and tables.

##### Four outcomes

On each test string a program gives the right answers; or it meets a situation that is not in its table, or an undecided moment, and says that it does not know; or it gives a wrong symbol without warning. A run is *exact* if its program answers every test string right. The program *abstains* if it answers no string wrongly and says, on some string, that it does not know; it is *false* if it answers some string wrongly without warning. A run ends with *no program* if the learner announces nothing, and the learner then gives the reason. A task is *solved* if every seed gives an exact program; there are ten seeds, 0 to 9, unless the text says otherwise. Beside these outcomes the benchmark’s own score is reported (Delétang et al., 2023): the share of the scored answer symbols that are right, averaged over the test lengths. Table 1 gives it for the learner and for the published networks, and Tables 13 and 14 for the two gradient baselines.

**Table 13.** Gradient descent on the learner’s experience: accuracy (%) at lengths 41 to 500 of networks trained on the strings of at most ten symbols that the learner heard (condition A; mean over seeds).

| Task | LSTM | Transformer |
|---|---|---|
| even pairs | 100.0 | 49.9 |
| modular arithmetic | 20.6 | 21.1 |
| parity check | 100.0 | 49.8 |
| cycle navigation | 23.0 | 20.8 |
| stack manipulation | 35.3 | 40.4 |
| reverse string | 53.4 | 50.6 |
| bracketed arithmetic | 18.8 | 20.3 |
| solve equation | 20.4 | 21.7 |
| duplicate string | 51.6 | 50.4 |
| missing duplicate | 97.6 | 52.5 |
| odds first | 50.9 | 50.4 |
| binary addition | 30.4 | 49.1 |
| binary multiplication | 32.8 | 47.5 |
| square root | 51.2 | 50.0 |
| bucket sort | 28.9 | 23.4 |

**Table 14.**
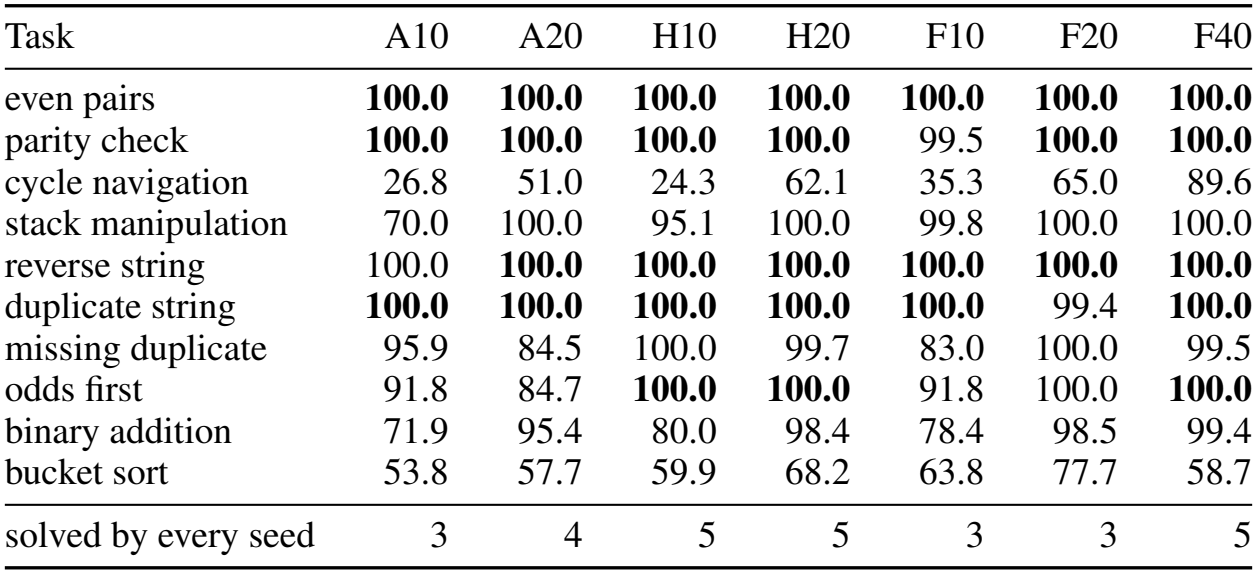
The same machine, trained by gradient descent through time: accuracy (%) at lengths 41 to 500 of the first paper’s machine (mean over seeds; bold: solved by every seed). A*N* : trained on the strings the learner heard, of at most *N* symbols; H*N* : the same, with a fifth of them set aside for validation; F*N* : on fresh strings of at most *N* symbols at every step (F40: the first paper’s protocol).

| Task | A10 | A20 | H10 | H20 | F10 | F20 | F40 |
|---|---|---|---|---|---|---|---|
| even pairs | **100.0** | **100.0** | **100.0** | **100.0** | **100.0** | **100.0** | **100.0** |
| parity check | **100.0** | **100.0** | **100.0** | **100.0** | 99.5 | **100.0** | **100.0** |
| cycle navigation | 26.8 | 51.0 | 24.3 | 62.1 | 35.3 | 65.0 | 89.6 |
| stack manipulation | 70.0 | 100.0 | 95.1 | 100.0 | 99.8 | 100.0 | 100.0 |
| reverse string | 100.0 | **100.0** | **100.0** | **100.0** | **100.0** | **100.0** | **100.0** |
| duplicate string | **100.0** | **100.0** | **100.0** | **100.0** | **100.0** | 99.4 | **100.0** |
| missing duplicate | 95.9 | 84.5 | 100.0 | 99.7 | 83.0 | 100.0 | 99.5 |
| odds first | 91.8 | 84.7 | **100.0** | **100.0** | 91.8 | 100.0 | **100.0** |
| binary addition | 71.9 | 95.4 | 80.0 | 98.4 | 78.4 | 98.5 | 99.4 |
| bucket sort | 53.8 | 57.7 | 59.9 | 68.2 | 63.8 | 77.7 | 58.7 |
| solved by every seed | 3 | 4 | 5 | 5 | 3 | 3 | 5 |

#### D.2. Experiment 1: the benchmark without backpropagation

##### Why

The benchmark of Delétang et al. (2023) asks whether a procedure learned from strings of up to 40 symbols holds on strings of 41 to 500. Networks trained by gradient descent fail on most of its tasks unless they are given a stack or a tape (Table 1). The first paper’s machine runs programs for these tasks in the memory of Eq. (1), and it found them by gradient descent through time from hundreds of thousands of strings (He & Wu, 2026a). If programs are written by comparison, the same programs should be written from a few hundred short strings, by comparing, solving and counting.

**Design** (E1; make e1). The fifteen tasks of the benchmark (Table 11). Experiences of *N∈ {*4, 6, 8, 10, 12, 14, 16, 20*}*, 64 strings per length; 10 seeds; library and questions as in the protocol: 1,200 runs. Beside the learner, Table 1 gives the published accuracies of networks trained by the benchmark’s protocol on strings of up to 40 symbols: LSTM and Tape-RNN, the best of 30 runs (Delétang et al., 2023); the Transformer, the best of the eleven positional encodings of Ruoss et al. (2023); and the first paper’s machine, the mean of three seeds (He & Wu, 2026a). The LSTM, Transformers and Tape-RNN were trained on at least 10^6^ batches of 128 fresh strings. Two baselines train networks by gradient descent on the learner’s own strings: an LSTM and two Transformers (Appendix D.2.1), and the first paper’s machine itself (Appendix D.2.2).

##### Result

Tables 1 and 12, and Figure 3a. From the standard experience thirteen tasks are exact for every seed, each learned in 0.1 to 56 seconds of one processor core, with a median of 0.5. Of the 1,200 runs of the experiment, none announced a false program, and 3 a program that was never wrong and abstained on some test string, where a situation of its table had not been met: bracketed arithmetic from *N* = 8; solve equation from *N* = 10 (2 seeds). The programs in step and by inner speech were exact at 10^6^ symbols wherever they were tested, those of eleven of the tasks solved. The square root is announced at no size. Arithmetic modulo five is announced at no size of this grid, since the table of its 50 states is not complete from so few strings; Appendix D.5.1 gives it more.

###### Questions

At the standard experience the learner asks nothing for eleven of the thirteen tasks it solves. For multiplication it asks a median of 990 questions, and at most 997 (Experiment 6). For equations, questions are asked in two of ten seeds, at most five questions. In such an experience a subtraction of a negative number is followed by one digit only; the moment after it is equivalent to two states of the forecast that are distinguished, and one statement settles it.

###### The same program

For five tasks every seed writes the same program, symbol for symbol, from its own experience (parity, cycle navigation, reverse string, duplicate string, odds first). For the others the programs differ in entries of the table that experience has or has not met. Appendix C.6 prints the programs of seed 0.

##### Reading

The tasks fall into the four ways in which the learner finds a program: in step, by inner speech, by trial and by recursion. Eight have a program in step, whose recalls are found from minimal pairs, whose thresholds are found from what the recalls leave unexplained, and whose state, where one is needed, is carried by the last answer: even pairs, the four tasks that copy (reversal, duplication, the missing duplicate and odds first), the stack, binary addition and sorting. Parity, cycle navigation and bracketed arithmetic, three tasks with one answer at the end, need inner speech; bracketed arithmetic needs a counter of depth as well, which the learner finds as the count that every experience brings back to zero. Equations need trial, and multiplication recursion. Neither has a program of the other kinds, and the learner reaches them only after it has found, for each earlier way, why that way gives no program (log[‘tried’] in the records). On the ten tasks given to both, from the same strings of at most ten symbols, the learner is exact for every seed on all ten and gradient descent through time on the same machine on three (Appendix D.2.2). The experience needed is small because each thing learned is fixed by few dependences (Theorem 2) and by few situations; and the programs hold at every length because they use coincidences alone (Theorem 4).

##### D.2.1. Baseline: networks trained on the learner’s strings

###### Why

The published scores of Table 1 are of networks trained on strings of up to forty symbols, drawn anew at every step. To see what gradient descent does with the learner’s own experience, an LSTM and two Transformers are trained on exactly the strings that the learner heard.

**Design** (B1; code/baselines/gradient baselines.py). An LSTM (Hochreiter & Schmidhuber, 1997) of one layer of 256 units, and Transformer encoders (Vaswani et al., 2017) of five layers, width 64 and eight heads, one with sinusoidal positions and one with rotary positions (Su et al., 2024); the better of the two Transformers is reported. As in the benchmark and in the first paper, a network reads the symbols of the question and then one placeholder for each answer symbol, and says an answer symbol at each placeholder; the loss is the cross-entropy at the scored answer positions, and training is by Adam with step size 10^*−*3^ and gradients clipped at norm one. *The learner’s strings* (condition A): the experience of the learner at *N* = 10, the distinct strings among 64 random strings of each length from 1 to 10, drawn with seeds 0, 1 and 2 of Experiment 1. A step draws a length and then 32 of the strings of that length, and training stops when every scored symbol of the training strings is right, or after 30,000 steps. *The benchmark’s fresh strings, with a tenth of its steps* (condition B): new random strings of a random length from 1 to 40 at every step, 10^5^ steps of batch 128 (12.8 million strings in a run), one step size and three seeds; the benchmark itself trains for 10^6^ steps and reports the best of 30 runs. Three seeds in each condition and fifteen tasks: 270 runs. The test is that of the learner at 24 of its lengths: 32 fresh strings of each of the lengths 41, 61, …, 481 and 500.

###### Result

Table 13. Every network fits the strings it was given. The LSTM then solves two tasks at lengths 41 to 500, both of them regular (even pairs and parity), and the Transformers solve no tasks. Trained on the benchmark’s fresh strings, with a tenth of its steps, the LSTM solves three tasks for every seed, and the Transformers solve no tasks.

###### Reading

What these networks take from a few hundred short strings is the strings themselves. Whether that is due to gradient descent or to the networks is the question of the second baseline, in which the network is the first paper’s machine.

##### D.2.2. Baseline: the first paper’s machine, trained by gradient descent

###### Why

The networks of the first baseline differ from the learner in what they are as well as in how they learn. The first paper’s machine differs from it only in how it learns: it is Eq. (1) with a recurrent controller, and gradient descent through time finds its programs (He & Wu, 2026a). Training it on the learner’s experience compares two ways of learning in one machine.

**Design** (B2; code/baselines/bptt first paper.py). The first paper’s machine (code/orrery p1), with its relaxation and schedule (Adam, 12,000 steps of 32 strings), its ladder of at most six restarts, stopped at the first whose validation reaches 99%, and its choice of checkpoint, all unchanged; it is tested as the exact discrete machine. Only the experience differs. *Same experience* (A*N* ): the strings the learner heard, of at most *N* = 10 or 20 symbols, and no others; a step draws 32 of them of one length, and the strings themselves serve for validation. *Part held out* (H*N* ): the same strings with a fifth of those of every length set aside; the machine is trained on the rest, and the checkpoint and the restarts are decided by the part set aside. *Fresh strings* (F*N* ): the first paper’s protocol, 384,000 new strings in a run, of at most *N* = 10, 20 or 40 symbols; F40 is the protocol of the first paper itself. The tasks are the ten of the benchmark that the learner solves from its standard experience without composition, trial or recursion: the eight in step, parity and cycle navigation. Three seeds, the test of the first baseline, and 165 hours on an accelerator in all.

###### Result

Table 14. On the learner’s strings of at most ten symbols, gradient descent solves 3 of the 10 tasks for every seed (15 of 30 runs are solved, and 22 of 30 fit their strings), and on its strings of at most twenty, 4; with a part held out, 5 and 5. On fresh strings of at most ten, twenty and forty symbols it solves 3, 3 and 5. No condition solves sorting, binary addition or the stack for every seed.

###### Reading

With one machine and one experience, the learner is exact on all ten tasks from strings of at most ten symbols (Table 12, *N* = 10), and gradient descent through time on three of them; fresh strings four times as long raise this to five. The difference lies in how the program is found.

#### D.3. Experiment 2: a learner that knows what it knows

##### Why

A learner that extrapolates can be wrong where nobody checks. By Theorem 4, a learner that merges only what experience has decided, and announces only what it has tested, should seldom announce a false program, however little it has heard. The test is made where experience is thin and the room for guessing is largest, and against three variants of the learner, each without one of the safeguards of part (iv) of the postulate: leaving undecided what experience has not decided, testing on the longest strings, and comparing again.

**Design** (E3 and E7; make e3, make e7). *The census*. Every run of the complete learner on the thirty tasks at 64 strings per length is classified: the runs of Experiments 1 and 5, and runs at the sizes between, *N* = 3, 5, 7, 9 and 11 (tag census).

These are thirteen sizes from *N* = 3 to 20, with ten seeds each: 3,900 runs. *Made to guess* (tag guess; the lesion open): consolidation merges values in the order of how often they were shown equivalent, whatever else they are equivalent to; a moment that several states support is given to the best supported, as learners that seek the fewest states do; and nothing is compared again. It is run on the experiences of the census at *N* = 3, 5, 7, 9 and 11, and on arithmetic modulo five in the larger experiences of Experiment 4: 1,580 runs, compared run for run with the complete learner on the same 1,580 experiences. *Untested* (tag untested; the lesion prediction): everything is found from all the strings, and the program is announced without the test on the longest strings; the experiences of the census at the same five sizes, 1,500 runs. *Single pass* (tag no twins once; the lesion look): the learner leaves undecided what experience has not decided, and what consolidation has merged, and each step of a table of inner speech, is not compared again when more is known. It is run on the standard experience without minimal pairs of Experiment 4, where inner speech has to carry tasks on thin evidence: 300 runs. All three variants are also run as lesions on the standard experience (Experiment 3). *All runs*. The census, the runs with questions of Experiment 4, the runs of Experiment 6, and the larger experiences of Experiment 4 with questions and without, are the 5,680 runs of the complete learner in this paper.

##### Result

Table 15 and Figure 3b. Of the 3,900 runs of the census, 2,742 gave an exact program, 6 a program that was never wrong and abstained on some test string, 1,152 no program, and none a false program. Of all 5,680 runs of the complete learner, none announced a false program; of the 4,157 programs it announced, 4,150 were exact and 7 abstained on some test string.

**Table 15.**
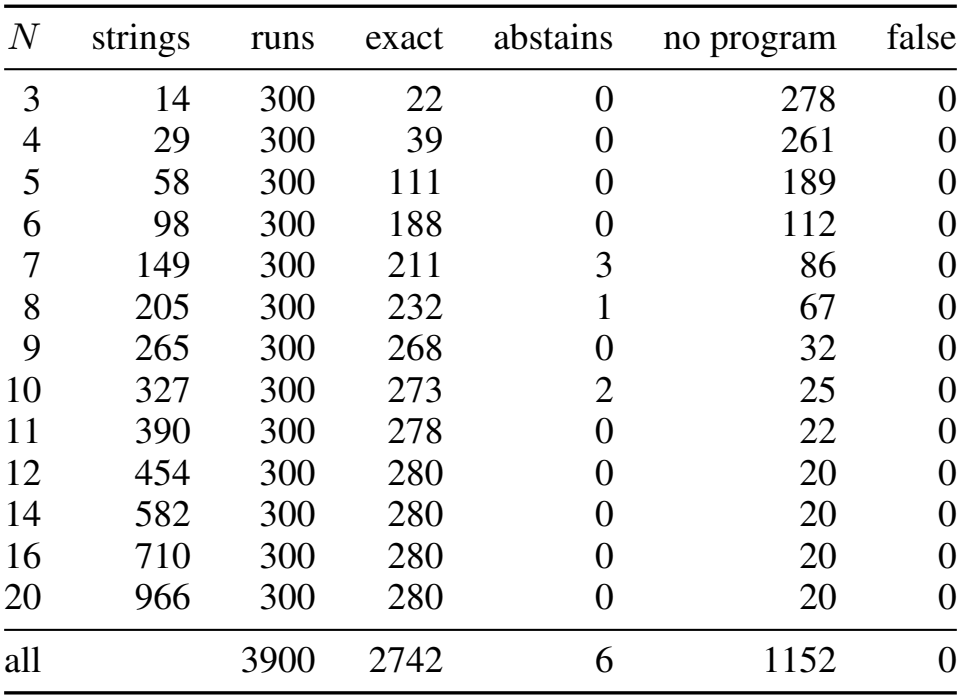
What the learner announces, over the thirty tasks and all seeds, against the longest string heard: runs whose program is exact, abstains on some test string, is false, and runs with no program (strings: the median number of strings in an experience).

| *N* | strings | runs | exact | abstains | no program | false |
|---|---|---|---|---|---|---|
| 3 | 14 | 300 | 22 | 0 | 278 | 0 |
| 4 | 29 | 300 | 39 | 0 | 261 | 0 |
| 5 | 58 | 300 | 111 | 0 | 189 | 0 |
| 6 | 98 | 300 | 188 | 0 | 112 | 0 |
| 7 | 149 | 300 | 211 | 3 | 86 | 0 |
| 8 | 205 | 300 | 232 | 1 | 67 | 0 |
| 9 | 265 | 300 | 268 | 0 | 32 | 0 |
| 10 | 327 | 300 | 273 | 2 | 25 | 0 |
| 11 | 390 | 300 | 278 | 0 | 22 | 0 |
| 12 | 454 | 300 | 280 | 0 | 20 | 0 |
| 14 | 582 | 300 | 280 | 0 | 20 | 0 |
| 16 | 710 | 300 | 280 | 0 | 20 | 0 |
| 20 | 966 | 300 | 280 | 0 | 20 | 0 |
| all |  | 3900 | 2742 | 6 | 1152 | 0 |

Made to guess, the learner announced three false programs in 1,580 runs, by task and experience: binary addition (*N* = 7: 2); modular arithmetic (16 *×*512: 1). On the same experiences the complete learner announced none. The programs of the learner made to guess were exact in 912 runs and abstained in 2; those of the complete learner, in 930 and 4. Untested, the learner announced fourteen false programs in 1,500 runs: majority (*N* = 3: 10); modular arithmetic (*N* = 3: 1); bracketed arithmetic (*N* = 7: 1); solve equation (*N* = 7: 1); addition in base three (*N* = 5: 1). It announced 1,348 programs where the complete learner announced 893, of which none were false. In a single pass, on the experiences without minimal pairs, the learner solved 6 of the thirty tasks, as many as the complete learner solved there (6), and it announced one false program in 300 runs, for the majority of a string, where the complete learner announced none.

##### Reading

Each false program of a variant answers every string it heard, and the longest strings, correctly, and errs beyond them, through the safeguard that the variant lacks (Appendix D.3.1). The complete learner falls short of an exact program in two ways, and says so in both. At the smallest sizes it mostly reports that it has no program, and gives the reason. And a program it announces is sometimes incomplete; the program then says, of a test string that leads it to a situation it has not met, that it does not know. The test on the longest strings has a cost, which the untested runs show: the longest strings are kept from the learner, and with them the larger part of a short experience, so that at these sizes the complete learner announces fewer programs than a learner that tests nothing. The test gives in return the bound of Theorem 4, which holds for a program however it was found. Over all experiments, comparing again took a value out of its class, or made a step of inner speech undecided, in 135 of the 4,157 programs that the complete learner announced (binary addition: 128; solve equation: 4; bracketed arithmetic: 2; modular arithmetic: 1).

##### D.3.1. What a guess looks like

A learner guesses wherever it chooses between possibilities that experience has left equal (Section 3.5). The false programs of the variants show three kinds of guess: a value merged with one of two classes, a moment given to one of several states, and an early decision that is not compared again.

###### A value: addition from 126 strings

From 126 strings of at most 7 symbols (seed 3), the learner made to guess writes a program for binary addition that has the two recalls, the threshold and the carry of the true one, that answers every string it has heard, and that predicts the 50 longest. One merge in it is a guess. The benchmark writes its numbers with leading zeros, and the last act “said zero, recalled zero and zero” had been met only where digits were still to come. There it is equivalent to two classes of last act that are distinguished: the first moment, after which “nothing left to add” means the number zero, and an ordinary digit, after which it means the end of the sum. The learner made to guess merges it with the class it was more often shown equivalent to, the first moment, and two more last acts follow it there. Its program then says 0 where a sum ends in a carry out of two such zeros (as in 60534 + 225329, lowest digit first, written with twenty and twenty-one digits), on eighteen of the 14,720 test strings. The complete learner leaves that last act undecided and merges it with nothing. After it, the program answers where the first moment and an ordinary digit would give the same answer, and nowhere else; it answers this sum and every test string. Of the ten experiences of this size, one for each seed, the complete learner made nine exact programs and one that was never wrong and abstained on eleven test strings; the learner made to guess made six exact programs, two false ones, and no program in two.

###### A moment: arithmetic modulo five, settled by one question

In one experience of arithmetic modulo five (seed 8 at 16 *×*512; code/demo.py runs it) nothing heard continues the beginning 1+1*×* 2*×* 2 with a product. The beginning is equivalent to five states that are distinguished, those after 0, 1+4, 2+3, 0 *−*2+2 and 4+1: all have the value zero, and they differ in the product still in progress. The learner made to guess takes the first of five equally supported candidates. Its table of 50 states has a wrong entry, which the 1,024 longest strings do not reach, and one test string in thirteen is answered wrongly. The complete learner asks what 1+1 *×*2 *×*2 *×*2 is. The answer, four, leaves the state after 1+4, and the table is exact (Appendix A.7). The question is a neighbour made on purpose: the beginning, continued by *×*2, after which each of the five candidates would give a different answer.

###### An early decision: the counter of majority

The third kind needs no tie between candidates: a decision that experience supported when it was taken, and that is not compared again when a state found later turns out to be equivalent as well. The majority of a string is one threshold, the side of “ones less zeros”, which the complete learner solves for from the dependences that minimal pairs show. Without minimal pairs (Experiment 4) there are no dependences, and the learner falls back on inner speech, which can grow a counter that stops counting: nine states, which count the ones ahead, or the zeros ahead, only a few steps far, the last state returning to itself whatever is heard. This table answers every string heard, the longest among them; the shortest string that it answers wrongly has eight symbols, and was not heard. The complete learner does not announce it. The step into the last state was given a state of its own while the table was grown; the finished table shows its moments to be equivalent as well to another state, from which the last one is distinguished, so experience has not decided where the count ends. Comparing again makes the step undecided, one of the longest strings is then not answered, and the learner reports that it has no program. Comparing only once, the learner announces it in one run, and it is wrong. A finite automaton fitted to a count is the error of recurrent networks on this benchmark; the learner avoids it by comparing again every decision it took early.

###### How fine the distinction is: even pairs

Two strings of one symbol form a minimal pair, so an experience without minimal pairs holds only one of them. The table of inner speech that the complete learner then announces for even pairs is exact, and it has one moment beside the states of the task: the string of one symbol that it did not hear, which continues as two of the states do and has no answer. In a single pass, the table gives that string the answer of the state to which it happened to be assigned, and for the experience with seed 0 that answer is wrong. A learner that may ask settles it with one question.

#### D.4. Experiment 3: what each part is for

##### Why

If the postulate describes how programs are learned, each of its parts should carry tasks that the others cannot reach, and the losses should be those that the theory names.

**Design** (E2; make e2). The thirty tasks, the standard experience, ten seeds. One part of the learner is switched off at a time; the name of the lesion in the code is given in brackets.

- *Comparison of minimal pairs* (difference): no minimal pair is compared, so no dependence is found.
- *Solving for key counters* (solve): no key counter is solved from dependences; the count of all symbols heard and the counters that a symbol resets remain.
- *Thresholds* (residues): no threshold is solved for, and no counter turned by what a recall returns.
- *Consolidation* (sleep): nothing is merged.
- *The test* (prediction): everything is found from all the strings, and the program is announced untested.
- *Leaving undecided* (open): the learner is made to guess, as in Experiment 2.
- *Comparing again* (look): a single pass, as in Experiment 2.
- *Inner speech* (inner), and *composition, trial and recursion* (closure).
- *Questions* (play): no question may be asked.

With the complete learner, these are eleven conditions of 300 runs each.

##### Result

Table 16 and Figure 3c. The complete learner solves 28 tasks, and so does each of the three variants of Experiment 2. Without comparison eight tasks remain, and each other lesion loses the tasks listed in the table. At this size of experience false programs appear under two lesions: without comparison the learner announces three false programs and without thresholds three.

**Table 16.** Lesions: tasks (of thirty) solved from the standard experience when one part is removed; runs that announced a false program; runs whose program abstained on some test string; and the tasks lost.

| Part removed | solved | false | abstains | tasks lost |
|---|---|---|---|---|
| nothing (the complete learner) | 28 | 0 | 0 | – |
| comparison of minimal pairs | 8 | 3 | 0 | conjunction of two strings, binary addition, binary multiplication, binary subtraction, duplicate string, first symbol repeated, and 14 more |
| solving for key counters | 27 | 0 | 0 | stack manipulation |
| thresholds | 23 | 3 | 0 | binary addition, binary multiplication, bucket sort, majority, sort descending |
| consolidation | 25 | 0 | 19 | binary addition, binary subtraction, addition in base three |
| the test (announced untested) | 28 | 0 | 0 | – |
| leaving undecided (made to guess) | 28 | 0 | 0 | – |
| comparing again (single pass) | 28 | 0 | 0 | – |
| inner speech | 24 | 0 | 0 | cycle navigation, bracketed arithmetic, parity, solve equation |
| composition, trial, recursion | 26 | 0 | 0 | binary multiplication, solve equation |
| questions | 26 | 0 | 2 | binary multiplication, solve equation |

##### Reading

Each part carries tasks that no other part reaches, and the losses are those that the theory names. Without comparison the programs that remain are those without a recall: thresholds alone (the two sorts) or inner speech alone.

Without solving, the stack is lost, since its counter of depth is on no list. Without thresholds addition and the sorts are lost, whose ends and orders are thresholds, and with them multiplication, whose program of differences has 3 thresholds, and majority, whose program is one. Without consolidation the tables of the sums (binary addition and subtraction, and addition in base three) are not complete from the standard experience, and the learner says so: their programs abstain on the test strings that reach a missing entry. Without inner speech parity, cycle navigation and bracketed arithmetic are lost, and equations, whose forecast is a table of inner speech. Without composition, trial and recursion, multiplication and equations are lost. Without questions, multiplication is lost, since it needs the answers to shorter questions, and so are equations for the seeds whose forecast has an undecided moment; their programs abstain. The false programs without comparison and without thresholds are of one kind, and come from the same three experiences under either lesion. The majority of a string needs a threshold, the side of “ones less zeros”; without it the task is left to inner speech, and a learner that may ask settles, with 1 to 26 questions, a counter of nine or ten states, which counts no further than four or five ahead. It agrees with every string heard, every answer given and every one of the longest strings; the shortest string that it answers wrongly has nine or ten symbols, and was not among them. At twelve symbols, the length tested, it is wrong on 14 to 26 of the 4,096 strings, and 64 such strings drawn at random let it pass with probability 0.67 to 0.80: this is the bound of Theorem 4, which holds at the length tested and not beyond. The three variants of Experiment 2 lose nothing at this size of experience: they solve 28, 28 and 28 tasks, and none of them announces a false program, since little is left undecided at the standard experience, and a learner that does not guess leaves little for the longest strings to refute. Experiment 2 tests them where experience is thin.

#### D.5. Experiment 4: minimal pairs and questions

##### Why

The theory says that what teaches a dependence is the minimal pair. Removing the pairs from an experience should remove exactly the tasks that need a recall, and a learner that can ask should be able to make the pairs itself.

**Design** (E3; make e3). *Without minimal pairs* (tag no twins): the standard experience with its minimal pairs taken out. At every length the strings of the standard experience are taken in order, and a string is kept unless it differs in one symbol from a string already kept. Further strings are then drawn, from a second generator so that the standard draw is not disturbed, until the length has as many strings as before, or as many as it can hold without minimal pairs. The shortest lengths, of which the standard experience holds every string, keep at most half of them, since two strings of one symbol always form a pair: an experience of 454 strings keeps 388 (medians over tasks and seeds). The thirty tasks, ten seeds; no question may be asked. The learner that compares once receives the same experiences (Experiment 2). *As large, with its pairs*: the census experience whose median number of strings is nearest, *N* = 11 (Experiment 2). *With questions* (tag play *f* ): from the experience without minimal pairs, the learner changes one symbol in a share *f* of its experiences, *f* = 0.1, 0.25, 0.5 or 1, and asks for the answer, which gives the experience a minimal pair. A change that leaves the task is tried again elsewhere; if no change of one symbol is a question of the task that the learner has not heard, nothing is added. With *f* = 1 a pair was made for 99.6% of the experiences. *More strings*: arithmetic modulo five and three other tasks with more and longer strings (Appendix D.5.1).

##### Result

Without minimal pairs six of the thirty tasks are solved, and no run announces a false program; every run without an exact program reports that it has none (Figure 3d). The loss comes from the pairs: the census experience of *N* = 11, as large (390 strings) and with its pairs, gives 278 of 300 exact runs and solves 27 tasks, 26 of them without a question. With questions in a tenth of the experiences 20 tasks are solved, in a quarter 27, in half 28, and in all of them 28. No run with questions announced a false program; these runs are among the 5,680 of Experiment 2.

##### Reading

The tasks that need no recall do not need minimal pairs, and the others do; a learner that asks gets them back. The tasks solved without minimal pairs are those whose programs are thresholds or inner speech alone: the two sorts, by thresholds, and even pairs, parity, cycle navigation and the missing duplicate, by inner speech (the missing symbol of a doubled string of bits is the parity of the symbols heard). For the others the learner reports that it has no program; for the majority of a string, the single pass of Experiment 2 shows what a learner announces that does not compare again (Appendix D.3.1). Each question asks for a neighbour made on purpose: here a minimal pair, made by substituting one symbol; for an undecided moment, a continuation that distinguishes two states, as in arithmetic modulo five below; and for recursion, the neighbour by deletion (Experiment 6). The standard experience is full of minimal pairs, and from it the learner asks only in the last two cases.

##### D.5.1. More experience, and questions

###### Why

Inner speech needs every step of its table to have been taken in the shorter strings of the experience, and every state to have been distinguished from every other. Arithmetic modulo five with precedence has 50 states over eight symbols, and the standard experience does not hold them all. The experiment asks how much experience the table needs, how much of it a few questions replace, and what a guess does in their place.

**Design** (E7; make e7). The two tasks of the benchmark that the standard experience leaves without a program, arithmetic modulo five and the square root, and for comparison bracketed arithmetic and equations, at the experiences *N* strings per length 12*×* 64, 12 *×*256, 12 *×*512, 16*×* 128, 16 *×*256, 16 *×*512, 16*×* 1024, 20 *×*512; the square root, which uses its whole budget of time, at four of them. Arithmetic modulo five is run three times: with questions, without (no question may be asked), and with the learner made to guess. Ten seeds.

##### Result

Table 17 and Figure 3e. With questions, arithmetic modulo five is exact for every seed from 5,843 strings, with at most one question (median 0); from 3,795 strings, nine seeds of ten are exact, after a median of 15 questions (without questions, none); at that experience the tenth seed reports that the program of the shorter strings does not answer the longest. Without questions arithmetic modulo five is exact for every seed from 11,135 strings; where it is not, the learner announces nothing, and no run of it announces a false program. Made to guess, it announces one false program (Appendix D.3.1). Bracketed arithmetic is exact for every seed at every experience, without a question. Equations are exact for every seed at every experience; the learner asks at most five questions for them, and only at 12*×* 64. The square root is announced at no experience.

**Table 17.**
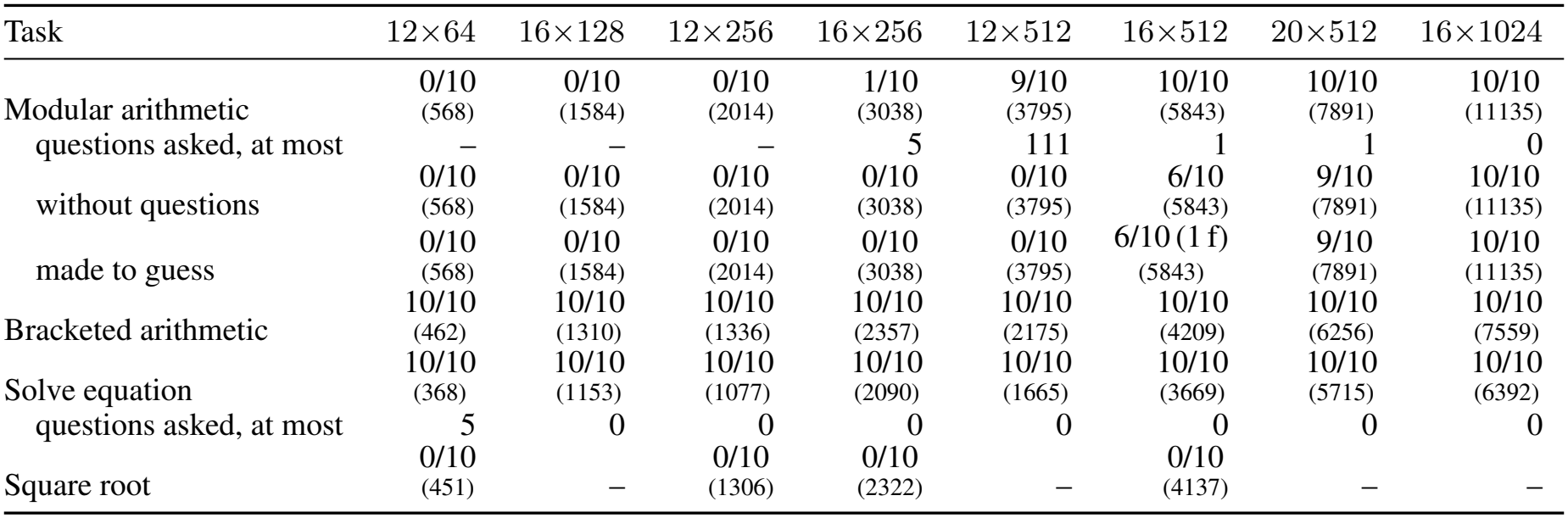
More experience: seeds exact of those run (f: runs that announced a false program; below, the median number of strings), for each experience *N×* strings per length, in the order of the number of strings. For arithmetic modulo five and for equations, also the most questions asked by a seed whose program is exact; for arithmetic modulo five, the same runs when no question may be asked and when the learner is made to guess.

| Task | 12×64 | 16×128 | 12×256 | 16×256 | 12×512 | 16×512 | 20×512 | 16×1024 |
|---|---|---|---|---|---|---|---|---|
| Modular arithmetic | 0/10<br>(568) | 0/10<br>(1584) | 0/10<br>(2014) | 1/10<br>(3038) | 9/10<br>(3795) | 10/10<br>(5843) | 10/10<br>(7891) | 10/10<br>(11135) |
| questions asked, at most | – | – | – | 5 | 111 | 1 | 1 | 0 |
| without questions | 0/10<br>(568) | 0/10<br>(1584) | 0/10<br>(2014) | 0/10<br>(3038) | 0/10<br>(3795) | 6/10<br>(5843) | 9/10<br>(7891) | 10/10<br>(11135) |
| made to guess | 0/10<br>(568) | 0/10<br>(1584) | 0/10<br>(2014) | 0/10<br>(3038) | 0/10<br>(3795) | 6/10 (1 f)<br>(5843) | 9/10<br>(7891) | 10/10<br>(11135) |
| Bracketed arithmetic | 10/10<br>(462) | 10/10<br>(1310) | 10/10<br>(1336) | 10/10<br>(2357) | 10/10<br>(2175) | 10/10<br>(4209) | 10/10<br>(6256) | 10/10<br>(7559) |
| Solve equation | 10/10<br>(368) | 10/10<br>(1153) | 10/10<br>(1077) | 10/10<br>(2090) | 10/10<br>(1665) | 10/10<br>(3669) | 10/10<br>(5715) | 10/10<br>(6392) |
| questions asked, at most | 5 | 0 | 0 | 0 | 0 | 0 | 0 | 0 |
| Square root | 0/10<br>(451) | – | 0/10<br>(1306) | 0/10<br>(2322) | – | 0/10<br>(4137) | – | – |

###### Reading

A table of 50 states is learned, and it is the most expensive thing in the paper: several thousand strings. What counts is the number of strings, whatever their length: in the order of the number of strings the table appears abruptly, and a learner that may ask needs fewer of them. The cost is that of a flat table, in which experience has to distinguish every state from every other, and most of the cost is spent on the last few entries. With half the strings the table is complete except for a handful of moments, each equivalent to several states that are distinguished, and then three things can be done. The learner made to guess takes the best supported state; such a guess is often wrong, and the longest strings catch most of the wrong ones, though not all. The learner that may not ask leaves the moments undecided, and announces nothing, because the longest strings pass through them. The learner that may ask puts one question for each moment. A learner that found separately the two parts of which each state is made, the sum so far and the product in progress, would need far fewer strings; such a learner is an open problem.

*Questions come last*. More experience does not make the learner ask more. At every larger experience the program by trial leaves nothing undecided on what was heard, and the learner asks nothing for equations; in seventeen of these 70 runs it also holds a table of inner speech that would answer an equation without trial and leaves moments undecided, and it announces the program that leaves nothing undecided. A question is put only where the learner’s own ways have left no program that leaves nothing undecided (Appendix C.3): for the table of 50 states, which no other way reaches, and, at the standard experience, for the seeds whose forecast of equations has an undecided moment.

#### D.6. Experiment 5: withheld tasks and the constants

##### Why

A learner designed while looking at fifteen tasks may be fitted to them, in its steps or in its constants. Fifteen further tasks were fixed before the learner was written and were used in no decision about it, and each constant is moved away from its value.

**Design** (E4; make e4; dm/tasks extra.py). The fifteen withheld tasks of Table 11: rotation by one; last symbol first; swapping halves, and reversing each half; repeating each symbol; interleaving halves; the first symbol repeated; exclusive or and conjunction of two strings; increment; times three; subtraction; addition in base three; a sort of three kinds in descending order; and majority. The grid of Experiment 1, ten seeds, no library and no questions: 1,200 runs. The constants are varied over all thirty tasks (Appendix D.6.1).

##### Result

Table 18 and Figure 3a. From the standard experience all fifteen tasks are exact for every seed, all by programs in step, each learned in at most 1.6 seconds. Over all 1,200 runs, none announced a false program, and none a program that abstains.

**Table 18.**
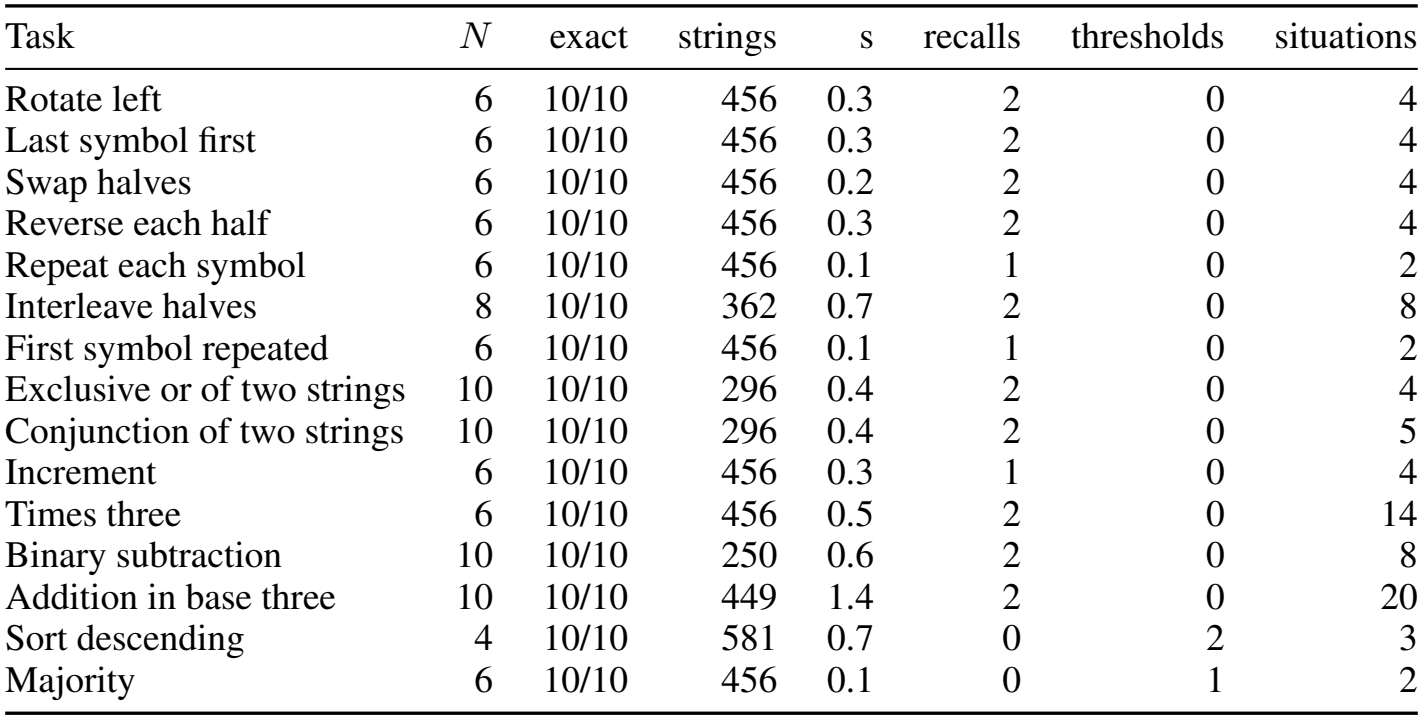
Withheld tasks. *N* : the smallest of the grid from which every seed is exact. At the standard experience: seeds exact, strings (median), seconds to learn (median), and the size of the program that most seeds write: recalls, thresholds, and situations of the table.

##### Reading

The same comparisons find tasks of new kinds. The exclusive or of two strings is the case of Theorem 1: each digit alone is independent of the answer, and the first difference of a minimal pair shows that the answer depends on it. The conjunction of two strings is the case that a test of credit by “always or never” would lose, since each digit matters only where the other is one; the first difference finds both recalls. Times three needs a second recall, of the digit before, beside the carry that the last act holds. Majority is one threshold, the side of “ones less zeros”.

##### D.6.1. Changing the constants

###### Why

The learner has four constants (Appendix C.5): the *margin*, the bits by which whatever is kept must explain more than it costs to state; *luck*, the standard deviations by which a relation must account for more dependences than chance; *idle*, the share of returns below which a kind of record may be filtered out of a recall; and *major*, the share of all differences above which a recall is part of every program. If the results depended on them delicately, the constants would be fitted to the tasks.

**Design** (E8; make e8). Each constant is set below and above its value, one at a time, the others unchanged: the margin to 10 and 40 bits (its value is 20), luck to 3 and 5 standard deviations (4), idle to 0.05 and 0.2 (0.1), and major to 0.1 and 0.25 (0.15). The thirty tasks, the standard experience, ten seeds: 2,400 runs.

###### Result

Table 19. Under each of the eight settings the learner solves the same 28 tasks as with the usual constants (Experiment 3), and none of the 2,400 runs announced a false program.

**Table 19.** The constants: tasks (of thirty) solved, runs that announced a false program, and runs whose program abstained on some test string, with one constant changed.

| Setting | solved | false | abstains |
|---|---|---|---|
| margin = 10 (default 20) | 28 | 0 | 0 |
| margin = 40 (default 20) | 28 | 0 | 0 |
| luck = 3 (default 4) | 28 | 0 | 0 |
| luck = 5 (default 4) | 28 | 0 | 0 |
| idle = 0.05 (default 0.1) | 28 | 0 | 0 |
| idle = 0.2 (default 0.1) | 28 | 0 | 0 |
| major = 0.1 (default 0.15) | 28 | 0 | 0 |
| major = 0.25 (default 0.15) | 28 | 0 | 0 |

###### Reading

The constants are levels of evidence, and within these ranges they change the outcome of no task: what the learner finds is fixed by the comparisons, and the constants say only how much evidence a part must have before it is kept.

#### D.7. Experiment 6: experience made by the machine

##### Why

For some tasks the learner finds no program in step. The first digit of a sum said highest digit first depends on every carry, and no program in step that says only its answers gives it on all inputs (Proposition 7); for multiplication the learner finds no program in step either. By Theorem 5, a learner with programs already learned can reach such tasks by learning from experience that it makes itself: running a program it knows (composition), trying each candidate answer (trial), and comparing an answer with the answer to the question one symbol shorter (recursion).

**Design** (E5; make e5). *Composition* (tag think): the sum of two numbers, the sum in base three, the increment and the product by three, each said highest digit first (Table 11), with the program of the same task said lowest digit first in the library; the standard experience, ten seeds. The learner hears its own answer, said lowest digit first, as further input, and learns in step on what it heard and said (Appendix C.4). *Trial*: equations, in Experiment 1. *Recursion*: multiplication at every *N* of the grid of Experiment 1, with questions and without; without questions also with 256 and 1,024 strings per length at *N* = 12. *Cost*: the moments that the learned multiplication lives through to answer (the symbols heard and said by the programs it calls), against the length of the question, at lengths doubling from 50 to 3,200.

##### Result

*Composition*: all four tasks are exact for every seed (one of them, the increment, in step without composition). *Trial*: equations are exact for every seed from *N* = 12 (Table 12). The learner finds which symbol stands for the answer, predicts the last symbol of a true statement with a table of 15 states of inner speech and a counter of depth for the brackets, and tries each of the five values in the place of the unknown. *Recursion*: Table 20. From the standard experience with questions, 10/10 seeds are exact, after a median of 990 questions. The answers are exact at every length tried, up to 3,200 symbols, and the moments to answer grow as the power 2.02 of the length (Figure 3f; the seconds on one core grow as the power 1.5, the overhead of the interpreter being linear). Without questions no seed is exact at any experience: of the strings of a standard experience, a fifth (19%) have their neighbour by deletion in memory.

**Table 20.** Multiplication by recursion: seeds exact when the learner may ask (it asks for the answers along the way of its questions; median number of questions) and when it may not; –: not run.

| *N*, strings per length | with questions | questions | without questions |
|---|---|---|---|
| 4, 64 | 0/10 | 4 | 0/10 |
| 6, 64 | 0/10 | 53 | 0/10 |
| 8, 64 | 10/10 | 189 | 0/10 |
| 10, 64 | 10/10 | 495 | 0/10 |
| 12, 64 | 10/10 | 990 | 0/10 |
| 12, 256 | – | – | 0/10 |
| 12, 1024 | – | – | 0/10 |
| 14, 64 | 10/10 | 1688 | 0/10 |
| 16, 64 | 10/10 | 2598 | 0/10 |
| 20, 64 | 10/10 | 5085 | 0/10 |

##### Reading

What the learner says is stored like what it hears, and comparing it writes programs that no comparison of what was heard alone writes. The program that the learner finds for the differences is one row of the schoolbook procedure, and the program it announces adds the rows: a procedure of quadratic time built from programs in step, by the method of Babbage’s engine. It needs the answers along the way of its questions, and passive experience seldom holds them, too seldom to find the program of the differences. A learner that may ask gets them. Recursion and the undecided moment are the two places in the paper where questions are needed, and in both the question is a neighbour made on purpose: by deletion in the one, by substitution of a beginning in the other.

#### D.8. Experiment 7: the engine in neural circuits

##### Why

Section 5 places the learner in neural circuits, and the account should hold at that level (Section 5.5). Five claims are tested in simulations of the rings of Eq. (4) and of threshold units: that the wells of a ring make gains act as whole numbers; that a learned program runs as a circuit; that a ring learns the gains of a counter by the local rule of Theorem 6, from dependences that nobody has sorted; that a sweep compares two episodes; and that consolidation is a circuit that searches for nothing.

**The rings** (E9; code/neural/, experiments/e9 neural.py). A counter is held by rings of rate neurons, the continuous attractor networks with adaptation of Eq. (4) (Section 5.5), in the form used in the first paper (He & Wu, 2026a; Mi et al., 2014; Chu et al., 2024). The neuron w ith preferred phase *θ ∈* (*−π, π*] has synaptic input *U* (*θ, t*), adaptation current *V* (*θ, t*) and firing rate *r*(*θ, t*), and 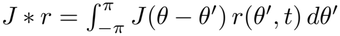. Here *τ* is the time constant of the neurons, the unit of time below. *ρ* is the density of neurons on the ring. *J* is the recurrent kernel, a Gaussian of the distance on the ring whose width is two neurons, so that the network can hold a bump of activity at any phase. [*U* ]_+_ = max(*U*, 0), and *k* is the strength of divisive inhibition, set to half the largest value at which a bump persists. The weak periodic input *ϵ* cos(*pθ*) makes *p* wells, one for each state of a ring of period *p*, and *ϵ* is their depth. *V* is an adaptation current of strength *m* and time constant *τ*_*v*_. *I*(*θ, t*) is the input from outside the ring. The reading of a ring is the well nearest to the phase of the population vector of its bump. An event turns a ring by a pulse: for a time proportional to the gain, 4*τ* for each well, the recurrent input receives in addition an asymmetric part proportional to *J*^*′*^ *\*r*, where *J*^*′*^ is the derivative of *J*, and this carries the bump forward or back by the sign of the gain. When a counter is run the wells then catch the bump, within 10*τ*, so that it moves by the whole number nearest to the gain. A reset is a brief strong input at the well of zero, and noise of strength *σ* enters *I* in every neuron. Without adaptation (*m* = 0) the bump stays where it is put; with *m > τ/τ*_*v*_ it travels along the ring by itself (Mi et al., 2014). A counter is a set of rings of coprime periods (5, 7, 9, and 11 for the longer strings), and its reading is recovered from the wells of its rings by the Chinese remainder theorem. Except in the replay below, a ring has 16 neurons to a well, wells of depth *ϵ* = 1 and no adaptation. In the code the ring is discrete, and Eq. (4) is integrated by Euler steps of 0.1*τ* .

##### Design

*(a) Staircase*. One event of gain *g* from rest, for *g* from *−*2.5 to 2.5, on a ring of period 11; and ten events in a row, for *g* from 0.5 to 1.5. *(b) Circuits*. Each program in step that the learner wrote for the benchmark from the standard experience is compiled as Lemma 2 says: for each recall, a key counter and a query counter with their gains for every kind of event; for each threshold, a counter with a latch that holds its side; rings of two states whose return to zero is the event “every second symbol” (or “every second answer”); a place cell for each pattern of wells of a key counter under which a record is stored, the latest record replacing the earlier; and the table of the program, which receives the records at the active place cells and the latched sides. The circuit is run on fresh strings of 10, 20, 40, 80 and 120 symbols and compared, symbol by symbol, with the program and with the answers due (three seeds of the experience; four strings at each of five lengths). Then, for four of the programs (reversal, the stack, addition and sorting; seed 0), it is run on strings of twenty symbols with noise of strength *σ* in every neuron. *(c) A counter learned in a ring*. The key counter of the first recall of a task, a pair of rings of periods 7 and 9 with one gain for each kind of symbol, starts from gains drawn uniformly from (*−* 0.4, 0.4): the counter of the stack (five kinds of symbol), and those of reversal and of copying. All pairs of dependences of the standard experience are used, unsorted: two answers of one experience up to four steps apart, and the two records they depend on, found by the first differences of minimal pairs. They are presented in random order, three times over. For each pair, the stretch of the experience between the two records is replayed in a sweep, in which the pulses of its events follow one another with no time to settle. The travel of the bump is compared with how far the query counter of the recall moves between the two answers (an answer turns it back for the stack and for reversal, forward for copying). The miss *µ* is felt through the firing field of the record’s place cell, as 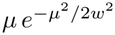 with *w* half a well, and changes the gains by the rule of Theorem 6 with *η* = 0.5. A trial succeeds if the gains, run one event at a time, come to move the counter as the gains that elimination finds from the same dependences within the first two thirds of the presentations, and keep doing so to the end. The program is then compiled with the real gains that the ring ended with in place of its whole numbers, and run on eight fresh strings of twenty symbols. In the control the miss is felt in proportion at any distance. Twenty trials of each task with the field, and twenty without. *(d) Replay*. Two episodes of twelve symbols that differ in one place are stored along one ring of twelve wells, one record to a well (8 neurons to a well, a kernel of width 3 neurons, adaptation *m* = 0.5 with *τ*_*v*_ = 5*τ*, wells of depth 0.05). After a brief push the bump is left to itself. A mismatch cell receives the records of both episodes at the active place cells. *(e) Consolidation*. The circuit of Appendix B.7, threshold units that excite the units of the values shown equivalent to theirs and a veto from every distinguished pair, on the evidence of every input of the tables of the programs in step learned for the benchmark and the withheld tasks from the standard experience (seed 0): 46 inputs of 21 programs.

##### Result

*(a)* Every gain moved the counter by the whole number nearest to it, and a gain halfway between two whole numbers by the one nearer zero; ten events in a row moved it ten wells for every gain within 0.35 of one. *(b)* Seven of the eight programs compile into rings, place cells and latches alone. The recall that does not is the second recall of the missing duplicate, whose query counter is the sum of two counters, one of which a symbol resets; it needs a counter set to where another stands (Lemma 2). On 420 strings of up to 120 symbols the circuits answered as the programs on all; with noise, on every string tried (eight of each program at each strength) up to *σ* = 1, and on none at *σ* = 1.2. *(c)* Of the 172 pairs of dependences of the stack, 26% cannot be served by the counter of the stack. From random gains the ring came to act as (+1, +1, *−* 1, +1, +1), the gains that elimination gives, in 20/20 trials, after a median of 124 dependences (Figure 4d). The gains it ended with were within 0.24 of whole numbers, and the circuit run with them answered all of 160 strings as the program does. Without the field, 0/20 trials succeeded. Over the 3 tasks of the run, 3 had their counter learned and held in every trial with the field (60/60 trials). Where every dependence is of the kind that the recall serves, as in reversal and copying, the field is not needed. *(d)* The bump travelled through the twelve wells in order, at 10 time constants a record, and the mismatch cell fired only at the record where the episodes differ (Figure 4b). *(e)* The circuit gave the classes of the learner in 46 of 46 cases, in at most 4 sweeps of spreading activity (Appendix B.7).

##### Reading

The three levels of Marr are one system. What the algebra solves, the ring learns by a rule that uses only the miss at a recalled record and the counts of events; what the ring has learned acts as whole numbers because of its wells; and the programs written by comparison run on the rings unchanged. The ring itself provides two things that the rule requires. The miss is graded although the running counter is not, because a sweep is faster than the wells. And the dependences need not be sorted by recall, because those that a counter cannot serve miss by whole wells, and the field of a place cell does not reach so far (Eq. (9)). Without the field the rule settles between the solutions of different recalls, where elimination picks the one on which most dependences agree.

#### D.9. Experiment 7, continued: the local rule alone

##### Why

By Theorem 6, the gains that the learner finds by elimination are also the rest state of a rule that uses only the miss of a dependence and the counts of events that the counters hold.

**Design** (E6; code/flow/flow.py, run by experiments/e6 flow.py). A counter has one real gain for each kind of event. The dependences are those of reversal (two kinds of event) and of the stack of the benchmark (six: two symbols, a pop, two pushes, and the answer step), drawn from random strings of 1 to 10 symbols. At each dependence the gains change by*− η µ* Δ**N***/* |Δ**N** |^2^, where *µ* is the miss of Eq. (3) and Δ**N** the numbers of events of each kind between the record and the answer; the offset of the recall is +1. The gains start at one for every kind, the counter of time, or are drawn from a standard normal distribution; 200 runs each. Recorded: the number of dependences after which the gains, rounded, are those of the program and remain so, and the share of the dependences of strings of 41 to 500 symbols that are served.

##### Result

*Convergence*. Every run reaches the program. At *η* = 1 the median number of dependences is 1 for reversal from the counter of time and 13 from random gains, and 75 and 126 for the stack, the dependences of about 20 and 34 strings. The squared error of the gains shrinks with each dependence by the factor that the spectrum of the dependences predicts (0.895 predicted, and 0.892 to 0.896 measured, for reversal). *The step*. In a single run at *η* = 0.2, the share of the long dependences that the rounded gains serve is at most 0.006 for the first 38 dependences, and 1 from the next on. Without rounding, the gains after 100 dependences serve, on average over the runs, 91% of the dependences of the strings heard and 5% of the long ones. *No backward pass*. On 300 random experiences the correction computed from the counts equals the gradient obtained by a backward pass through the moments between the record and the answer, to 2 *×*10^*−*13^. If the turn that an event gives is made to vary with the reading of the counter, with strength 0.2 or 1, the two separate: their mean cosine is 0.99 and 0.55. *The multiplier*. With multipliers that integrate the miss and leak, as in the second paper (He & Wu, 2026b), the flow of all dependences has the frequencies of that paper’s rhythm law to ten digits. Without a leak the gains circle the solution and do not arrive, and no setting of the leak is faster than letting the multiplier equal the miss. *Noise and wrong pairs*. With noise of standard deviation *σ* in the miss, rounding gives the program in at least 99% of runs at every point of the grid with 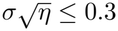 for reversal and *≤*0.2 for the stack. When a share of the dependences is paired with the wrong record, the gains settle at a least-squares point, which leaves the region of gains that round to the program at about a fifth of wrong pairs. At the smallest rate, rounding gives the program in at least 99% of runs up to a tenth of wrong pairs, and up to a fifth if the miss is clipped at one unit. *Two counters*. For a dependence between two counters, as in duplication, the flow conserves the sum of the two gains, and the solution it reaches has the scale that the start gave it. With wrong pairs that scale decays to zero, unless the key counter, under which records are stored, is held fixed. *Recalibration*. In the experiment of Jayakumar et al. (2019), landmarks are moved in proportion to the rat’s own motion, so that one lap of the track is *G* laps past the landmarks. The hippocampal map follows the landmarks, and when they are switched off the gain of path integration is found between one and the final *G*, and stays there. In the model the dependence is that the landmark be found again where it was stored, the miss is the slip of the counter against the landmarks, and the rule becomes *θ← θ− η*(*θ −G*). With *G* ramped away from one and the landmarks then removed, the gain moves toward *G*, remains between one and *G* in proportion to the exposure, and persists unchanged without landmarks, since without a dependence there is no force; the fields shift in the direction observed. The rate *η* is the one free constant, and nothing is fitted to the data: with *η* = 0.01, 0.03 and 0.1 per turn of the landmarks, the kept gain has slope 0.34, 0.65 and 0.91 against the final *G*.

##### Reading

Solving the dependences and following the rule give the same relations. The learner counts because counting is exact and fast and has no bias under wrong pairs; a brain may follow the rule. What matters for the theory is the content of the learning signal: a displacement between where a counter stands and where a record is found, known when the record is recalled, and credited to kinds of event by counters that have already formed the sum. The rule descends the squared miss. What it does without is the transport of an error through time, and it can do without it because each event turns a counter by an amount fixed by its kind.

### E. The brain: correspondence and predictions

This appendix holds the correspondence between the learner and the brain and what it claims (Appendices E.1 and E.2); how one requirement accounts for functions that separate theories give the hippocampus, with the predictions that separate each reading (Appendix E.3); a protocol for each of the eleven predictions (Appendix E.4); the scope of the theory beside the accounts it competes with (Appendix E.5); and the results that would refute it (Appendix E.6). Section 5 gives the argument in brief, and Appendix D.8 the simulations of the learner in neural circuits.

#### E.1. The parts of the difference engine

The learner has the four parts of the first paper’s memory: counters, records, recalls and a table (Section 2). It learns by comparing remembered experiences (Section 3). Table 21 sets each part, and each step of the comparison, beside the structure that the theory takes to carry it, with the evidence for the assignment and the prediction that tests it. The first four rows are those of the first paper (He & Wu, 2026a); the others are proposed here.

**Table 21.** The learner and the brain. Each part of the learner and each step of the comparison, the structure that the theory takes to carry it, the evidence for the assignment, and the prediction of Appendix E.4 that tests it. The first four rows are the first paper’s (He & Wu, 2026a); the others are proposed here.

| In the learner | In the brain | Evidence | Test |
|---|---|---|---|
| a counter, turned by each kind of event by its gain | a module of grid cells in entorhinal cortex, advanced by path integration | Hafting et al. (2005); Stensola et al. (2012); McNaughton et al. (2006) | P11 |
| a record: a moment stored with the readings of all counters | the hippocampal index of the cortical pattern of a moment | Teyler & DiScenna (1986); Teyler & Rudy (2007) |  |
| a recall: a coincidence of a key counter and a query counter | a place cell, which fires when the counters return to the readings at which its record was stored | O'Keefe & Dostrovsky (1971) |  |
| the table: from coincidences to acts | prefrontal cortex and striatum | Wallis et al. (2001); Yin & Knowlton (2006) |  |
| storing an experience whole, after one exposure | one-trial plasticity in CA3 and CA1 | Marr (1971); Nakazawa et al. (2003); Bittner et al. (2017) | P1 |
| keeping neighbours apart | pattern separation in the dentate gyrus | Leutgeb et al. (2007); McHugh et al. (2007); Bakker et al. (2008) | P3 |
| recalling a neighbour by its likeness to the present | pattern completion in CA3 | Nakazawa et al. (2002); Neunuebel & Knierim (2014) | P2 |
| comparing the neighbour with the present | CA1, which receives the recalled pattern from CA3 and the present one from entorhinal cortex by the direct path | Lisman & Otmakhova (2001); Colgin et al. (2009); Duncan et al. (2012) | P2, P3 |
| writing a program needs the comparison, and running it does not | the dentate gyrus is needed to encode and not to retrieve; habits and overtrained responses run without the hippocampus | Lee & Kesner (2004); Kheirbek et al. (2013); Packard & McGaugh (1996); Knowlton et al. (1996) | P3 |
| a sweep: two moments in one cycle, and the graded miss of a dependence | theta sequences; awake and sleeping replay | Skaggs et al. (1996); Foster & Wilson (2006; 2007); Karlsson & Frank (2009) | P2, P5 |
| a well: the whole number of a gain | discrete states of a ring attractor; predicted |  | P5 |
| a dependence kept as a relation between two counters | object-vector, landmark-vector, trace, goal-vector and task-lag cells | Høydal et al. (2019); Deshmukh & Knierim (2013); Poulter et al. (2021); Sarel et al. (2017); El-Gaby et al. (2024) | P4 |
| the miss of a dependence corrects a gain | recalibration of the gain of path integration, partial and lasting, by landmarks moved in proportion to the animal's motion, or by optic flow alone | Jayakumar et al. (2019); Madhav et al. (2024) | P5 |
| a counter for each kind of event | cells that count laps, elapsed time, accumulated evidence and the steps of a task | Sun et al. (2020); Pastalkova et al. (2008); Nieh et al. (2021); El-Gaby et al. (2024) | P5, P8 |
| a reset to zero | cells that mark the boundaries of events; correction of path integration at a boundary | Zheng et al. (2022); Hardcastle et al. (2015) |  |
| a threshold: the side of zero on which a counter stands | predicted |  | P8 |
| the state carried by the last act | splitter cells; sequences with a shared middle distinguished; codes made orthogonal, or pushed apart, where the futures differ | Wood et al. (2000); Frank et al. (2000); Agster et al. (2002); Sun et al. (2025); Chanales et al. (2017) | P7 |
| consolidation merges what was shown equivalent and never distinguished | inference and integration after sleep | Ellenbogen et al. (2007); Lewis & Durrant (2011) | P6 |
| an answer is an event | corollary discharge; inner speech | Crapse & Sommer (2008); Vygotsky (1986) | P10 |
| questions that make minimal pairs | exploration that changes one thing; variation sets in speech to children | Cook et al. (2011); Küntay & Slobin (1996); Waterfall et al. (2010) |  |
| an undecided moment: equivalent to two states that are distinguished | alternation between two maps or two futures, one per theta cycle; vicarious trial and error; more exploration when evidence is confounded | Jezek et al. (2011); Kay et al. (2020); Redish (2016); Schulz & Bonawitz (2007) | P7 |

#### E.2. What the correspondence claims

##### One operation

The assignment of each step to a subfield follows the standard account of that subfield: storage in one trial and completion from a part (Marr, 1971), the index (Teyler & DiScenna, 1986), separation in the dentate gyrus and completion in CA3 (McNaughton & Morris, 1987; Treves & Rolls, 1994; Rolls, 2013), and the meeting in CA1 of the pattern recalled by CA3 with the present one brought by the direct entorhinal path (Lisman & Otmakhova, 2001; Hasselmo et al., 2002). What the theory adds is the reading of the sequence as one operation with one purpose. A learner that assigns credit by minimal pairs needs each experience stored whole after one exposure, since the experience that will later form a minimal pair with it cannot be foreseen; it needs similar experiences stored apart, since the experiences worth comparing are nearly the same; it needs a neighbour recalled by its likeness to the present; and it needs the two compared (Section 5.1). The comparison is thus a function of the circuit as a whole, and what it yields is the moment on which an answer depended.

##### The signal of a minimal pair

The theory needs a signal that compares the present with a recalled experience like it, and such signals have been recorded. The human left hippocampus signals novelty “only in conditions where one contextual component was novel and the other repeated” (Kumaran & Maguire, 2007a); human CA1 responds more as more is changed between a recalled scene and the one in view (Duncan et al., 2012), and the hippocampal mismatch signal is largest for outcomes close to the prediction without being identical with it (Long et al., 2016); CA1 cells fire when a familiar target is first found moved (Fyhn et al., 2002); and rats with hippocampal lesions fail to notice familiar cues in a new pairing (Honey et al., 1998). In each of these the present is compared with one remembered experience. The comparison of two remembered experiences with each other is a prediction (P2; Appendix E.5).

##### How a difference may be carried

CA1 can carry the result of a comparison in three ways, and the theory needs one of them. It can carry it in its rate, as the mismatch responses of human CA1 grow with the number of changes (Duncan et al., 2012). It can carry it in which cells fire, since the plateau potentials that create a new place field in CA1 need input from entorhinal cortex and from CA3 together (Bittner et al., 2015). And it can carry it in when they fire within the theta cycle, since the input from CA3 and the input from entorhinal cortex dominate CA1 at different phases and in different bands of gamma (Hasselmo et al., 2002; Colgin et al., 2009), and inhibition locked to one phase or the other improves encoding or retrieval separately (Siegle & Wilson, 2014). The simulation of Section 5.2 uses a rate: a cell that receives the records of both replayed episodes fires only where they differ (Figure 4b; Appendix D.8).

##### Exact when it runs, graded when it replays

A counter is a ring attractor whose bump of activity is held in one of a few wells. When the counter runs, events come more slowly than the wells act: a pulse of gain *g* carries the bump *g* wells, the wells catch it, and the counter moves by the whole number nearest to *g*. When a stretch of experience is replayed in a sweep, its events follow one another faster than the wells act, the bump adds their pulses as they are, and it arrives at a graded distance from the record, the miss of Eq. (3) (Appendix B.10). The same ring is therefore exact when it runs and linear when it replays. This is why the correspondence assigns the graded miss to sweeps and the whole number to the running code, and P5 tests it: a gain that changes is graded in replay and whole in behaviour.

##### Where the table is held

The table of a program is a finite map from situations to acts. Prefrontal neurons encode abstract rules such as “same” and “different” and apply them to pictures never seen (Wallis et al., 2001); prefrontal cortex holds rules as patterns of activity that bias processing elsewhere (Miller & Cohen, 2001); and the striatum learns maps from situations to actions and keeps them as habits (Yin & Knowlton, 2006). The theory places the table in these structures because they hold such maps. Running a program needs its table, its counters and the index; writing it needs the comparison as well (Appendix E.5).

#### E.3. One requirement, many functions

Section 5.1 reads six bodies of work on the hippocampus through one requirement, credit by comparison. Appendix F.6 describes these accounts; here each reading is stated with the predictions that separate it from the account it extends.

##### Memory systems

Memory for facts and events needs the hippocampus, and the learning of skills does not (Cohen & Squire, 1980; Squire, 1992). In the theory facts and events are records in the index, which a comparison needs, and a skill is a table with its gains, learned by counting. Within the hippocampal circuit the line falls at the stage that keeps neighbours apart: the dentate gyrus is needed to encode a spatial or contextual memory and not to retrieve it (Lee & Kesner, 2004; Kheirbek et al., 2013), and silencing it during learning should remove learning from minimal pairs and spare rules that one cue decides (P3).

##### Configural and relational learning

Configural association theory held the hippocampus necessary for learning combinations of cues (Sutherland & Rudy, 1989), and relational theory for binding items into relations that can be used flexibly (Cohen & Eichenbaum, 1993; Eichenbaum, 2004). Rats with hippocampal damage solve discriminations that single cues decide and are impaired on transverse patterning, in which each cue is correct on half the trials on which it appears (Alvarado & Rudy, 1995). This is the case of Theorem 1, in which no statistic of single cues credits a cause. Under some conditions such rats also acquire configural discriminations, and the authors of configural theory concluded that the hippocampus speeds the learning of combinations, which cortex can learn without it (Rudy & Sutherland, 1995). The theory reads both results in the same way. A rule with joint causes can be learned without comparison, from statistics of the conjunctions themselves, by crediting cues in sets (Proposition 8); what needs the comparison is learning it in a step, at the trial that completes a minimal pair (P1, P3).

##### Complementary learning systems and schemas

In complementary learning systems the hippocampus stores episodes fast and replays them to a cortex that learns slowly (McClelland et al., 1995; Kumaran et al., 2016), and once a schema has been learned, a new association learned in one trial soon needs the hippocampus no more (Tse et al., 2007). In the theory replay pairs episodes in order to compare them, and what the comparison writes, gains and a table, runs in entorhinal and prefrontal cortex and in the striatum. A schema is a program already written, whose states consolidation has merged: a new association, stored in the index after one experience, adds an entry to its table with no comparison, and so it soon runs without the hippocampus, which is still needed to learn the next association (Tse et al., 2007). Consolidation merges what experience has shown equivalent and never distinguished, and leaves apart what merely overlaps (P6).

##### Comparators

Comparator theories place in the hippocampus a match of the expected with the present, whose mismatch signals novelty, arouses attention and admits new information to memory (Gray, 1982; Vinogradova, 2001; Lisman & Grace, 2005). In the theory the mismatch also locates a cause: where two experiences differ is what an answer depended on, and what CA1 reports is credit. Two results follow that a signal of novelty does not predict. After a surprise, CA1 cells with fields at the place of the difference are over-represented in replay (P2); and silencing CA1 delays learning at the outcome that completes a minimal pair, and not at an equally surprising outcome that completes none (P3).

##### Maps and the states of tasks

The hippocampus has been described as a map of space (O’Keefe & Nadel, 1978) and of the states of tasks (Behrens et al., 2018), and models learn such maps from the statistics of experience: by predicting future occupancy (Stachenfeld et al., 2017), by gradient descent over many environments (Whittington et al., 2020), by expectation and maximization over copies of each observation (George et al., 2021), or by inferring latent causes (Gershman et al., 2010; Gershman & Niv, 2010). In the theory the map of a task is a program’s counters and its states, and the states are found from differences: two pasts are split where a common continuation is answered differently, and merged only where experience has shown them equivalent (Theorem 3). Three predictions separate this from inference of a map: the cells that hold a dependence form when a comparison first shows it (P4); a gain that changes is graded in replay and whole in behaviour (P5); and a moment that experience leaves equivalent to two distinguished states is held undecided, whatever the practice of each (P7).

##### Replay for credit and planning

Replay has been taken to carry value back from a reward (Foster & Wilson, 2006) and to plan a path (Pfeiffer & Foster, 2013), and the two uses have been joined in replay prioritized by the expected value of each update (Mattar & Daw, 2018). In the theory it also supplies the second experience of a comparison, so that after a surprise of either sign it holds the present episode together with its neighbour of the other outcome; replay prioritized for value holds such pairs no more often than chance (P2).

#### E.4. Protocols for the predictions

Table 22 lists the eleven predictions with the result from which each is derived; P1 to P7 are those of Table 2. Each protocol gives the preparation, the manipulation and what is measured (for P4 and P8 to P11 together, as the design), the result that the theory requires, what the main alternative account predicts, and the result that would refute the theory. The predictions concern what is computed and in what order, and they hold at the pace at which an animal learns (Appendix E.5).

**Table 22.** The eleven predictions, the result of the theory from which each is derived, and the main alternative account from which each separates it.

|  | Prediction | From | Main alternative |
|---|---|---|---|
| P1 | A rule with joint causes (an exclusive or) is learned in a step once minimal pairs have shown each cause, and not from as many experiences with matched statistics and no pairs. | Th. 1 | associative, gradient and Bayesian learning |
| P2 | After a surprise of either sign, sweeps hold the present episode together with its nearest neighbour of the other outcome more often than chance, and CA1 fires where the two differ. | Th. 1 | replay prioritized for value |
| P3 | Silencing CA1 at the first outcome that completes a minimal pair, and not at an equally surprising one that completes none, delays learning; silencing the dentate gyrus during learning spares only rules of one cue. | Th. 1 | learning by prediction error |
| P4 | Vector cells form for an object that the outcome depends on, more than for an equally salient one, at the first informative comparison and before behaviour improves. | Th. 2 | fields formed by exposure |
| P5 | When reward moves from every fourth lap to every fifth, the replayed lap cycle stretches with each miss, while the running lap code and anticipatory licking switch together in one step. | Th. 6 | hidden-state inference; temporal differences |
| P6 | Sleep, more than an equal time awake, merges the codes of situations met in one context with one outcome, and not of situations that only share an outcome. | Th. 3 | schemas built from overlap |
| P7 | After a new start that leads only into the stretch two routines share, the code there alternates between their codes more than on the routines' own laps, and the first choice where they part falls at chance, however unequal their practice. | Th. 3 | latent-cause inference; clone-structured graphs |
| P8 | The end of a reproduced sequence is signalled by cells for a threshold between two counts. | Th. 3 | cells tuned to numbers alone |
| P9 | Combinations never met bring hesitation and the mismatch signal, and are not answered as the nearest familiar one. | Th. 4 | generalization by similarity |
| P10 | A task that cannot be answered in step is learned after its parts, whose answers are replayed before it is answered. | Th. 5 | learning the task directly |
| P11 | A learned procedure is exact at once up to the range of the entorhinal code, and aliased beyond it. | Th. 4 | decline with distance from the lengths trained |

##### P1. Minimal pairs against matched statistics (Theorem 1)

*Preparation*. Healthy adults, and patients whose damage is confined to the hippocampus.

*Manipulation*. Subjects learn to predict an outcome that is the exclusive or of two of five binary cues, *A* and *B*; the cues *C, D* and *E* are irrelevant. One group sees the sixteen combinations in which an even number of cues is present, and no two of these differ in one cue. The other group sees the sixteen combinations in which *E* is present exactly when one of *C* and *D* is; among these, sixteen pairs differ in one cue, each in *A* or in *B*, and the two members of every such pair have different outcomes. In both sets each cue is present on half the trials, each pair of cues occurs equally often in each of its four combinations, and the outcome is independent of every single cue and of every pair of cues other than *A* and *B* together, so that every rule of at most two cues agrees with the outcome on the same share of trials in the two sets. Each set also agrees with one rule of more cues, the exclusive or of *C, D* and *E* in the first and of all five cues in the second, and that rule is wrong on every combination its group has not seen.

*Measurement*. Accuracy trial by trial; for each subject, the trial at which accuracy steps up; and transfer to the sixteen combinations that the subject’s group has not seen.

*Required*. In the group with minimal pairs, accuracy steps up soon after the trials that complete the first pair differing in *A* and the first differing in *B*, wherever these fall in the sequence, and the rule then transfers to new combinations. In the group without minimal pairs learning is slow or by rote, and it does not transfer. Neither the step nor the transfer appears in patients, or when the two members of each pair are separated by more than memory holds.

*Alternative*. Associative, gradient and Bayesian learners that weigh rules by their agreement with the examples receive the same evidence for every rule of at most two cues from either set, and learn both gradually and at about the same rate. A learner that prefers rules of fewer cues favours the exclusive or of *A* and *B* more in the second set, where the other rule that fits has five cues, than in the first, where it has three, and may learn and transfer somewhat better there; none ties a step to the trials that complete the pairs.

*Refuted if* the two groups learn at the same rate and transfer alike, or if the step in the group with minimal pairs is unrelated to the trials that complete its pairs.

##### P2. Paired replay after a surprising outcome (Theorem 1)

*Preparation*. Rats with electrodes in CA1 and CA3 run laps of a figure-eight maze. A cue early on the lap and a cue late on it decide together, by their exclusive or, whether the lap ends in reward, so that each of the four kinds of lap has its own code on the common stem, as splitter cells give (Wood et al., 2000), and the kind of lap held in a replay event can be decoded.

*Manipulation*. The animal learns the rule, and in doing so meets outcomes that its behaviour shows it did not predict, of both signs: rewards where its running or licking showed that it expected none, and omissions where it expected reward.

*Measurement*. The kinds of lap decoded in sharp-wave ripples and in theta sequences in the seconds after each outcome, the place fields of the CA1 cells active in these events, and the animal’s next predictions.

*Required*. After a surprise of either sign, more events than chance hold the present lap together with the most similar earlier lap that had the other outcome, a lap that differs from it in one cue. CA1 cells with fields at the place of the differing cue are over-represented in these events, relative to their share of replay after predicted outcomes, and they fire again at the first place at which the animal’s acts on the two laps differed. These events predict the next change of behaviour: the kind of lap whose outcome the pair decides is the next to be predicted correctly.

*Alternative*. Replay prioritized for the propagation of value (Mattar & Daw, 2018) replays the path that led to the outcome, backward, when the outcome makes another action better: it grows when reward increases, and falls when reward is reduced on a linear track, where no other action is better (Ambrose et al., 2016). It holds pairs of laps no more often than chance.

*Refuted if* replay after surprise is explained by the path to the reward and by the size and sign of the prediction error alone, with pairs of laps that differ in one cue at chance.

##### P3. Credit on the informative trial (Theorem 1)

*Preparation*. Mice learn the rule of P2, with optogenetic silencing of CA1 for one second at the outcome, triggered in closed loop; in other mice the granule cells of the dentate gyrus are silenced.

*Manipulation*. The order of laps is fixed in advance, so that the informative trials are known: those that complete a minimal pair, a lap that differs in one cue from an earlier lap with the other outcome. The first four laps are one of each kind, the first two differing in both cues, so that the third and the fourth each complete a pair in each cue. CA1 is silenced at the outcome of the first informative trial only; in a yoked control, at the outcome of an uninformative trial whose outcome is as surprising, such as the first lap of a kind neither of whose neighbours in one cue has yet been run. The dentate gyrus is silenced throughout learning, or only after the rule has been learned; under the same silencing a rule that one cue decides is learned as a control.

*Measurement*. The trial at which each animal’s behaviour steps up, which in single animals is often abrupt (Gallistel et al., 2004), and the rule it has learned, tested on all four kinds of lap.

*Required*. Silencing CA1 at the first informative outcome leaves the behaviour of that trial unchanged and moves the step to the next informative trial; silencing it at the equally surprising uninformative outcome costs nothing. Silencing the dentate gyrus during learning prevents learning from minimal pairs and spares the rule that one cue decides, which co-occurrence suffices to learn, since that cue alone is correlated with the outcome (compare Theorem 1); silencing it after learning spares the rule learned, since the dentate gyrus is needed to encode a memory and not to retrieve it (Lee & Kesner, 2004; Kheirbek et al., 2013).

*Alternative*. If learning accumulates over trials, as in associative and reinforcement learning whose update scales with the prediction error (Rescorla & Wagner, 1972; Sutton, 1988), a silenced trial costs in proportion to the surprise of its outcome, so the first informative trial and the yoked trial, equally surprising, cost the same. Configural accounts, in which the hippocampus is needed for rules of joint cues and not for rules of one cue (Sutherland & Rudy, 1989), predict the dentate result as well; that half tests only the claim that learning from minimal pairs needs neighbours stored apart.

*Refuted if* silencing the first informative outcome delays learning no more than silencing the equally surprising uninformative one, or if learning from minimal pairs survives the loss of pattern separation.

##### P4. Vector cells for the records an outcome depends on (Theorem 2)

*Design*. The reward at a goal depends on an object met earlier on the path, at a distance that varies from trial to trial; a second object, equally salient and met as often, is irrelevant. Cells are recorded in entorhinal cortex, hippocampus and subiculum throughout learning.

*Required*. Cells that fire at a fixed displacement from the relevant object, of the kind recorded in these regions (Høydal et al., 2019; Deshmukh & Knierim, 2013; Poulter et al., 2021), are found for the relevant object in greater number than for the other, and they appear at the trial of the first informative comparison, before choice improves. Such a cell is the recall that serves the dependence: it fires where the query counter stands where the key counter stood at the object.

*Alternative*. Object-vector cells respond to any object placed in an environment (Høydal et al., 2019); if they form with exposure alone, two objects met equally often acquire them in equal number.

*Refuted if* vector coding is the same for both objects throughout.

##### P5. Graded in replay, whole in behaviour (Theorem 6)

*Preparation*. Mice run laps of an unchanging track and are rewarded on every fourth lap, a regime in which many CA1 cells fire most on one lap of the four (Sun et al., 2020). CA1 is imaged throughout, and licking is recorded.

*Manipulation*. The reward moves to every fifth lap. In a second group, sharp-wave ripples or theta sequences are disrupted in closed loop during the transition, with the place code left intact.

*Measurement*. Trial by trial, the lap code while the animal runs, the lap cycles replayed in ripples and theta sequences, and anticipatory licking.

*Required*. The lap counter is a ring with wells, and its gain for a lap is graded. On each trial the gain moves in proportion to the miss between the lap at which the reward was expected and the lap at which it came, so that the replayed lap cycle stretches from four laps toward five, gradually, with cells keeping their order. The running counter moves by whole wells (Theorem 6): the lap code while running and anticipatory licking switch together, in one step, at the trial at which the graded gain comes nearer to its new whole value than to its old one, and thereafter they are stable against small misses. Disrupting sweeps during the transition stops both the stretch and the switch, and spares running by the old rule.

*Alternative*. Hidden-state inference reads the new schedule as a new hidden state, and predicts a remapping of the lap code, once the evidence for that state suffices, with no graded precursor in replay (Sanders et al., 2020); learning by temporal differences predicts that licking moves gradually, trial by trial (Sutton & Barto, 2018).

*Refuted if* the lap code while running changes gradually with the replayed cycle, or remaps to an unrelated code, or if the switch has no graded precursor in replay, or if licking moves as gradually as the replayed cycle.

##### P6. What consolidation merges (Theorem 3)

*Preparation*. Adults, with the patterns of hippocampal and prefrontal activity measured by functional imaging before and after a night of sleep, or after an equal time awake.

*Manipulation*. A design of acquired equivalence (Shohamy & Wagner, 2008) with three kinds of pair of situations, matched in how similar they look: pairs met in a common context that call for the same act there; pairs that call for the same act but are never met in a common context; and pairs that some context distinguishes.

*Measurement*. The similarity between the patterns of the two members of each pair, and the transfer to one member of what is then learned about the other.

*Required*. After sleep the patterns of the first kind of pair are more similar than before, and those of the other two kinds are not; the change is absent after an equal time awake, and it predicts the transfer. The second kind is the test. Its members have never been compared, so that to merge them is to guess (Section 3.4); a learner that seeks the fewest states merges them, since nothing distinguishes them. Merging what experience has not decided is the source of the false programs of the learner made to guess (Section 4.2; Appendix B.7).

*Alternative*. Accounts in which sleep builds schemas from the overlap of replayed memories (Lewis & Durrant, 2011) predict that the second kind converges as well, since its members share an outcome.

*Refuted if* sleep makes the patterns of all similar situations converge, or those of none, or makes those of the second kind converge as much as those of the first.

*Second night*. Consolidation compares again. After a first night, one situation of a merged pair is met in a new context, where no other member of its class is met and where it calls for the same act as the situations of a second class, which other contexts distinguish from the first. The theory requires that after the next sleep its pattern be less like those of the first class than it was, and no more like those of the second: it has become undecided, and it joins neither until experience distinguishes the two for it (Theorem 3). A consolidated memory becomes labile when its recall meets a prediction error (Nader et al., 2000; Sevenster et al., 2013), which is the condition under which the learner compares a merge again. The theory is refuted if what one night has merged stays merged whatever is met afterwards, or if the situation changes its class.

##### P7. An undecided moment is held open (Theorem 3)

*Preparation*. Rats learn two routines that begin at different starts, *S*_1_ and *S*_2_, run through a common corridor *C* to an event *E*, and end differently after it, at *X*_1_ and *X*_2_. The first routine is practised three times as often as the second. In *C* the hippocampal code distinguishes the two routines, as splitter cells do, and the routine can be decoded from each theta cycle.

*Manipulation*. A new start *S*_3_ is introduced and followed by *C* and by the outcome that the two routines share there, so that experience shows *S*_3_ equivalent to both routines and distinguishes it from neither; *S*_3_ may be made to look more like one start than the other. Probe trials then follow *S*_3_ by *C* and *E*, where the routines differ.

*Measurement*. On each theta cycle in *C*, the routine decoded from the part of the cycle that represents the present position and the past, and separately from the part that looks ahead; behaviour in *C* and at *E*; and the first choice after *E*.

*Baseline*. In trained rats theta sequences alternate between possible future paths from one cycle to the next, even while the animal runs (Kay et al., 2020), and sweeps alternate between left and right whatever lies ahead (Vollan et al., 2025). The test is therefore made against laps from *S*_1_ and *S*_2_, on which the part that looks ahead may alternate while the code of the present stays with its routine.

*Required*. On laps from *S*_3_ the code of the present position in *C* alternates between the codes of the two routines from one theta cycle to the next, more than on laps from *S*_1_ and *S*_2_, as it does when the cues of two rooms are exchanged (Jezek et al., 2011), although the path in *C* is the same. In *C* the animal acts without hesitation, since there the two routines call for the same acts. At *E*, where they differ, it hesitates: it pauses, its sweeps run down both continuations (Johnson & Redish, 2007; Redish, 2016), and when it is free to seek the event that would distinguish the routines, it turns to it. The sweeps come before the sampling, and where something else the animal has learned already decides the act, it does not sample at all: the learner asks only where nothing it can do by itself has given a program that leaves nothing undecided (Appendix C.3). On the first probe the animal takes the more practised routine, or the routine whose start looks more like *S*_3_, no more often than the other. One probe with its outcome ends the alternation: the code settles on the routine with which the outcome agreed, or on a new one if it agreed with neither.

*Alternative*. Latent-cause inference (Gershman et al., 2010) and clone-structured graphs (George et al., 2021) assign *S*_3_ to the more probable state, the more practised or the more similar routine, and the first choice follows it.

*Refuted if* the code settles on the more practised or the more similar routine before any informative experience and the first probe is answered accordingly, or if the alternation on laps from *S*_3_ does not exceed that on laps from *S*_1_ and *S*_2_.

##### P8. A threshold between two counts (Theorem 3)

*Design*. A monkey or a human subject sees *k* items, *k* varying from trial to trial, and then makes *k* responses, as in tasks of sequence reproduction.

*Required*. The end of responding is a threshold: a counter turned forward by each item and back by each response, which reaches zero when the responses equal the items. Cells in the hippocampus or the parietal cortex fire at the response at which the number of responses reaches the number of items, for every *k*, and they are not tuned to a number, unlike the number neurons of Nieder (2016).

*Alternative*. If the end is read from cells tuned to numbers, cells tuned to *k* fire at the end of a trial of *k* items, and no cell fires at the end for every *k*.

*Refuted if* the end of responding is signalled only by cells tuned to particular numbers.

##### P9. Not knowing is separable from error (Theorem 4)

*Design*. A procedure is learned from situations that leave out some combinations of what its recalls return, each value having been met in other combinations; the combinations left out are then presented.

*Required*. A program that meets a situation not in its table says that it does not know (Theorem 4). On these trials the subject hesitates, samples or checks, and the hippocampal mismatch signal is present; its answers are not those of the nearest familiar situation; and on familiar situations of equal difficulty none of these signs appear.

*Alternative*. Generalization by similarity answers a combination left out as it answers the nearest familiar one, without hesitation. A learner that represents graded uncertainty hesitates where its uncertainty is high, on difficult familiar situations as on new ones; on the theory hesitation follows whether a combination has been met, whatever its difficulty.

*Refuted if* unfamiliar combinations are answered as fast as familiar ones, with the errors of interpolation.

##### P10. A task that cannot be answered in step is learned after its parts (Theorem 5)

*Design*. Subjects learn a procedure *P* aloud, and then a task whose answer is a function of what *P* would say, such as a sum to be given highest digit first, which composition answers by running addition and reading its answer backward. Magnetoencephalography decodes the items of *P* ‘s answer (Liu et al., 2019).

*Required*. The second task is not learned before *P* is. In the interval before the answer the items of *P* ‘s answer are reactivated in the order in which *P* says them, and the answer follows the order in which the second program recalls them; disrupting the interval abolishes the answer.

*Alternative*. A learner that acquires the second task directly from its examples learns it as fast whether or not *P* has been learned, and does not reactivate *P* ‘s answer before answering.

*Refuted if* the second task is learned as fast without *P* .

##### P11. Exact at once on long sequences, aliased beyond the range (Theorem 4)

*Design*. A procedure learned on short sequences is tested on sequences of increasing length.

*Required*. Performance does not decline with length up to the range of the entorhinal code. Beyond it the procedure fails by confusing moments whose readings agree in every module, as the counters of the first paper do beyond their range (He & Wu, 2026a; Fiete et al., 2008).

*Alternative*. Learners that fit the lengths they were trained on decline with distance from them, as networks trained by gradient descent do on the benchmark (Delétang et al., 2023).

*Refuted if* accuracy declines smoothly with distance from the lengths trained.

#### E.5. Scope and alternative accounts

The paragraphs below fix the scope of the theory: what it leaves to other systems, which teaching signals it leaves in place, where minimal pairs come from, what has been observed of the comparison itself, and what sets the pace of learning in an animal.

##### Writing a program and running it

The theory draws a line between writing a program, which needs the comparison, and running it, which needs the counters, the index and the table. Habits and motor skills are acquired without the hippocampus. Patients with amnesia acquire mirror reading, motor skills, the repeating sequence of a serial reaction-time task, the classification of strings by an artificial grammar and probabilistic classifications (Cohen & Squire, 1980; Corkin, 1968; Reber & Squire, 1994; Knowlton et al., 1992; 1996); an overtrained response in a maze runs under the caudate when the hippocampus is inactivated (Packard & McGaugh, 1996); and a songbird learns its song with a circuit of the basal ganglia (Fee & Goldberg, 2011). Within the hippocampal circuit the line falls at the dentate gyrus, which is needed to encode a spatial or contextual memory and not to retrieve it (Lee & Kesner, 2004; Kheirbek et al., 2013). A regularity among events that are seen is learned by counting, as the rules of the second paper are (He & Wu, 2026b), and a skill is a table and a set of gains. Neither involves a dependence on an earlier moment that has to be recalled, and the theory requires that neither depend on comparison. The line falls where the theory puts it in a study of artificial grammars: amnesic patients classified new strings as well as controls, and fell behind in recognizing the strings they had studied and in judging new strings by comparison with them (Knowlton et al., 1992).

The hippocampus also does more than compare: statistical learning of a stream of syllables, which infants achieve in minutes (Saffran et al., 1996), was absent in a patient with complete loss of both hippocampi (Schapiro et al., 2014). The theory places the comparison in the hippocampal circuit and leaves its other functions as they are. Its prediction here concerns the dentate gyrus, the stage that keeps neighbours apart: silencing it removes learning from minimal pairs and spares rules that one cue decides (P3).

##### Graded teaching signals

The brain has teaching signals that are graded errors. Dopamine neurons report an error of temporal difference (Schultz et al., 1997), climbing fibres an error of movement (Ito, 2001), motor adaptation is proportional to error (Shadmehr et al., 2010), and neurons of the cortex receive instructive signals in their dendrites, specific to each neuron (Francioni et al., 2026). The miss of a dependence is a signal of the same family (Theorem 6), and the recalibration of path integration, which the correspondence counts as evidence, is the correction of a gain in proportion to an error. What the theory excludes is narrower: an error carried back through the steps of a computation. An error of temporal difference assigns credit along one trajectory, by repetition (Sutton, 1988); a comparison assigns it between two trajectories, at once, however many steps lie between the cause and the answer (Theorem 1). How dopamine assigns credit to a cause far back in time is itself under test, with a retrospective account and a prospective one set against each other (Jeong et al., 2022; Qian et al., 2025).

##### Where minimal pairs come from

Mill thought that nature seldom offers two instances alike in all but one circumstance (Mill, 1843), and learners often do without them. Infants tune to the sounds of their language from the distributions of sounds, before they know words that form minimal pairs (Werker & Tees, 1984; Maye et al., 2002), and at fourteen months they miss a switch between two words that differ in one sound (Stager & Werker, 1997); schoolchildren do not vary one thing at a time unless they are taught to (Chen & Klahr, 1999); and adults learning words keep one conjecture about a word and test it on the next occasion (Trueswell et al., 2013).

The theory names four sources of minimal pairs. Short experience is full of them: the 32 strings of five bits contain 80 (Theorem 1), and a long experience is made of short ones wherever counters are reset. An agent makes them by acting: an animal that returns to a place and does one thing differently, a child who changes one block of a tower and a scientist who varies one condition each make the minimal pair of an experience they remember; children explore in ways that separate confounded causes (Schulz & Bonawitz, 2007; Cook et al., 2011); and the learner asks (Section 4.4). Teachers and caregivers supply them, by contrasting cases (Schwartz & Bransford, 1998) and by variation sets, in which an utterance is repeated with one part changed (Küntay & Slobin, 1996; Onnis et al., 2008); about a fifth of the speech addressed to children comes in such sets (Waterfall et al., 2010), and nine-month-olds learn a contrast of sounds when its two members are paired with two different objects (Yeung & Werker, 2009). And memory makes them: replay joins pieces of experience into sequences never run (Gupta et al., 2010; Ó lafsdóttir et al., 2015), and trial, in which the learner tries each candidate answer in one place, makes minimal pairs in thought (Theorem 5). The theory thereby gives exploration a precise target: the experience that differs in one place from one that is remembered.

##### Two past episodes compared

The comparison observed so far is of the present with one recalled memory (Kumaran & Maguire, 2007a; Duncan et al., 2012). The waking comparison of the theory is of this kind: the present experience with its nearest neighbour in memory, at the moment of the outcome. The comparison of two past episodes, brought together in one sweep, is a prediction (P2). Its nearest precedents are three. Early in learning, coordinated ripples before correct choices represent both options of a choice (Singer et al., 2013), and disrupting awake ripples impairs the part of an alternation task that depends on the past (Jadhav et al., 2012); people evaluate a new choice by reactivating together, in hippocampus and medial prefrontal cortex, two memories learned apart (Barron et al., 2013); and ripples reactivate together items whose relation has been inferred (Barron et al., 2020). Much replay holds single trajectories, or a random walk on the map (Krause & Drugowitsch, 2022; Stella et al., 2019), and a theta cycle holds one of two alternatives (Jezek et al., 2011; Kay et al., 2020). None of these studies paired experiences that differ in one place and followed learning from their first difference; P2 does.

##### The pace of learning

The hippocampal codes of two corridors that differ in one cue took days to separate (Sun et al., 2025); transitive choices can be made from associative values without inference (Van Elzakker et al., 2003); and the effects of sleep on memory vary with the task (Cordi & Rasch, 2021), with no benefit for classical insight problems in one study (Schönauer et al., 2018). The learner idealizes in three respects: its memory is exact, its tasks are deterministic, and its kinds of event are given. With noise a single difference becomes a statistic over many pairs, and an animal has first to learn which events are of one kind; the number of pairings that this takes sets the pace, days where the learner takes seconds. The claims of the theory concern what is computed and in what order, and the predictions test these at any pace: a step at the trial that completes a minimal pair (P1, P3), and a graded change in replay before a whole one in behaviour (P5).

#### E.6. Results that would refute the theory

Four results would refute the theory as a whole. If a rule with joint causes were learned as fast, and with the same step, from experiences without minimal pairs as from experiences with them, the statistics of the cues being matched (P1), credit would not be a comparison. If sweeps and ripples after a surprising outcome held the present episode together with its neighbour of the other outcome no more often than chance (P2), replay would not supply the second experience of the comparison. If silencing CA1 at the first informative outcome cost no more than silencing it at an equally surprising uninformative one, or if learning from minimal pairs survived the loss of the hippocampus or of pattern separation in the dentate gyrus (P1, P3), the comparison would not be made in the hippocampus. And if the codes of dependences were unrelated to displacements between counter readings, so that learning a dependence on an earlier cue left the entorhinal and hippocampal maps unchanged (P4, P5), a dependence would not be kept as a relation between counters.

Two further results would refute parts of the theory. If sleep made the patterns of situations that share an outcome but never a context converge as much as those of situations that share both (P6), consolidation would not follow the rule of Theorem 3, which merges only what experience has shown equivalent and never distinguished. And if a moment equivalent to two distinguished states were settled by practice or by likeness before any informative experience (P7), the brain would guess where the learner does not.

### F. Related work

The difference engine answers two questions that are usually asked apart: how a learner arrives at a procedure that holds beyond the examples it was given, and how it finds the earlier moment on which an answer depended. Machine learning answers the first with networks trained by gradient descent, with the search of program synthesis, and with the methods of grammatical inference (Appendices F.1 and F.2). It answers the second with backpropagation, with the rules proposed to replace it in brains, and with the credit assignment of reinforcement learning (Appendix F.3). Proposition 8 places the learner between these traditions: a learner that keeps no experience credits causes that act jointly in sets or by gradients, and a learner without counters carries what it has learned to other lengths by search. The learner answers both with one method, the method of difference, which comes from the logic of experimental inquiry (Appendix F.4) and which children use when they compare and explore (Appendix F.5). Theories of the hippocampus have each explained a part of what this method asks of a memory (Appendix F.6). Appendix F.7 places the paper in its series.

#### F.1. Learning procedures from data

##### Networks and the lengths they reach

Networks trained by gradient descent learn procedures from examples, and the procedures hold beyond the lengths trained when the network has a memory of the right structure. Delétang et al. (2023) trained recurrent networks, LSTMs, Transformers, and recurrent networks given a stack or a tape on the fifteen tasks of Table 1, which are graded by the Chomsky hierarchy, on strings of up to 40 symbols, and scored them on strings of 41 to 500. Recurrent networks and Transformers failed beyond the regular tasks, LSTMs added counting, and the networks with a stack or a tape reached the higher levels; randomized positional encodings raised the accuracy of Transformers on the same tasks by 12.0% on average (Ruoss et al., 2023). Analyses of what finite networks can represent agree. With finite precision, LSTMs and networks of rectified linear units can count and gated recurrent units cannot (Weiss et al., 2018b); LSTMs recognize balanced brackets of one kind, as counter machines do, and fail on two kinds, which recurrent networks with a stack learn (Suzgun et al., 2019a;b); self-attention represents parity and nested brackets with a confidence that falls toward chance as the length grows (Hahn, 2020); Transformers trained on formal languages learn some counter languages and fail on some regular ones (Bhattamishra et al., 2020); Transformers of logarithmic precision lie in uniform TC^0^ (Merrill & Sabharwal, 2023); and recurrent networks with rational weights are universal (Siegelmann & Sontag, 1995). These results concern what a network can hold and what gradient descent finds in it. The first paper of this series showed what the memory of Eq. (1) can run (He & Wu, 2026a); this paper asks what experience writes in it.

##### What a network reads at each step

Length generalization has been obtained by changing what a network reads at each step: positional codes suited to the task (Kazemnejad et al., 2023; McLeish et al., 2024; Cho et al., 2024), scratchpads that write out the intermediate steps (Nye et al., 2021; Anil et al., 2022), formats of the data (Lee et al., 2024), and tasks that a short program in a language matched to the Transformer solves at every length (Zhou et al., 2024); products of many digits remain hard (Jelassi et al., 2023; Dziri et al., 2023). The difference engine shares the diagnosis that extrapolation depends on what is read at each step. The learner finds its recalls by solving the dependences that minimal pairs show, and its table reads coincidences alone, which mention no length; this is why, within bounds on its parts, a program that is right up to a length the bounds fix is right at every length (Theorem 4). Benchmarks of compositional generalization ask the same question of combinations. Sequence-to-sequence networks fail on new combinations of commands and on longer commands in SCAN (Lake & Baroni, 2018); accuracy falls as the compounds of test and training diverge in CFQ (Keysers et al., 2020); length generalization is the productivity of the taxonomy of Hupkes et al. (2020); and meta-learning over many compositional episodes reaches systematic generalization like that of people (Lake & Baroni, 2023). These benchmarks vary one part at a time to test what has been learned, and the learner learns from the same variation: a minimal pair is the smallest difference between two compounds.

##### Networks with a structured memory

Networks given an external memory with structure learn procedures that hold beyond the lengths trained. A controller with a differentiable stack, queue, deque, tape or addressable memory learns copying, sorting, associative recall and transductions (Graves et al., 2014; 2016; Joulin & Mikolov, 2015; Grefenstette et al., 2015), and these are the networks that reach the higher levels of the benchmark (Delétang et al., 2023). LSTMs learn to execute short programs with a curriculum (Zaremba & Sutskever, 2014); the Neural GPU learns binary addition and multiplication from numbers of up to twenty bits and answers much longer ones, with errors on some highly symmetric inputs (Kaiser & Sutskever, 2016; Price et al., 2016); networks learn pointer algorithms from inputs and outputs (Kurach et al., 2016); and networks learn to compute for more steps on harder inputs (Graves, 2016; Dehghani et al., 2019; Schwarzschild et al., 2021; Bansal et al., 2022). Neural algorithmic reasoning trains networks to imitate classical algorithms, often supervising each step of an execution, and tests them on inputs larger than those trained (Reed & de Freitas, 2016; Veličković & Blundell, 2021; Ibarz et al., 2022). These works show that extrapolation needs a memory with structure, and they learn its use by gradient descent, often from execution traces. The learner computes the movements of such a memory in closed form: the stack, the queue and the counter are solutions of linear equations among counter readings (Theorem 2). From inputs and outputs alone it recovers the part of an execution trace that its recalls need, the earlier moment on which each answer depends.

##### Program synthesis

Program synthesis writes exact programs that can be read, by searching a space of programs. FlashFill finds programs over strings from a few examples with an algebra of version spaces over a language of string operations (Gulwani, 2011); sketching fills the holes of a partial program by inductive synthesis guided by counterexamples (Solar-Lezama, 2008); networks speed the search by predicting which functions a program uses, or write programs directly (Balog et al., 2017; Devlin et al., 2017); and DreamCoder grows a library of abstractions in a cycle of waking and sleeping (Ellis et al., 2021). Gulwani et al. (2017) survey the field. Recursive programs have been built from few examples by taking differences between the traces of successive examples (Summers, 1977; Kitzelmann & Schmid, 2006).

##### Inductive logic programming

Inductive logic programming searches for a logic program that, with background knowledge, entails the positive examples and none of the negative ones (Muggleton, 1991; Cropper & Dumančić, 2022), and it has been made differentiable and trained by gradient descent (Evans & Grefenstette, 2018). Shapiro’s Model Inference System found the false clause of a program that had derived a false fact by asking an oracle whether the facts used in the derivation were true, and replaced the clause (Shapiro, 1983); it assigned blame within one computation by asking questions. The difference engine assigns credit by comparing two computations, and asks its questions, which are membership queries (Appendix F.2), where experience leaves a choice.

##### Equations for programs

The learner shares with these methods the aim of an exact program that can be read, and finds it by solving equations. Each recall is credited by a comparison, its counters are solved for by elimination in time proportional to the number of dependences, and its table is counted; a list of all counters with gains up to *g* over *K* kinds of event would have (2*g* + 1)^*K*^ entries (Theorem 2). The test on the longest strings plays the part of the verifier in synthesis guided by counterexamples. The recursion of the learner takes the difference between two answers of the task itself, with an addition that the learner has learned, which is the method of Babbage’s engine (Babbage, 1864); Summers took differences between traces.

##### Program induction in cognitive science

The learning of number words, of concepts from one example and of procedures has been modelled as the induction of programs under a prior, by search in a space of programs (Piantadosi et al., 2012; Lake et al., 2015; Rule et al., 2020; Dehaene et al., 2022). In the difference engine the bias of the learner lies in the format of its memory. Within that format experience fixes the program, or leaves a part of it undecided, and the learner searches no space of programs. Chollet (2019) measures intelligence by how efficiently skills are acquired from priors and experience; the experiments report, beside accuracy, the strings, the questions and the seconds that the learner needs.

##### Rules in networks of neurons

Whether networks of neurons can hold rules that apply to every input is an old question of cognitive science. Fodor & Pylyshyn (1988) held that connectionist networks cannot account for the systematicity of thought; Marcus (2001) argued that people learn rules over variables that networks trained by gradient descent do not extend beyond their training items; and Smolensky (1988) took symbolic rules to describe, approximately and at a higher level, what networks compute. In the difference engine the learning is graded and the program is exact. A local rule moves each gain a little after each miss, the wells of the attractor that holds a counter round the gains to whole numbers (Theorem 6), and a table that receives coincidences alone applies at every length.

#### F.2. Grammatical inference and queries

The states of the learner’s programs are found by a method of grammatical inference, the field that learns automata and grammars from examples (de la Higuera, 2010), and its questions are those of learning from queries.

##### Identification

An automaton is identified from the answers to all strings up to a length that depends on its number of states (Trakhtenbrot & Barzdin’, 1973), and the states of the smallest automaton are the classes of beginnings that no continuation distinguishes (Myhill, 1957; Nerode, 1958; Rabin & Scott, 1959). Identification in the limit is possible from an informant, which labels strings as in or out of the language, and impossible from positive examples alone for classes that contain every finite language and at least one infinite one (Gold, 1967); a learner that is told the answers to the strings it hears is in the position of the first. From an incomplete sample the smallest consistent automaton is hard to find (Gold, 1978; Angluin, 1978) and hard even to approximate (Pitt & Warmuth, 1993), and learning automata from random examples alone in polynomial time would break the RSA cryptosystem (Kearns & Valiant, 1994).

##### State merging

State-merging learners build an automaton from the tree of a sample by merging states: those whose continuations up to a given length agree (Biermann & Feldman, 1972); those that share a continuation, for the reversible languages (Angluin, 1982); any pair, taken in a fixed order, whose merge leaves no negative example accepted, in the algorithm RPNI (Oncina & García, 1992); or first the pair that the most labelled continuations support (Lang et al., 1998). Exact identification can be posed as colouring the graph of distinguished states and solved by satisfiability (Heule & Verwer, 2010); L# keeps two states apart exactly when a witness separates them (Vaandrager et al., 2022); and Rivest & Schapire (1993) learn without a reset, from one continuing experiment.

##### Consolidation as state merging

Consolidation (Theorem 3) belongs to this family and is its most conservative member. It merges what experience has shown equivalent and never distinguished; it keeps apart two states that experience has never compared; and a value equivalent to two classes that are distinguished it merges with neither, leaving it undecided. Its relation of being distinguished is the apartness of L#, and it acts as a veto on every merge. From complete experience the classes are those of Myhill and Nerode. From incomplete experience the learner keeps more states than the smallest automaton would need, and where a learner that seeks the fewest states has to choose, it asks or says that it does not know (Appendix B.7). Experiment 2 shows what the choice costs: made to guess, as learners that seek the fewest states do, the learner announces false programs on experiences from which the complete learner announces none (Section 4.2).

##### Queries

A learner that chooses its examples can learn what a learner given random examples cannot. Angluin’s L* asks membership queries, for the label of a chosen string, and equivalence queries, which propose a hypothesis and return a counterexample, and with them identifies the smallest automaton in time polynomial in its size (Angluin, 1987; 1988). When equivalence queries are replaced by tests on sampled strings, the learner is probably approximately correct in the sense of Valiant (1984): a conjecture that errs on a share *ε* of the strings passes *r* sampled tests with probability at most (1*− ε*)^*r*^. This is the bound of Theorem 4, and the test on the longest strings is such a sampling. The learner’s questions are membership queries, asked where its own experience leaves something undecided: a missing minimal pair, a continuation that would distinguish two candidate states, or the question one symbol shorter that recursion needs.

##### Active learning and curiosity

Active learning more generally chooses its queries by uncertainty, by the disagreement of a committee or by the expected reduction of error, and among its settings is the synthesis of queries (Settles, 2009). Curiosity rewards a learner for the errors of its model of the world, for their reduction, or for progress in compression (Schmidhuber, 1991b;a; 2010; Pathak et al., 2017), and it has been sorted into kinds and related to the seeking of information in brains (Oudeyer & Kaplan, 2007; Gottlieb et al., 2013). The questions of the learner have a discrete target: the other half of a minimal pair, or the experience that would distinguish two states which experience has left equivalent to one moment.

##### Automata extracted from networks

Automata have been extracted from trained recurrent networks by clustering their hidden states and minimizing (Giles et al., 1992; Omlin & Giles, 1996; Jacobsson, 2005), or by running L* with the network as the oracle for membership (Weiss et al., 2018a), and the automata extracted often generalize better than the networks. The learner arrives at automata of this kind from experience directly: its inner speech is an automaton grown by merging the moments of experience on evidence (Appendix C.3).

#### F.3. Credit assignment

##### The problem

Minsky (1961) named the problem of credit assignment in his analysis of Friedberg’s program learner, which reinforced the instructions of programs that succeeded and learned little (Friedberg, 1958; Friedberg et al., 1959). “The credit for a working program”, he wrote, “can only be assigned to functional groups of instructions, e.g., subroutines.” The learner assigns credit to one recall at a time, by a comparison (Theorem 1), and builds programs from programs it has learned (Theorem 5).

##### Backpropagation and its alternatives in brains

Backpropagation through time stores a computation and carries the error back through its steps (Rumelhart et al., 1986; Werbos, 1990); real-time recurrent learning carries the sensitivities forward, at a cost that grows with the fourth power of the size of the network (Williams & Zipser, 1989). Proposals for how a brain might assign the same credit send the error back through fixed random weights (Lillicrap et al., 2016); let it arise as a difference of activity, in predictive coding (Whittington & Bogacz, 2017; 2019; Lillicrap et al., 2020) or by inferring activity before plasticity (Song et al., 2024); contrast two phases of one network, free and nudged toward the target (Scellier & Bengio, 2017), or two forward passes, on positive and on negative data (Hinton, 2022); carry the error in dendrites and in bursts of spikes (Sacramento et al., 2018; Payeur et al., 2021); or keep an eligibility trace at each synapse and broadcast a learning signal, a three-factor rule for which there is experimental support (Murray, 2019; Bellec et al., 2020; Gerstner et al., 2018). Instructive signals specific to each neuron have been observed in cortical dendrites (Francioni et al., 2026), and Richards et al. (2019) set these proposals in a common frame. For credit across time, memory and attention solve problems on which backpropagation through time struggles (Lillicrap & Santoro, 2019).

##### Credit between experiences

Each of these rules assigns credit to the parameters of one network from the activity of that network, and those for credit across time need the effect of an early event to survive, in a trace or a stored state, until the outcome. The comparison of the difference engine assigns credit between two remembered experiences and needs no such trace: the two ends are compared, and the steps between them are not consulted. The local rule that then corrects the gains (Theorem 6) has the form of a three-factor rule. During a sweep a trace at the synapses of each kind of event holds the count of its events, and the miss multiplies it. The rule descends a squared miss and carries no error back through the steps of a computation, because the counters have already summed the events between the record and the answer.

##### Credit in reinforcement learning

In reinforcement learning credit is the return carried back along one trajectory, by eligibility traces or by temporal differences between successive predictions (Sutton, 1988; Sutton & Barto, 2018). A temporal difference compares two moments of one experience; the difference engine compares two experiences at one moment. Later methods redistribute the return to the steps that predict it (Arjona-Medina et al., 2019), weight each action by the probability of the outcome in hindsight (Harutyunyan et al., 2019), condition baselines on the future (Mesnard et al., 2021), or estimate the contribution of an action to a rewarding outcome (Meulemans et al., 2023). Each estimates, with a model, what would have followed another action. A memory that holds the other experience of a minimal pair supplies that counterfactual by recall, as an experience that happened, and a question to the world produces it where memory lacks it.

##### Episodic memory for credit

Gershman & Daw (2017) proposed that episodic memory lets a learner estimate values from little data and bridge long delays between actions and rewards, and reviewed the evidence in humans and animals. Two methods carry credit by recall. Temporal value transport recalls a past memory by attention and carries the value of the present back to the moment of that memory, across delays over which discounting would lose it (Hung et al., 2019); sparse attentive backtracking passes gradients through a few past states that the present recalls, which its authors liken to being reminded (Ke et al., 2018). These are the nearest relatives of the difference engine in machine learning: they skip the steps between a cause and its effect by recalling the moment of the cause. They recall one past moment and carry value or error to it. The difference engine recalls a second experience, like the first except at the cause, and finds the cause as the place where the two differ, so that a comparison decides what receives credit.

##### Episodic control and replay

Stored episodes are replayed to train value functions (Lin, 1992; Mnih et al., 2015), with priority for large errors of temporal difference (Schaul et al., 2016), or relabelled with the goal they reached (Andrychowicz et al., 2017). They serve directly as a table of what was done and what followed, which is episodic control (Lengyel & Dayan, 2008; Blundell et al., 2016; Pritzel et al., 2017); they reinstate earlier states of working memory when a task recurs (Ritter et al., 2018); and, shaped by prediction, they guide agents toward goals (Wayne et al., 2018). In these systems an episode is an example, to be averaged with others or acted on again. In the difference engine an episode is one half of an experiment, and its use is its difference from another. One store can serve both purposes: in the account of Lengyel & Dayan (2008) the hippocampus supports episodic control, and in this one it supports comparison.

##### Minimal pairs in trained networks

Machine learning uses minimal pairs to assign credit within networks already trained. Causal mediation analysis and activation patching run a network on two inputs that differ in one respect and exchange the activity of one component between the two runs, which has located the agreement of verbs with their subjects, the recall of facts and whole circuits (Vig et al., 2020; Finlayson et al., 2021; Meng et al., 2022; Wang et al., 2023; Conmy et al., 2023); interchange interventions test whether a network implements a causal abstraction (Geiger et al., 2021; 2025); and good practice corrupts a prompt into its minimal pair and avoids adding noise (Zhang & Nanda, 2024; Heimersheim & Nanda, 2024). Delta debugging isolates the cause of a failure by narrowing the difference between an input that fails and one that does not (Zeller & Hildebrandt, 2002). Minimal pairs also evaluate networks: contrast sets perturb test examples minimally to probe decision boundaries (Gardner et al., 2020); classifiers trained on examples that people revised minimally to change the label rely less on spurious features (Kaushik et al., 2020; Teney et al., 2020); and pairs of sentences that differ in one place, one acceptable and one not, test the grammatical knowledge of language models (Linzen et al., 2016; Marvin & Linzen, 2018; Warstadt et al., 2020). In each of these the method of difference analyses a computation that exists. In the difference engine it writes the computation, and the theory proposes that brains write their circuits by the comparison with which interpretability reads the circuits of trained networks.

#### F.4. Causation and induction

The method of the learner comes from the logic of experimental inquiry, and the problem it answers is the problem of induction.

##### The canons of experimental inquiry

Bacon set beside his table of the instances in which a nature is present a table of its absence in proximity, negative instances sought among the things most akin to the positive ones, and a table of degrees (Bacon, 1620/2000). The sixth of Hume’s rules by which to judge of causes and effects is the method of difference: “the difference in the effects of two resembling objects must proceed from that particular, in which they differ”; the fifth and the seventh are agreement and concomitant variation (Hume, 1739–1740). Hume grounds the rule in expectation: “for as like causes always produce like effects, when in any instance we find our expectation to be disappointed, we must conclude that this irregularity proceeds from some difference in the causes.” Herschel stated the method of difference together with its need for experiment, and the study of residual phenomena (Herschel, 1830). Mill (1843) made five canons of these: agreement, difference, the joint method of agreement and difference, residues, and concomitant variations. He called them methods of elimination, a term he took from the theory of equations, and observed that nature seldom offers two instances alike in all but one circumstance, while an experiment produces them “almost as a matter of course, provided the process does not last a long time”.

##### The canons in the learner

The learner uses three of the canons: difference for its dependences (Theorem 1), the joint method to judge what a recall returns where nothing depends on it (Appendix B.3), and residues for what the recalls leave unexplained (Theorem 3). Its elimination is literal, of unknowns from linear equations (Theorem 2). Hume’s disappointed expectation is where the learner looks: a dependence is found where one experience disappoints the expectation that a similar one gives, at the first answer at which the two differ. A memory of episodes keeps the rare pairs that nature offers, a question is the experiment, and Mill’s proviso is Theorem 1: short experiences are dense in minimal pairs.

##### Interventions and counterfactuals

On the interventionist account *X* causes *Y* if an intervention on *X* would change *Y* (Woodward, 2003). Pearl gave interventions and counterfactuals a calculus on structural models (Pearl, 2009), in which a counterfactual is evaluated on a twin network, two copies of a model that share their background and differ in one variable (Balke & Pearl, 1994); Lewis analysed causation as counterfactual dependence at the closest world (Lewis, 1973); and Halpern and Pearl defined actual causes as dependences under some contingency (Halpern & Pearl, 2005). A minimal pair held in memory is a counterfactual that was observed, and a question that changes one symbol of a string is an intervention. The other experience is the closest world, and a dependence that one minimal pair shows, though other pairs at the same place need not, is a dependence under a contingency: in Mill’s words, “an indispensable part of the cause”.

##### Faithfulness and parity

Causal discovery from observations infers structure from conditional independences, assuming that every independence reflects the structure, the condition of faithfulness (Spirtes et al., 2000). A parity violates the condition: each operand of an exclusive or is independent of the result (Theorem 1), and one intervention, or one minimal pair, shows the dependence.

##### Grue, quus and the following of rules

Hume held that causes are known by custom, from the constant conjunction of one thing with another (Hume, 1748/2000), and finitely many examples agree with infinitely many rules. Goodman’s *grue*, which applies to things examined before a given time if they are green and to other things if they are blue, made the point for induction, and Goodman asked which predicates are projectible (Goodman, 1955). Wittgenstein made it for rules: “no course of action could be determined by a rule, because every course of action can be made out to accord with the rule” (Wittgenstein, 1953). Kripke’s *quus*, which agrees with addition for numbers below 57 and gives 5 otherwise, turned the point into a paradox about meaning (Kripke, 1982). The learner meets such rules in two steps. A rule that parts from experience at a point it names has to know how far a count has gone, and the table of the learner receives coincidences alone. Within bounds on its parts, a program of coincidences that is right up to a fixed length is right at every length the counters hold (Theorem 4): in Goodman’s terms, the format of memory fixes which predicates are projectible. The test on the longest strings then refutes the rules that part from the task within the range of experience, such as a staircase of thresholds or a gain matched to the longest operand heard.

##### Tests and hypotheses

To announce a program only after it has predicted experiences longer than those it came from is the severe test of Popper (1959), the prediction on which Whewell set most store, of “cases of a kind different from those which were contemplated in the formation of our hypothesis” (Whewell, 1840), and a novel fact in the sense of Lakatos (1970). Where the recalls leave an answer unexplained, the learner posits what would make that answer a matter of course, a threshold or a state, and solves for it, which is the form of hypothesis that Peirce (1934) described. Fodor argued that learning by testing hypotheses presupposes the hypotheses (Fodor, 1975). The learner’s class of programs is given in advance and bounded (Theorem 4), and within it the counters are found by solving equations, which needs no list of them.

#### F.5. Comparison in development

Children learn by comparing, and they look for the experiences that comparison needs.

##### Structure mapping

Analogy maps the relations of one structure onto another (Gentner, 1983), and differences are found more easily between things that are alike, since they stand out against an alignment (Gentner & Markman, 1994). People abstract a schema from two analogous cases more readily than from either alone (Gick & Holyoak, 1983); children learn relational categories when instances are compared side by side (Kotovsky & Gentner, 1996; Gentner & Namy, 1999); preschoolers extended a new word to a relational match after comparing two examples of it, and the same examples shown one at a time did not have this effect (Christie & Gentner, 2010); students who compared two worked solutions side by side gained more procedural knowledge and flexibility than students who studied them in turn (Rittle-Johnson & Star, 2007; Alfieri et al., 2013); contrasting cases prepare students to learn from instruction (Schwartz & Bransford, 1998); and a concept is learned from a near miss, an example that fails to be an instance in one respect (Winston, 1975). Gentner (2010) proposed that comparison, with the learning of symbols, drives cognitive development. A minimal pair is the limiting case of an aligned comparison: two experiences alike in everything but one place. The difference engine gives these findings a mechanism and a place in the brain: in the theory, the dentate gyrus keeps the two experiences apart, CA3 recalls one in the presence of the other, and CA1 locates their difference.

##### Contrast in language

The minimal pair is the device by which phonology finds its units: two sounds belong to different phonemes when putting one for the other in one position changes the meaning of a word (Trubetzkoy, 1939/1969), and the paper takes the term from there. Speech to children supplies minimal pairs of a larger grain: about a fifth of what caregivers say comes in variation sets, an utterance repeated with a part changed (Küntay & Slobin, 1996; Waterfall et al., 2010), and variation sets help adults learn words and phrases (Onnis et al., 2008). Infants also learn much without minimal pairs, from the distributions of sounds and from the transitional probabilities of a stream of syllables (Maye et al., 2002; Saffran et al., 1996). In the theory a regularity among events that are seen can be learned by counting, while a dependence on causes that act jointly is invisible to statistics that involve fewer than all of them and is shown by one comparison (Theorem 1). Appendix E.5 gives the evidence on where minimal pairs come from and on what infants learn without them.

##### Children as experimenters

Children explore more when the evidence they have seen leaves causes confounded (Schulz & Bonawitz, 2007), and in exploratory play they design interventions that separate the candidate causes (Cook et al., 2011); explaining outcomes that are inconsistent with what they expected directs how they test hypotheses (Legare, 2012); sixteen-month-olds use the distribution of outcomes to infer whether a failure lay in themselves or in a toy, and act on the inference (Gweon & Schulz, 2011); and eleven-month-olds learn more about objects that violated their expectations and test them in ways specific to the violation (Stahl & Feigenson, 2015). Children of seven to ten years learn to design experiments that vary one thing at a time when they are taught to, and seldom from probing questions alone (Chen & Klahr, 1999). Causal learning in children has been modelled with causal Bayes nets, which learn from interventions as well as from observations (Gopnik et al., 2004). In the theory a question is an experiment of a definite kind: an experience that differs in one place from one that is remembered, or one that would distinguish two states left equivalent to one moment. The theory predicts that children explore where their experience leaves a moment undecided, which is where the evidence they have seen is confounded.

#### F.6. Theories of the hippocampus and replay

Theories of the hippocampus have been built around different parts of what it does: storing episodes, declarative memory, relations among items, configural associations, the transfer of memories to cortex, the comparison of expectation with experience, maps of space and of tasks, and replay. In the theory of this paper these functions follow from one requirement, credit by comparison (Section 5.1), and each account explains a part of what the requirement implies. The paragraphs below take the accounts in turn. Appendix E.3 states the reading of each in the theory with the predictions that separate it from the account it extends, and Appendix E gives the correspondence of the learner with the circuit.

##### Storage, separation and completion

That the hippocampus stores patterns in one trial and completes them from parts goes back to Marr (1971); that it stores an index to cortical patterns, to Teyler & DiScenna (1986); and that the dentate gyrus separates similar patterns while the recurrent collaterals of CA3 complete them, to McNaughton & Morris (1987) and Treves & Rolls (1994), reviewed by Rolls (2013). The difference engine takes these assignments as they stand and says what their sequence is for: storing whole, keeping apart, recalling a neighbour and comparing are the steps of one operation, the assignment of credit by a minimal pair.

##### Declarative memory

In the theory of declarative memory the hippocampus and related structures are needed for memory of facts and events, and not for skills and habits, simple conditioning or priming (Squire, 1992); patients with amnesia acquire a skill of reading mirror-reversed words at the rate of controls and keep it for months (Cohen & Squire, 1980). The difference engine accounts for the line between the two. Writing a program that recalls an earlier moment needs the comparison of two stored experiences; running a program needs its counters, its table and, if it recalls records, the index; and a habit or a skill, which recalls no earlier moment, is learned by counting (Appendix E.5).

##### Relational memory

Relational theories hold that the hippocampus binds items into representations of their relations, so that memories can be compared and used flexibly in new situations (Cohen & Eichenbaum, 1993; Eichenbaum, 2004). Rats with damage to the hippocampal system learn pairs of associations and fail to infer from them the transitive and symmetric relations that intact rats infer (Bunsey & Eichenbaum, 1996; Dusek & Eichenbaum, 1997), and Eichenbaum & Cohen (2014) updated a framework that bridges the declarative and the spatial views of the hippocampus. The difference engine says which comparison is made and what it yields: a present experience is compared with a neighbour recalled by its likeness, and the result is the moment on which an answer depended. A recall of the learner is a lookup in a memory of keys and values whose keys are counter readings, compared for equality, and the hippocampus has been described as such a memory (Gershman et al., 2025).

##### Configural association

Configural association theory held that the hippocampal system is needed to represent the joint occurrence of the elements of a compound, as in negative patterning, where reward follows each of two cues alone and not the two together (Sutherland & Rudy, 1989). Animals without a hippocampal system turned out to learn such problems under some conditions, and the theory was revised: configural associations are formed in cortex, and the hippocampus enhances the units for conjunctions, making them less like the units for their elements and faster to learn (Rudy & Sutherland, 1995). Negative patterning is an exclusive or of the two cues, a rule whose causes act jointly. The difference engine gives the configural theory a reason: causes that act jointly are invisible to statistics of their parts and visible to comparison (Theorem 1). Appendix E.3 reads the revision in the same terms and gives the predictions that separate the two accounts (P1, P3).

##### Complementary learning systems

In the theory of complementary learning systems the hippocampus stores episodes fast and replays them, interleaved, to a slow cortical learner that extracts their structure (McClelland et al., 1995); its later form weights replay by relevance and adds generalization by the recurrent interaction of traces (Kumaran et al., 2016). REMERGE generalizes by letting recalled episodes activate one another recurrently, so that a probe retrieves what similar episodes have in common, and in this way accounts for transitive inference and acquired equivalence (Kumaran & McClelland, 2012). These accounts combine episodes by what they share. The difference engine uses two similar episodes for the one place where they differ, and agrees that the product of learning is held outside the hippocampus; Appendix E.3 gives its reading of replay and of schemas (P6).

##### Comparator and predictive accounts

The comparison of what is expected with what occurs has long been placed in the hippocampus: in the septo-hippocampal system, as a comparator that stops ongoing behaviour when expectation fails (Gray, 1982); in CA3, between septal and dentate input, for the habituation of orienting in rabbits (Vinogradova, 2001); in CA1, which receives the prediction of CA3 and the present that the perforant path brings (Lisman & Otmakhova, 2001), as each stage of a hierarchy of predictive coding compares its input with a prediction (Rao & Ballard, 1999); in the alternation of these inputs at opposite phases of theta, which separates encoding from retrieval (Hasselmo et al., 2002); and in a loop with the ventral tegmental area through which novelty controls the entry of information into long-term memory (Lisman & Grace, 2005). In the model of Ketz et al. (2013), the difference within one theta cycle between the pattern that CA3 recalls on CA1 and the pattern that entorhinal cortex imposes there is an error that trains the synapses from CA3 to CA1. Human imaging finds match and mismatch signals of this kind (Kumaran & Maguire, 2007b; Chen et al., 2011; Duncan et al., 2012; Long et al., 2016), and a mnemonic prediction error moves the hippocampus toward encoding (Bein et al., 2020).

##### Mismatch as credit

In these accounts a mismatch detects novelty, gates storage or trains the memory. In the difference engine it also locates the cause of an outcome. This is why the two things compared are a present experience and one past experience, recalled because it is like the present, and why the comparison is made at the outcome. The uses agree in their timing: the moment at which a mismatch calls for storage is the moment at which a minimal pair is complete.

##### Cognitive maps and the states of a task

Tolman inferred from the behaviour of rats that they learn a map of their surroundings (Tolman, 1948), O’Keefe & Nadel (1978) proposed that the hippocampus holds such a map, and the idea has been extended from space to the structure of tasks (Behrens et al., 2018). The Tolman-Eichenbaum machine learns, by gradient descent over many environments, to factor the structure of a task from its sensory content, and reproduces grid, place and object-vector cells (Whittington et al., 2020). Successor representations make place fields predict the expected future occupancy of places (Dayan, 1993; Stachenfeld et al., 2017). Inference of latent causes assigns experiences to hidden causes and infers a new cause when the old ones predict badly (Gershman et al., 2010; 2017), and reads remapping as the inference of a hidden state (Sanders et al., 2020). Clone-structured cognitive graphs learn, by expectation and maximization over copies of each observation, which past a present observation belongs to, and reproduce splitter cells (George et al., 2021; Raju et al., 2024). Each of these learns the hidden states of a task gradually, from the statistics of experience. In the difference engine the states are found from differences: two pasts are split where a common continuation is answered differently, and merged in consolidation where experience has shown them equivalent and never distinguished (Theorem 3). Appendix E.3 gives the three predictions that separate this from the inference of a map (P4, P5, P7).

##### Replay for planning and credit

Replay has been taken to consolidate memories (Wilson & McNaughton, 1994; Buzsáki, 2015), to plan (Pfeiffer & Foster, 2013; Momennejad et al., 2018), and to assign credit. Awake replay after a run reverses the order of the run, which suggested an evaluation of the sequence of the kind that reinforcement learning makes (Foster & Wilson, 2006), and the rate of reverse replay follows the change in reward (Ambrose et al., 2016). Replay prioritized where the expected gain from updating a value, weighted by the expected need for it, is largest explains when replay occurs, its forward and backward order, and its bias toward reward (Mattar & Daw, 2018); in people, backward replay of experiences other than the present one, after a reward, goes with efficient learning of the values of actions, and its authors propose it as a means of credit assignment (Liu et al., 2021). In the difference engine replay also supplies the second experience of a comparison, which is why it so often shows an experience other than the present one: a path never run, a path through space seen and never entered, or the arm not chosen (Gupta et al., 2010; Ó lafsdóttir et al., 2015; Carey et al., 2019). The accounts differ in which episodes a sweep brings together after a surprising outcome, and P2 separates them.

##### Sleep

Sleep is thought to extract the gist of memories and to integrate them into schemas (Lewis & Durrant, 2011; Stickgold & Walker, 2013), to renormalize synapses (Tononi & Cirelli, 2014), and to help in discovering what a practised task hides, as when sleep more than doubled the number of people who found a hidden shortcut (Wagner et al., 2004); in DreamCoder a phase of sleep compresses solutions into reusable abstractions (Ellis et al., 2021). Theorem 3 gives the first of these a criterion. Consolidation merges what experience has shown equivalent and never distinguished. What some experience distinguished stays distinct, and so does what was never met in a common context; and what was shown equivalent to two things that are distinguished is undecided and merged with nothing. In a circuit a distinguished pair, a prohibited pair of the second paper, silences whatever is like both of its members (Appendix B.7). P6 tests the criterion.

#### F.7. The series of three papers

Babbage designed two engines: the analytical engine, a universal computer, and the difference engine, which was to compute tables by adding differences (Babbage, 1864). The first paper of this series derived the memory of Eq. (1) from the requirement that a rule fixed on short inputs hold on inputs of every length, showed that it computes in step whatever a multitape Turing machine computes in step, and so found a counterpart of the analytical engine in the brain’s memory for space; it found its programs by gradient descent through a relaxation (He & Wu, 2026a). The second derived the mechanics by which rules act on a circuit, each rule a constraint that acts through a multiplier accumulating its violation, and learned the rules of visible variables by counting (He & Wu, 2026b).

This paper learns the programs themselves, by adding to counting the operation that a memory of episodes makes possible: comparison. Its circuits are the first paper’s rings; the flow of the gains is the second paper’s mechanics, with the multiplier of a dependence proportional to its miss when its leak is fast (Appendix B.10); and the pairs that consolidation keeps apart are the second paper’s prohibited pairs (Appendix B.7). The scaffold of grid states of Chandra et al. (2025) is a memory of the first paper’s kind, to which this paper adds how programs over it are learned. What remains is the stored program: a machine that keeps its own table in memory and recalls it as it recalls the world (Section 6).

### G. Code and reproduction

Every number, table and figure of the paper is produced by the code that accompanies it, from the results that the code stores. This appendix gives the layout of the code (Appendix G.1), the commands that reproduce the results and the way each number reaches the text (Appendix G.2), the software and the run times (Appendix G.3), and the names that the code uses for the terms of the paper (Appendix G.4). The learner, its circuits and every experiment on it need Python 3.10 or later and NumPy, and they run on processor cores; the figures need Matplotlib, the tests pytest, and the two gradient baselines PyTorch. From the root of the repository, make demo runs the learner on the main examples of the paper in about two minutes, and make test runs its 27 tests.

#### G.1. Layout and files

The repository holds the paper, the code, the results and the bibliography:

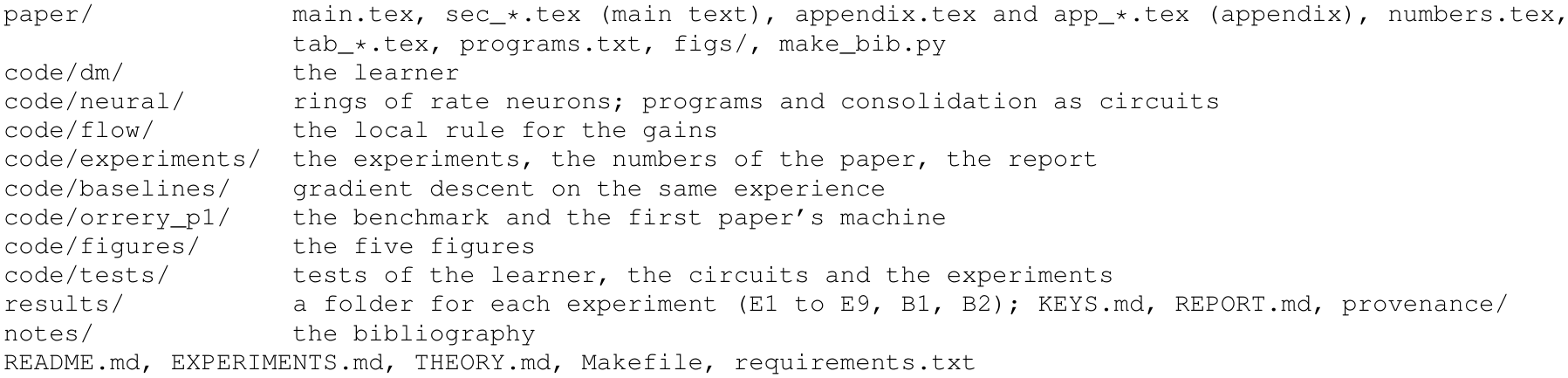

|  |  |
|---|---|
| paper/ | main.tex, sec_*.tex (main text), appendix.tex and app_*.tex (appendix), numbers.tex, tab_*.tex, programs.txt, figs/, make_bib.py |
| code/dm/ | the learner |
| code/neural/ | rings of rate neurons; programs and consolidation as circuits |
| code/flow/ | the local rule for the gains |
| code/experiments/ | the experiments, the numbers of the paper, the report |
| code/baselines/ | gradient descent on the same experience |
| code/orrery_p1/ | the benchmark and the first paper’s machine |
| code/figures/ | the five figures |
| code/tests/ | tests of the learner, the circuits and the experiments |
| results/ | a folder for each experiment (E1 to E9, B1, B2); KEYS.md, REPORT.md, provenance/ |
| notes/ | the bibliography |
| README.md, EXPERIMENTS.md, THEORY.md, Makefile, requirements.txt |  |

Table 23 gives the files of the code and what each holds.

**Table 23.** The files of the code.

|  |  |
|---|---|
| code/dm/clocks.py | counters, records and recalls; the moments of a set of experiences as arrays (Appendix C.1) |
| code/dm/machine.py | the learner, dm.learn (Algorithm 1): minimal pairs and dependences, key counters solved from pairs of dependences, queries, filters, thresholds; the order of its ways to a program and of its questions; the program in step |
| code/dm/law.py | the table counted over its features, consolidation, and the test on the longest strings |
| code/dm/inner.py | inner speech: the states that neighbours by continuation distinguish, the depth counter, undecided moments and the questions they are (Appendix C.3) |
| code/dm/closure.py | composition, trial and recursion; questions about statements (Appendix C.4) |
| code/dm/data.py, tasks_extra.py | experiences, questions made by changing one symbol, the test; the withheld tasks |
| code/orrery_p1/tasks.py | the benchmark of Delétang et al. (2023), as ported in He & Wu (2026a); with model.py, train.py and fast.py, the first paper's machine |
| code/experiments/common.py | tasks, experiences with and without minimal pairs and with questions, the test, the order in which the benchmark is learned (CURRICULUM), the fingerprint by which a program is tested once, the provenance of every result |
| code/experiments/answers.py | every task as a function of its string, for questions and for long test strings |
| code/experiments/suite.py | E1 to E5, E7 and E8; each writes its runs, a summary and its numbers |
| code/experiments/e6_flow.py, code/flow/flow.py | the local rule of Theorem 6 (E6) |
| code/experiments/e9_neural.py, code/neural/circuit.py | rings of rate neurons with adaptation; a learned program compiled into rings and place cells; gains learned in a ring; replay and the cell that compares two episodes (E9) |
| code/neural/sleep.py | consolidation as a circuit of two layers: equivalent values excite each other, and a distinguished pair vetoes (Appendix B.7) |
| code/experiments/make_numbers.py | numbers.tex, the tables tab_e⋆.tex and programs.txt, from the results; results/KEYS.md |
| code/experiments/make_report.py | results/REPORT.md, a plain report of the results |
| code/experiments/compare_runs.py | two sets of results compared run for run |
| code/baselines/gradient_baselines.py | B1: an LSTM and Transformers on the learner's experience and on fresh strings |
| code/baselines/bptt_first_paper.py | B2: the first paper's machine trained by gradient descent through time on the same experience |
| code/figures/fig1_concept.py, fig2_theory.py, fig3_results.py, fig4_brain.py | Figures 1 to 4; style.py and plotkit.py hold their common style |
| code/tests/test_smoke.py | 27 tests of the learner, the circuits and the experiments |
| code/demo.py | in about two minutes: a minimal pair, a counter solved for, addition, long multiplication from addition, an undecided moment asked about, left undecided and guessed, and programs run on a million symbols |
| Makefile, code/Makefile | the targets demo, test, pilot, full, numbers, figures, paper and zip; in code/, also one target for each experiment, e1 to e9 |
| code/make_zip.py | the whole release as one archive, with its SHA-256 |

#### G.2. Reproducing the results

The paper numbers its experiments by the claims they test, and the code names them E1 to E9, B1 and B2. Table 24 gives the correspondence, with the command that runs each experiment and the tables and figures it produces.

**Table 24.** The experiments of the paper and their names in the code. Commands are run from code/; suite stands for python3 -m experiments.suite.

| in the paper | in the code (results folder) | command | tables and figures |
|---|---|---|---|
| Experiment 1, the benchmark | E1 | suite e1 | Tables 1, 12; Fig. 3a |
| with networks | B1 | baselines.gradient_baselines | Table 13 |
| with the first paper's machine | B2 | baselines.bptt_first_paper | Table 14; Fig. 3a |
| Experiment 2, a learner that knows what it knows | E3 (census, guess, untested, no_twins_once), E7 (guess), with the runs of E1 and E4 | suite e3, suite e7 | Table 15; Fig. 3b |
| Experiment 3, what each part is for | E2 | suite e2 | Table 16; Fig. 3c |
| Experiment 4, minimal pairs and questions | E3 (no_twins, play_*f*), E7 | suite e3, suite e7 | Table 17; Fig. 3d, e |
| Experiment 5, withheld tasks and the constants | E4, E8 | suite e4, suite e8 | Tables 18, 19 |
| Experiment 6, experience made by the machine | E5 | suite e5 | Table 20; Fig. 3f |
| Section 5, the local rule | E6 | experiments.e6_flow | Appendix D.9 |
| Section 5, neural circuits | E9 | experiments.e9_neural | Fig. 4; Appendix D.8 |

##### Commands

From code/, the protocol of the paper is reproduced by

|  |  |
|---|---|
| python3 -m pytest -q tests | # 27 tests, about two minutes |
| python3 -m experiments.suite all --mode full --jobs 40 | # E1 to E5, E7, E8 (or one: e1, ...) |
| python3 -m experiments.e6_flow --mode full | # E6 |
| python3 -m experiments.e9_neural --mode full --jobs 40 | # E9 |
| python3 -m baselines.gradient_baselines --mode full --out ../results/B1 --device cuda |  |
| python3 -m baselines.gradient_baselines --mode full --out ../results/B1 --collect \<br>--tex ../paper |  |
| python3 -m baselines.bptt_first_paper --mode full --out ../results/B2 --device cuda |  |
| python3 -m baselines.bptt_first_paper --mode full --out ../results/B2 --collect \<br>--tex ../paper |  |
| python3 -m experiments.make_numbers | # numbers.tex, tab_e*.tex, programs.txt, KEYS.md |
| python3 -m experiments.make_report | # results/REPORT.md |
| python3 figures/fig1_concept.py | # and fig2_theory.py to fig5_circuits.py |
| cd ../paper && python3 make_bib.py && pdflatex main && bibtex main |  |
| pdflatex main && pdflatex main && pdflatex main |  |

From the root of the repository, make full JOBS=40 runs the experiments of the suite, E6 and E9 and then writes the numbers, the tables and the figures, and make paper writes the numbers and the figures from the stored results and compiles the paper; the baselines are run by their own commands. With --mode pilot, or make pilot, every experiment runs in a light form, with three seeds and fewer test strings. The suite also takes --tasks, --seeds and --shard i/k, which runs the *i*th of *k* parts of the jobs so that several machines can share an experiment; a last run without --shard finds every stored run and writes the summary.

##### How a number reaches the text

An experiment of the suite stores each run as one file under results/<E>/runs/, named by its mode, tag, task, experience and seed, and writes summary.<mode>.json, which keeps every run in brief, and numbers.<mode>.json; E6 and E9 write one file each, with every curve. The folders of results/ hold the summaries and numbers of every experiment of the paper, and the records of single runs of the two baselines. Every result file records the versions of Python and NumPy, the command and a hash of the learner’s source, and a stored run is reused only if it was made by the same source, so that an interrupted experiment, started again, goes on from the runs already stored. The same program is often written again, from another experience or under another setting of a constant; it is tested once, and recognized by a fingerprint of everything it is (its counters, recalls, thresholds and tables, and the programs it calls). Every number in the text is a key, written *\*res {*key}* in the source, which make numbers.py writes into paper/numbers.tex from the results; results/KEYS.md lists every key with its value, its source and its meaning, and a key without a value prints as its name in grey.

#### G.3. Software and run times

##### Software

The runs of the paper were made with Python 3.10.12 and NumPy 2.2.6 on Linux (x86-64), with forty processes on one machine for the full protocol, and the baselines B1 and B2 with PyTorch 2.7.1. The file requirements.txt asks for NumPy 1.24 or later, Matplotlib 3.7 or later for the figures, pytest 7 or later for the tests, and PyTorch 2.1 or later for the baselines. Under Python 3.13 and NumPy 2.5 the learner writes the programs of Appendix C.6 symbol for symbol as listed there. The two baselines are the only runs that need an accelerator.

##### Run times

The 15,060 runs of the suite (E1 to E5, E7 and E8) take 31 hours of one processor core to learn: E1 3.2 hours for its 1,200 runs, E2 4.1 for 3,300, E3 4.9 for 6,300, E4 0.1 for 1,200, E5 4.8 for 220, E7 10.9 for 440 and E8 2.7 for 2,400.

Testing takes longer. These runs announce 747 distinct programs, and testing them takes 232 hours of one core, of which 231 go to the 171 distinct programs that multiply by recursion: their time grows as the square of the length, and each takes about an hour and a quarter on its 14,720 test strings. A program in step or of inner speech is tested in half a minute at most, strings of a million symbols included. E6 takes about a minute of one core, and E9 took 451 seconds with forty processes. The gradient baselines took 66 hours on an accelerator for the 270 runs of B1, and 165 hours for the 210 runs of B2.

#### G.4. Names in the code

The code keeps older names for several terms of the paper. Table 25 translates them; Tables 3 to 5 in Appendix A.1 go the other way, from each term of the paper to its name in the code.

**Table 25.**
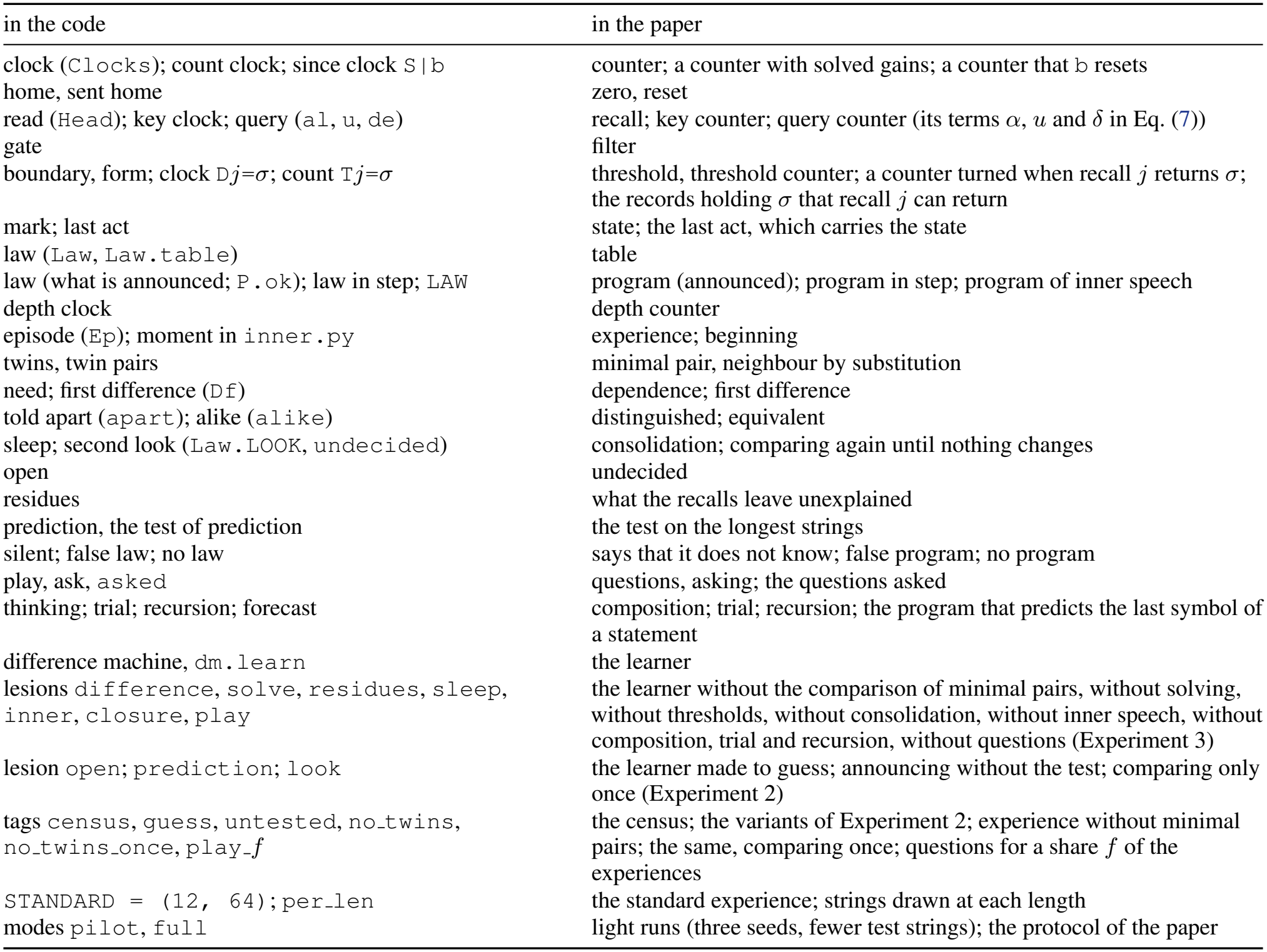
Names in the code and the terms of the paper.

| in the code | in the paper |
|---|---|
| clock (Clocks); count clock; since clock S\|b | counter; a counter with solved gains; a counter that b resets |
| home, sent home | zero, reset |
| read (Head); key clock; query (al, u, de) | recall; key counter; query counter (its terms α, u and δ in Eq. (7)) |
| gate | filter |
| boundary, form; clock Dj=σ; count Tj=σ | threshold, threshold counter; a counter turned when recall j returns σ; the records holding σ that recall j can return |
| mark; last act | state; the last act, which carries the state |
| law (Law, Law.table) | table |
| law (what is announced; P.ok); law in step; LAW | program (announced); program in step; program of inner speech |
| depth clock | depth counter |
| episode (Ep); moment in inner.py | experience; beginning |
| twins, twin pairs | minimal pair, neighbour by substitution |
| need; first difference (Df) | dependence; first difference |
| told apart (apart); alike (alike) | distinguished; equivalent |
| sleep; second look (Law.LOOK, undecided) | consolidation; comparing again until nothing changes |
| open | undecided |
| residues | what the recalls leave unexplained |
| prediction, the test of prediction | the test on the longest strings |
| silent; false law; no law | says that it does not know; false program; no program |
| play, ask, asked | questions, asking; the questions asked |
| thinking; trial; recursion; forecast | composition; trial; recursion; the program that predicts the last symbol of a statement |
| difference machine, dm.learn | the learner |
| lesions difference, solve, residues, sleep, inner, closure, play | the learner without the comparison of minimal pairs, without solving, without thresholds, without consolidation, without inner speech, without composition, trial and recursion, without questions (Experiment 3) |
| lesion open; prediction; look | the learner made to guess; announcing without the test; comparing only once (Experiment 2) |
| tags census, guess, untested, no_twins, no_twins_once, play_f | the census; the variants of Experiment 2; experience without minimal pairs; the same, comparing once; questions for a share f of the experiences |
| STANDARD = (12, 64); per_len | the standard experience; strings drawn at each length |
| modes pilot, full | light runs (three seeds, fewer test strings); the protocol of the paper |

## Notes

### Competing Interest Statement

The authors have declared no competing interest.

